# Independent somatic evolution underlies clustered neuroendocrine tumors in the human small intestine

**DOI:** 10.1101/2020.05.06.080499

**Authors:** Erik Elias, Arman Ardalan, Markus Lindberg, Susanne Reinsbach, Andreas Muth, Ola Nilsson, Yvonne Arvidsson, Erik Larsson

**Affiliations:** Endocrine and Sarcoma Surgery, Department of Surgery, Institute of Clinical Sciences, Sahlgrenska Academy at University of Gothenburg, Gothenburg, Sweden.; Department of Endocrine and Sarcoma Surgery, Surgical Clinic SU/S, Sahlgrenska University Hospital, Gothenburg, Sweden.; Department of Medical Biochemistry and Cell Biology, Institute of Biomedicine, Sahlgrenska Academy at University of Gothenburg, Gothenburg, Sweden.; Department of Laboratory medicine, Institute of Biomedicine, Sahlgrenska Cancer Center, Sahlgrenska Academy at University of Gothenburg, Gothenburg, Sweden.; Department of Pathology, Sahlgrenska University Hospital, Gothenburg, Sweden.

## Abstract

**Background:** Small intestine neuroendocrine tumor (SI-NET), the most common cancer of the small bowel, often displays a curious multifocal phenotype with several intestinal tumors centered around a regional lymph node metastasis. Although SI-NET patients often present with metastatic disease at the time of diagnosis, there is an unusual absence of somatic driver mutations explaining tumor initiation and metastatic spread. The evolutionary trajectories that underlie multifocal SI-NET lesions could provide insight into the underlying tumor biology, but this question remains unresolved. Here, we determined the complete genome sequences of 65 tumor and tissue samples from 11 patients with multifocal SI-NET, allowing for elucidation of phylogenetic relationships between tumors within individual patients. Intra-individual comparisons of whole genome sequences revealed a lack of shared somatic single-nucleotide variants and copy number alterations among the sampled intestinal lesions, supporting that they were of independent clonal origin. Furthermore, each metastasis originated from a single intestinal tumor, and in three of the patients, two independent tumors had metastasized. We conclude that primary multifocal SI-NETs generally arise from clonally independent cells, suggesting a contribution from of a cancer-priming local factor.

## Introduction

Cancer is considered to be an evolutionary process involving cycles of random oncogenic mutations, selection and clonal expansion (Yates and Campbell, 2012). In most malignant epithelial tumors (carcinomas), a single primary tumor will typically form, followed by metastases arising as cells disperse from the tumor. More rarely, carcinomas instead display *multifocality*, i.e. multiple separate tumors at the primary site, which may be due to predisposing germline variants, local mutagenic exposure, or localized spread or expansion of cancer-primed mutated clones (Curtius et al., 2018; Graham et al., 2011).

Of particular interest are small intestine neuroendocrine tumors (SI-NETs), the most common cancer of the small intestine, where ∼50% of cases display a striking multifocal phenotype that often involves 10 or more morphologically identical tumors clustered within a limited intestinal segment, commonly centered around a regional lymph node metastasis (Dasari et al., 2017; Gangi et al., 2018). SI-NET has a reported incidence of approx. 1.2 per 100,000 (Dasari et al., 2017) and often presents with distant metastases at diagnosis, thus precluding curative treatment (Choi et al., 2017; Gangi et al., 2018). However, the somatic mutational burden of SI-NET is low and there is an unusual absence of somatic driver mutations (Crona et al., 2015; Francis et al., 2013; Priestley et al., 2019). Consequently, underlying tumorigenic mechanisms are poorly understood and actionable genetic drug targets are lacking.

The evolutionary relationships that govern multiple intestinal tumors and metastases in SI-NET can give insight into the underlying biology, but earlier efforts to determine this have yielded conflicting results (Guo et al., 2000; Katona et al., 2006). A study of copy number alterations (CNA) in multifocal SI-NET primary tumors found that loss of chromosome 18 (chr18), the most frequent CNA, can affect different chromosome homologs in different samples within a patient (Zhang et al., 2020), compatible with an independent clonal origin or late loss of chr18. Alternatively, multifocal SI-NETs have been proposed to represent “drop metastases” originating from regional lymph nodes, consistent with the limited spatial distribution of the intestinal tumors (Wang et al., 2014). Adding to the challenge is the low mutational burden, which reduces the number of exonic genetic markers.

Here, we performed whole genome sequencing (WGS) on 61 separate intestinal tumors and adjacent metastases from 11 SI-NET patients, which made it possible to conclusively determine the evolutionary trajectory of multifocal SI-NET within single individuals.

## Materials and Methods

### Patients

11 patients who underwent surgery for SI-NET at Sahlgrenska University Hospital, Gothenburg, Sweden, were included in the study. The clinical characteristics of the patients are summarized in **Supplementary Table S1**. Patients were postoperatively diagnosed with well-differentiated neuroendocrine tumor grade 1-2 of the ileum (WHO 2019). Tumor grade and stage was based on one primary intestinal tumor and extent of lymph node metastases as standard clinical routine. Tumor samples were collected at surgery and were immediately snap-frozen in liquid nitrogen. A piece of each tumor was formalin-fixed and paraffin-embedded for studies by immunohistochemistry. Blood was collected and used as normal tissue in sequencing. We obtained consent from the patients and approval from the Regional Ethical Review Board in Gothenburg (Dnr:558-07, 833-18).

### Immunohistochemistry

Sections of all sampled tumors from 11 patients with SI-NET were subjected to antigen retrieval as detailed in **Supplementary Material and Methods**. Histopathological evaluation and assessment of tumor cell content was performed on hematoxylin and eosin-stained sections. Immunohistochemical staining was performed for chromogranin A, synaptophysin, serotonin and somatostatin receptor 2. These antibodies were used: anti-chromogranin A (MAB319; Chemicon; diluted 1:1000), anti-synaptophysin (SY38, M0776; Dako; diluted 1:100), anti-serotonin (H209; Dako; diluted 1:10) and anti-SSTR2 (UMB-1; Abcam; diluted 1:50). All collected tumor samples were reviewed by a board certified surgical pathologist (ON).

### DNA extraction

All tumor samples exhibited typical SI-NET morphology and expression of neuroendocrine markers. For DNA extraction, tumor samples of sufficient size and with a purity above 30% were included. DNA from fresh-frozen biopsies as well as blood from each patient was isolated using the allprep DNA/RNA Mini Kit (Qiagen) according to the manufacturer’s protocol.

### Whole genome sequencing

WGS libraries were constructed for 61 primary and metastatic tumors from the 11 patients. 11 blood normals and 4 adjacent tissue normals were additionally sequenced, for a total of 76 samples. The prepared libraries were sequenced on an Illumina Novaseq 6000 using 150 bp paired-end reads to an average depth of 36.6× (29.8-45.0).

### Mapping and somatic variant calling

Sequencing reads were mapped to the human reference genome (hg19) using BWA and somatic variant calling of matched tumor-normal specimens was performed using combined outputs from Mutect2 (GATK v4.1.4.0) and VarScan (v2.3.9). Additional details are provided in **Supplementary Material and Methods**.

### Phylogenetic analyses

Phylogenetic analyses were based on SNVs only, as indel calls contained a higher fraction problematic calls. Maximum parsimony phylogenies were reconstructed and visualized using MEGAX with default settings (Stecher et al., 2020). Bootstrapping was performed using 500 replications.

### Mutational signatures

The trinucleotide profile for each tumor was matched against known mutational signatures from COSMIC (Alexandrov et al., 2020) (v3 release) using the R package deconstructSigs (Rosenthal et al., 2016) (version 1.9.0). We used default parameters and a maximum of 4 signatures for each sample.

### Copy number analysis and homolog phasing

Copy-number alterations (CNAs) were called using XCAVATOR (Magi et al., 2017) using the RC mode and a window size of 2000 bp. The segmented copy number results were filtered to remove small segments (<25 kb) prior to visualization using IGV. Whole-chromosome alterations were phased to determine what chromosome homolog was deleted based on SNP calls reported by VarScan somatic (“germline” and “LOH” sets with a required coverage of 20). Only heterozygous SNPs were considered (variant allele frequency ranging from 0.25 to 0.75 in the blood normal). Variant allele frequencies for individual SNPs were compared between a reference sample, typically the metastatic primary tumor in each patient, and other samples of interest by means of scatter plots.

### Structural variant analyses

Structural variants (SVs) were called based on discordant read pairs and split reads using Manta (v.1.5.0) (Chen et al., 2016). Genomic coordinates of high confidence SV calls were annotated using AnnotSV (v.2.1) (Geoffroy et al., 2018). For each patient, all identified translocations were visualised in a circos plot (Krzywinski et al., 2009). A custom script was used to evaluate whether gene-gene fusions were in-frame or out-of-frame. To calculate the frames, exon and coding sequence coordinates were taken from the annotation file provided by AnnotSV.

## Results

### Evolutionary trajectories in multifocal SI-NET

We initially performed whole genome sequencing (WGS) on 6 intestinal tumors, 3 adjacent lymph node metastases, 2 peritoneal metastases and a blood normal from a patient with suspected SI-NET (Patient 1) that underwent surgery with curative intent (**Fig. 1a**). All lesions showed typical SI-NET morphology and stained positively for established diagnostic SI-NET markers (**Supplementary Fig. 1**). Samples were sequenced at an average coverage of 35.7-43.1× (**Supplementary Table S2**). This was followed by somatic mutation calling using strict filters including rigorous population variant filtering to avoid false positives in phylogenetic analyses. Resulting genome-wide single nucleotide variant (SNV) burdens varied from 353 to 1,749 (0.13-0.62 per Mb; **Supplementary Table S2**).

**Figure 1.**
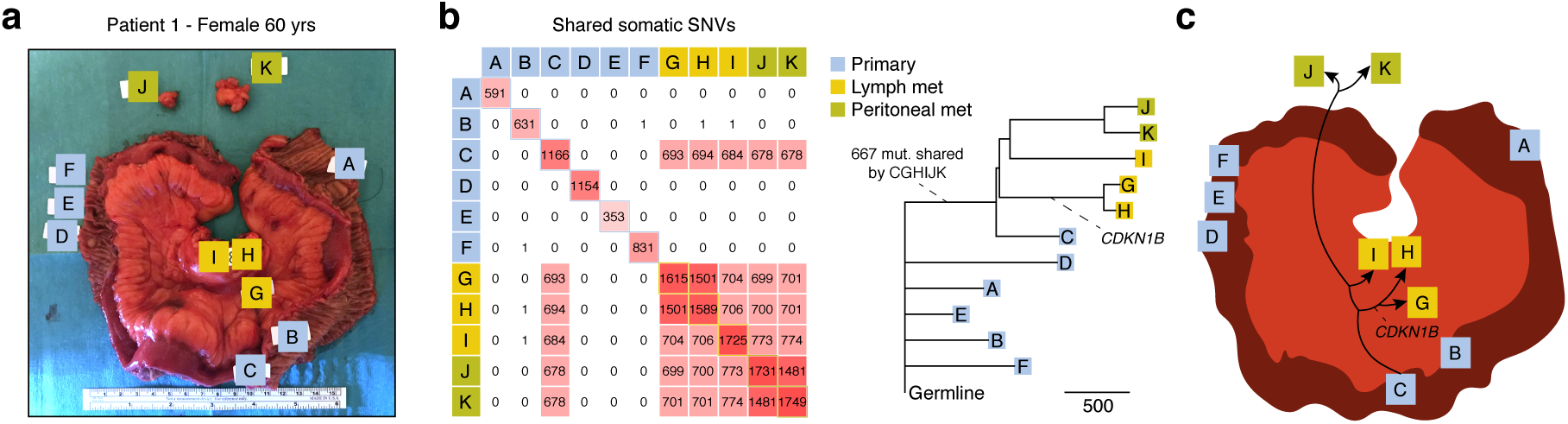
Whole genome sequencing of 11 primary tumors and metastases from a single SI-NET patient supports independent clonal evolution. **(a)** Section of the small intestine from a 60 year-old female patient harboring 6 primary tumors (A-F), 3 lymph node metastases (G-I) and 2 peritoneal metastases (J-K). **(b)** Pairwise analysis of shared somatic SNVs based on whole genome sequencing at 34.5-43.1× coverage. Primary tumor C and the 5 metastases shared a common set of 667 mutations. A maximum parsimony phylogenetic tree is shown, with the number of SNVs in each branch given by the scale marker. An indel in *CDKN1B*, a known driver events, is indicated. Bootstrap support was >= 98% for all major branches. **(c)** Proposed model, where all metastases originate from a single primary tumor, and where all primaries are unrelated in terms of somatic evolution.

We next determined pairwise shared somatic SNVs between samples. Surprisingly, a common set of 667 SNVs was shared between a single primary tumor (denoted C) and all 5 metastases (G-K), while overlapping mutations were essentially lacking between the other samples (**Fig. 1b**). The results were thus not compatible with a common clonal origin for the intestinal tumors, expected to result in hundreds to thousands of shared mutations genome-wide in the case of a late-onset cancer. Instead, phylogenetic analysis supported that the intestinal tumors had developed independently, with a single, centrally located, tumor metastasizing first to the lymph nodes and then further to the peritoneum (**Fig. 1b-c**).

To validate the initial findings, we analyzed 50 additional tumors from 10 SI-NET patients, plus matching blood normals, using WGS at 29.8-45.0× coverage (Patients 2-11; **Fig. 2; Supplementary Table S2**). The bulk of the material (Patients 3-11) consisted previously sampled tumors stored at a local biobank. Between 3 and 11 intestinal tumors and at least one lymph node metastasis was sequenced for each patient, and three cases included a liver metastasis. Macroscopically normal small intestinal mucosa samples were additionally included in four cases. Samples were selected to have sufficient tumor material and purity, and all tumor samples stained positively for SI-NET markers (**Supplementary Figs. 2-5)**. One tumor had lower-than-expected mutational burden (143 SNVs), explained by low sample purity, while others varied between 411 and 3390 SNVs (0.15-1.21 per Mb; **Supplementary Table S2**). In comparison, between 11 and 40 SNVs were called in the normal mucosa samples, where widespread clonal somatic mutations are not expected, supporting that somatic mutation calls had high specificity.

**Figure 2.**
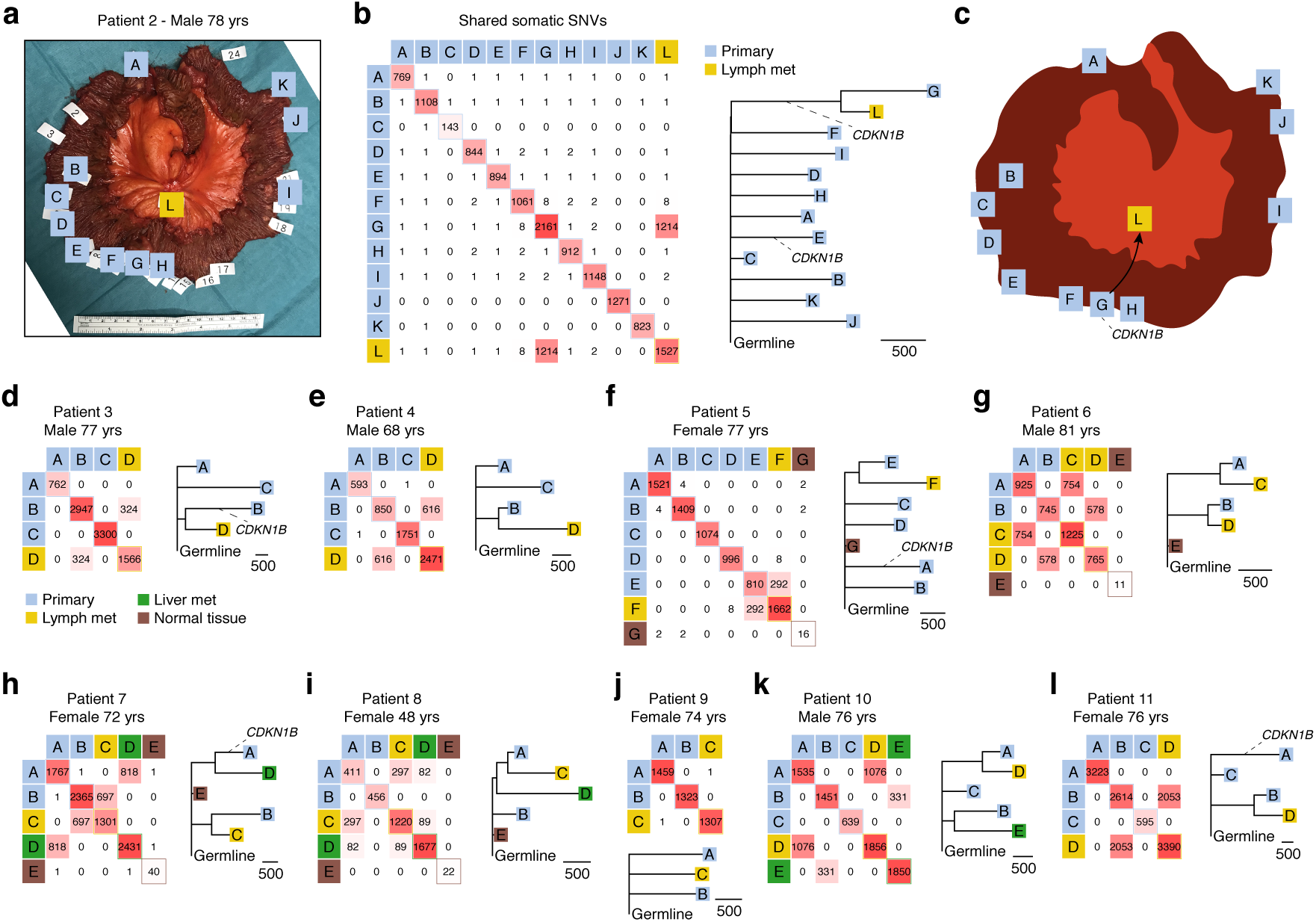
Whole genome sequencing of 50 primary tumors and metastases from 10 additional SI-NET patients confirms independent clonal origin. (**a**) Section of the small intestine from a 78 year-old male patient harboring 11 primary tumors (A-K) and one lymph node metastasis (L). White labels indicate all identified tumors while letters indicate sequenced samples with sufficient purity and tumor material. (**b**) Pairwise analysis of shared somatic SNVs. Primary tumor G and the metastasis shared 1219 mutations. The known *CDKN1B* driver event is indicated in the phylogenetic tree (the number of SNVs in each branch is given by the scale marker). (**c**) Proposed model. (**d-l**) Similar to panel b, based on archival material from 9 additional patients, ages 48-81, that also included liver metastases and normal mucosa samples. Bootstrap support was 100% for all major branches in the phylogenetic trees. Whole genome sequencing was performed at 29.8-45.0x coverage.

Mirroring the result from Patient 1, shared somatic mutations were essentially absent in between intestinal tumors in all 10 additional patients (**Fig. 2**). In contrast, strong SNV overlaps were seen between individual metastases and specific intestinal tumors, ranging from 82 to 2053 and above 290 SNVs in all pairs but one. For example, in Patient 2, a single intestinal tumor (G) out of 11 that were sampled showed a striking relatedness to an adjacent lymph node metastasis (L; 1,219 shared SNVs; **Fig. 2b**). Similar to Patient 1, the metastatic primary was centrally located in the resected section (**Fig. 2a,c**). Detailed positional data was lacking for the remaining patient samples, as these were derived from archival material.

While individual metastases showed credible overlaps only with a single primary tumor, in three cases (Patients 6, 7 and 10) we found that two independent primary tumors had metastasized to different lymph node or liver metastases (**Fig. 2g,h,k**). Given the lack of a common evolutionary trajectory for the primary tumors, this supports that acquisition of metastatic properties is not a rare event in multifocal SI-NET. In Patient 8, the sample phylogeny suggested two independent metastatic events from a single primary (A), with an earlier event giving rise to a liver metastasis (D) and a later event resulting in a lymph node metastasis (C; **Fig. 2i**). Shared SNVs typically had higher variant allele frequencies (VAFs) than other variants, in both primaries and metastases, consistent with continued somatic evolution and subclonal expansions following the metastatic events (**Supplementary Fig. 6**). In a single case (Patient 9), a lymph node metastasis could not be associated to any of two available intestinal tumors, likely explained by incomplete sampling of intestinal lesions (**Fig. 2j**).

False shared somatic SNVs can arise due to the fact that all samples from a given patient use the same blood normal sequencing data. However, only a small number of additional common SNVs were observed beyond the main genetic relationships (**Fig. 1b** and **Fig. 2b,d-l**), some of which could be dismissed as false positives by manual inspection (**Supplementary Table S3**). In Patient 5, one additional tumor (D) shared 8 credible SNVs with the metastasis (F) (**Fig. 2f**). This overlap grew to 66 when high-confidence variants were whitelisted for relaxed calling in other samples, allowing more sensitive detection of subclonal shared SNVs (**Supplementary Fig. 7**). These variants were present as low-VAF traces in the metastasis while having normal VAFs in the tumor, compatible with contamination during sample handling or, possibly, metastatic spread to the same lymph node from two tumors with a minor contribution from D (**Supplementary Fig. 8**). Low-VAF variants (13 in total) shared between an intestinal tumor (A) and a lymph node metastasis (C) were also uncovered in Patient 9 (**Supplementary Fig. 9**). In Patient 2, 8 SNVs were shared between the main metastatic primary (G), an adjacent primary (F, 5 mm apart, **Supplementary Fig. 10**) and the metastasis (L) (**Fig. 2b**). All had normal VAF distributions in all 3 samples arguing against contamination **(Supplementary Fig. 11**) and all but one passed manual inspection (**Supplementary Table S3**). Overlaps of this size are too small to represent common clonal origin in a late-onset cancer, and are likely explained by lineage-specific somatic mutations established early during organismal development (Lodato et al., 2015).

### Driver mutations and mutational signatures

The median age of the patients was 76 years (one subject, Patient 8, was 48 while others ranged from 68 to 81). The observed burdens and number of pairwise overlapping variants are thus roughly on par with a mutation rate similar to healthy human neurons, estimated to accumulate ∼23-40 SNVs/year (Lodato et al., 2018) (**Fig. 3**). Analysis of mutational signatures indeed supported major contributions from COSMIC Signature SBS5, an ubiquitous aging-associated mutational process, or SBS40 which is closely related to SBS5 (Alexandrov et al., 2020) (**Fig. 3**). The observed variability in signature loadings may in part be methodological, since the trinucleotide substitution profiles of the samples were in practice highly similar across the cohort (**Supplementary Fig. 12**). Within-patient burden variability was to a large degree explained by variable sample purity (**Supplementary Fig. 13**). These results confirm the mutationally quiet nature of SI-NET.

**Figure 3.**
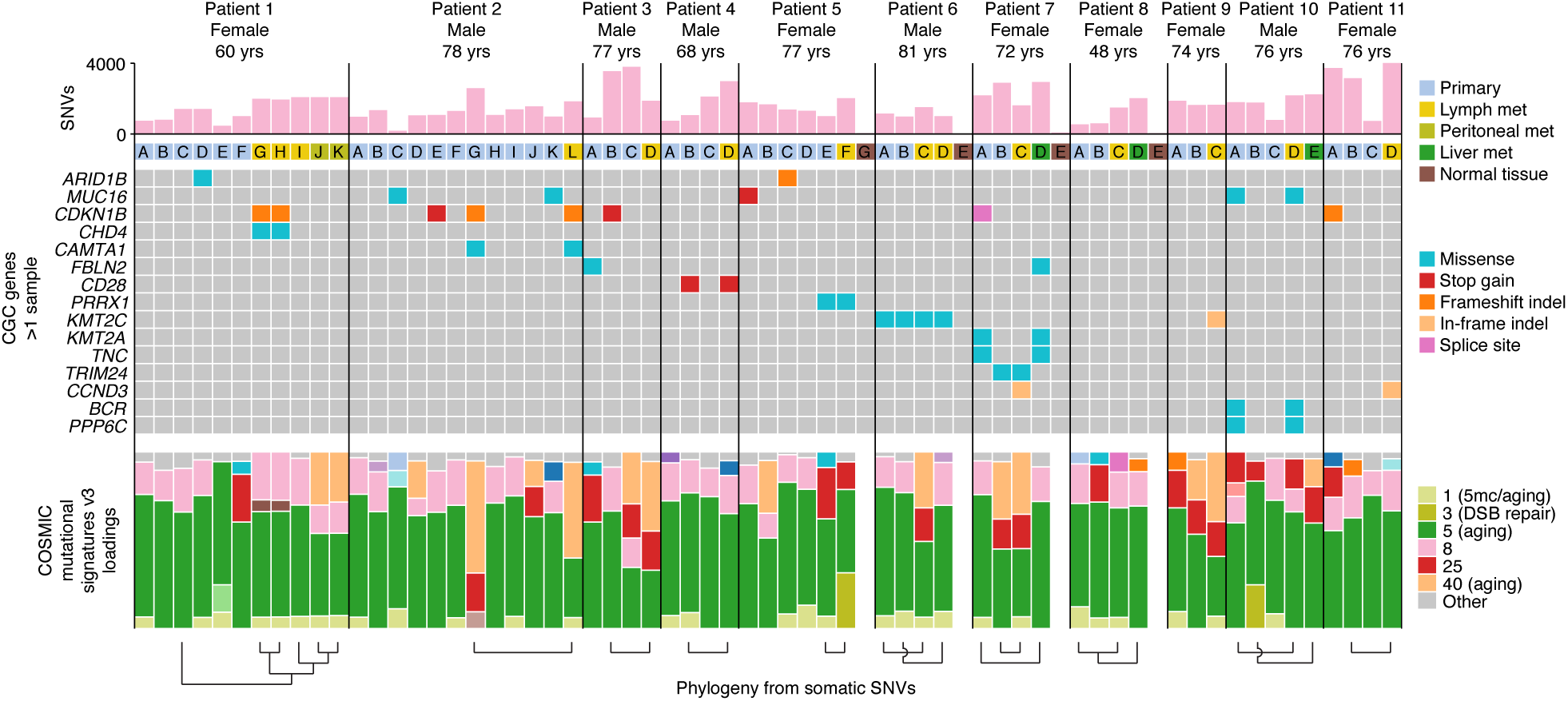
Overview of potential driver mutations and mutational signatures for all 11 patients. All CGC genes with non-synonymous mutations in more than one sample are shown. Mutational signature loadings, i.e. relative contributions, refer to COSMIC v3 definitions (Alexandrov et al., 2020) and were dominated by the closely related signatures SBS5 and SBS40 (see **Supplementary Fig. 16** for a full signature legend). Phylogenetic relationships previously inferred from somatic SNVs (**Fig. 1-2**) are indicated at the bottom. Somatic SNV burdens, potential driver mutations (Cancer Gene Census genes mutated in >1 sample) and mutational signature loadings in this figure were determined using less rigorous population variant filtering compared to the phylogenetic analyses (see **Materials and Methods**). CGC, Cancer Gene Census.

Analysis of potential somatic driver mutations in coding genes mirrored earlier published results, with infrequent variants in *CDKN1B* emerging as the main recurrent event (8 samples, 6 independent events; **Fig. 3; Supplementary Table S4**) (Francis et al., 2013). In Patient 1, the same *CDKN1B* frameshift indel occurred in two of the metastases (G, H) in a pattern consistent with the inferred phylogeny (**Fig. 1b-c**). In Patient 2, the same *CDKN1B* frameshift indel was found in the metastatic primary (G) and the metastasis (L), thus again consistent with the phylogeny, while one non-metastatic primary carried a *CDKN1B* stop gain variant (**Fig. 2b-c**). Patients 3 and 7 carried *CDKN1B* variants (stop gain and splice donor) in metastatic primary tumors that were missing in the corresponding metastases (**Fig. 2d,h**). These variants were among a large number of private variants present at relatively low VAF in the primaries, and may have arisen after metastasis (**Supplementary Figs. 14 and 15**). Other mutated Cancer Gene Census (Tate et al., 2019) genes included *MUC16* (5 samples, 4 independent events), a common false positive gene encoding the second largest human protein (Lawrence et al., 2013), *KMT2C* (5 samples, 3 independent events), *FBLN2* (2 independent events) and *ARID1B* (2 independent events; **Fig. 3**). *TERT* promoter mutations, frequent non-coding driver events in several cancer types (Horn et al., 2013; Huang et al., 2013), were absent, as were notable upstream mutations in other cancer genes (**Supplementary Fig. 16**). Consistent with other reports, there was thus a lack of obvious driver mutations beyond *CDKN1B*, which was not essential for metastasis.

### Copy number alterations

Analysis of somatic CNAs gave further support for the phylogenetic relationships inferred from somatic SNVs (**Fig. 4a**). The metastatic tumor in Patient 1 (C) carried a distinct segmental loss on chr11 found in all metastases (G-K) but not the other intestinal tumors. Similarly, the metastatic tumor in Patient 2 (G) exhibited chromothripsis on chr13 (abundant clustered CNAs that oscillated between two states (Korbel and Campbell, 2013), **Supplementary Fig. 18**), and this exact complex pattern was mirrored in the corresponding lymph node metastasis (L) (**Fig. 4b**). Patient 2 and 7 had alterations on chr11 and chr20, respectively, that were present in primary tumors but absent in related metastases and thus presumably occurred after metastasis. Approximately 70-75% of SI-NETs have been shown to harbor hemizygous loss of chr18 (Andersson et al., 2009), and we accordingly observed chr18 loss in 9 of the 11 patients and in 32 of the 61 tumor samples. However, phasing of chr18 loss based on germline single nucleotide polymorphisms (SNPs) revealed that tumors deemed unrelated based on SNVs often had undergone loss of different chromosome homologs, while related tumors always showed loss of the same homolog (*P* = 1.2×10^-4^, binomial test; **Fig. 4a,c** and **Supplementary Fig. 17**). Other whole chromosome events (4, 5, 14 and 20) showed similar patterns of concordant/discordant chromosome homolog loss or gain in agreement with the established phylogenies (**Fig. 4a**). Among other recurrent events were segmental deletions on chr11 and chr13, as shown previously (Banck et al., 2013; Francis et al., 2013; Walter et al., 2018). No distinctive CNAs were shared in ways that contradicted the SNV-based evolutionary trees (**Fig. 4a**).

**Figure 4.**
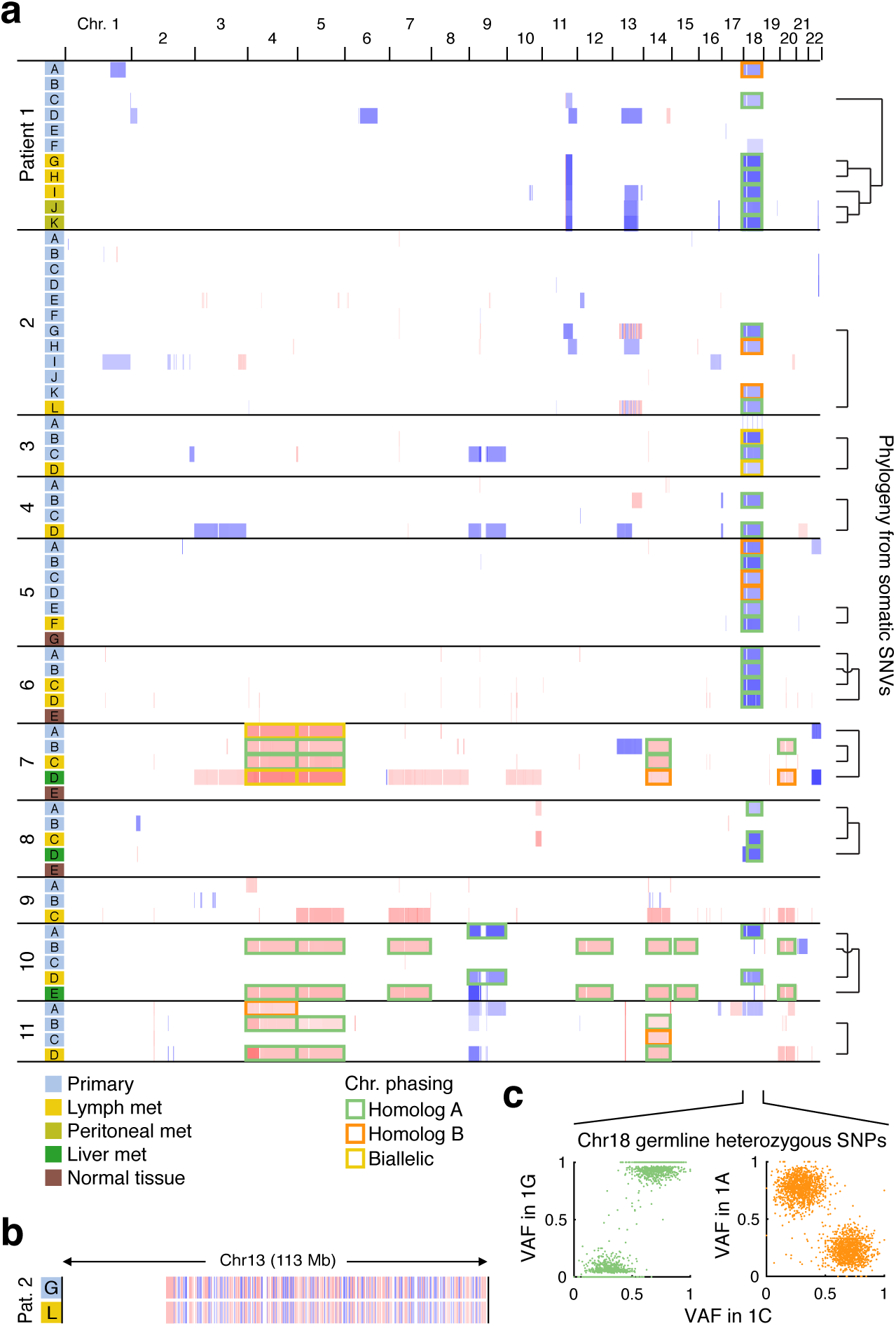
Somatic copy number alterations agree with phylogenies inferred from somatic SNVs. (**a**) Somatic CNA profiles for all samples (blue = deletion, red = amplification; segments <25 kb were excluded for visualization purposes). Phylogenetic relationships previously inferred from somatic SNVs (**Fig. 1-2**) are indicated to the right. Whole-chromosome losses and gains were phased based on germline SNPs (see **Supplementary Fig. 16**), with the two chromosome homologs indicated in green and orange and likely biallelic events shown in yellow. (**b**) Detailed view of chromothripsis on chr13 in Patient 2, present in the metastatic primary (G) and the metastasis (L). (**c**) Example of phasing of chr18 deletions in Patient 1.

Chromothripsis, not previously reported in SI-NET to our knowledge, was further supported by analysis of somatic structural alterations. Of 27 in-frame structural events, none of which were obvious drivers, 9 were intrachromosomal alterations on chr13 in Patient 2, all found in both the primary tumor and the metastasis shown by CNA to harbor chromothripsis on this chromosome (G and L; **Supplementary Fig. 19**).

## Discussion

We find that in multifocal SI-NET, tumor development is initiated in multiple clonally independent cells, leading to a group of tumors that are unrelated in terms of somatic genetic evolution. Furthermore, we encountered several cases where two independent intestinal tumors had metastasized, each giving rise to a distinct liver or lymph node metastasis. This suggests that acquisition of metastatic properties is not a rare event in multifocal SI-NET and underscores the importance complete surgical removal of all intestinal lesions. Previous exome-based analyses of paired single intestinal and metastatic samples have yielded puzzling results with a highly varying degree of genetic overlap and in some cases no overlap at all (Francis et al., 2013; Walter et al., 2018), seen in one patient also in the present study, and our results thus suggest that the relevant primary tumors have not been sampled in such cases.

Multifocal entero-pancreatic carcinomas are typically seen in hereditary cancer syndromes such as MEN-1 and familial adenomatous polyposis (FAP). Cohort studies support that some SI-NETs may have a heritable component (Dumanski et al., 2017; Walsh et al., 2011), and multifocality has been suggested to occur preferentially in familial cases (Sei et al., 2016; Sei et al., 2015). However, multifocality is common in all SI-NET (Choi et al., 2017; Gangi et al., 2018), and key clinical parameters such as survival or age of diagnosis, which is typically lower for hereditary cancer, are similar in patients with or without multifocality (Choi et al., 2017; Gangi et al., 2018). The observation that the most recurrent somatic genetic event in SI-NET, chr18 loss, can affect both the maternal and the paternal chromosome homologs in the same patient argues against contribution from a predisposing chr18 germline variant by loss of heterozygosity (Zhang et al., 2020). Furthermore, in stark contrast to germline-induced gastrointestinal tumors, SI-NET only affects a limited intestinal segment. Given the general lack of established genetic drivers, and in the light of our results showing clonal independence, we suggest that future studies could focus on cancer-priming local factors that may contribute to emergence of multifocal SI-NET. Such factors could theoretically include early embryonic genetic events present in blood as well as select EC cell lineages, thus precluding their identification as somatic events in the present study, or local environmental factors. While such factors remain elusive, it can be noted that SI-NET originates from epithelial enterochrommafin cells (EC-cells) that intrinsically interact with their surroundings by paracrine and endocrine serotonin secretion, synaptic-like connections with the enteric nervous system, and receptor-mediated nutrient sensing of luminal content (Bellono et al., 2017).

## Acknowledgements

The work described here was supported by the Swedish Research Council (E.L.), the Swedish Cancer Society (E.L. and O.N.), the Knut and Alice Wallenberg Foundation (E.L.), the BioCARE National Strategic Research Programme (O.N.), and grants from the Swedish state under the agreement between the Swedish government and the county councils, the ALF agreement (E.E. and O.N.). We wish to thank laboratory assistant Gulay Altiparmak, research nurse Maria Nilsson and surgical coordinator Jenny Oliver for skilled technical assistance, and Kerryn Elliott for critical reading of the manuscript. We further want to thank professor emeritus Bo Wängberg as well as previous and current surgical staff at the Section of Endocrine and Sarcoma Surgery, Department of Surgery, Sahlgrenska University Hospital.

## Competing interests

None.

## Supplementary appendix

### Supplementary Materials and Methods

#### Immunohistochemistry

Sections of all sampled tumors from 11 patients with SI-NET were subjected to antigen retrieval using EnVision FLEX Target Retrieval Solution (high pH) in a Dako PT-Link. Immunohistochemical staining was performed in a Dako Autostainer Link using EnVision FLEX according to the manufacturer’s instructions (DakoCytomation).

#### Mapping and somatic variant calling

The reads were mapped to the human reference genome (hg19 assembly) using BWA as part of Sentieon Genomics Tools (bwa-mem v0.7.15.r1140 for patients 1-5 and 0.7.17.r1188 for patients 6-11), which further also performs deduplication, realignment and sorting. Mutect2 was run using default parameters, and the output was fed into the filterMutectCalls tool with a mean median mapping quality score requirement of 10. With VarScan, the somatic tool was used to call variants by default cutoffs and a minimum variant allele frequency of 0.01 from SAMtools (v1.9) pileups of reads with Phred-scaled mapping quality >= 15 and variant base quality >= 20. The VarScan processSomatic tool was then used to get only the somatic SNVs and indels, where a minimum support of two reads was required for a variant to be called. The somaticFilter tool was also used to discard SNVs called within SNP clusters or within 3 bp of somatic and germline indels. Variants were further filtered to have coverage >=20 reads in the normal, to be present on both strands, and to be absent in the normal. The final set of variants was yielded by intersecting outputs from the two callers, followed by annotation using ANNOVAR (v2019Oct24) (Wang et al., 2010) and Variant Effect Predictor (VEP) (McLaren et al., 2010). To minimize false positive calls in phylogenetic analyses, population variants in dbSNP150 (provided by ANNOVAR) and the ENSEMBL variant database (provided by VEP) were removed. Analysis of potential driver mutations and mutational signatures was based on less stringent population variant filtering (dbSNP138), to avoid false negatives and since extensive SNP filtering can skew results from mutational signatures analyses. For high-sensitivity analysis of pairwise shared variants, mutations detected by the standard pipeline as described above in any of the two samples in a pair were whitelisted for detection using the unfiltered VarScan somaticFilter output, allowing more sensitive detection of overlapping subclonal mutations.

#### Sample purity and telomer lengths

Samples were assed for tumor cell content using PurBayes (Larson and Fridley, 2013) and telomere lengths were determined using Computel (Nersisyan and Arakelyan, 2015) using default settings, both presented in **Supplementary Table S2**.

#### Viral reads and transposition events

Analysis of WGS reads for viral content, which produced only low counts of expected *Herpesviridae* family reads or reads consistent with plasmid vector contamination thus giving no support for a role for DNA viruses in SI-NET, was performed using a previously established pipeline (Tang et al., 2013). TraFiC-mem was used for detection of somatic mobile element insertions (Tubio et al., 2014), which revealed only a single LINE1 transposition event (in *BCAS3* in sample 5F).

**Supplementary Figure 1.**
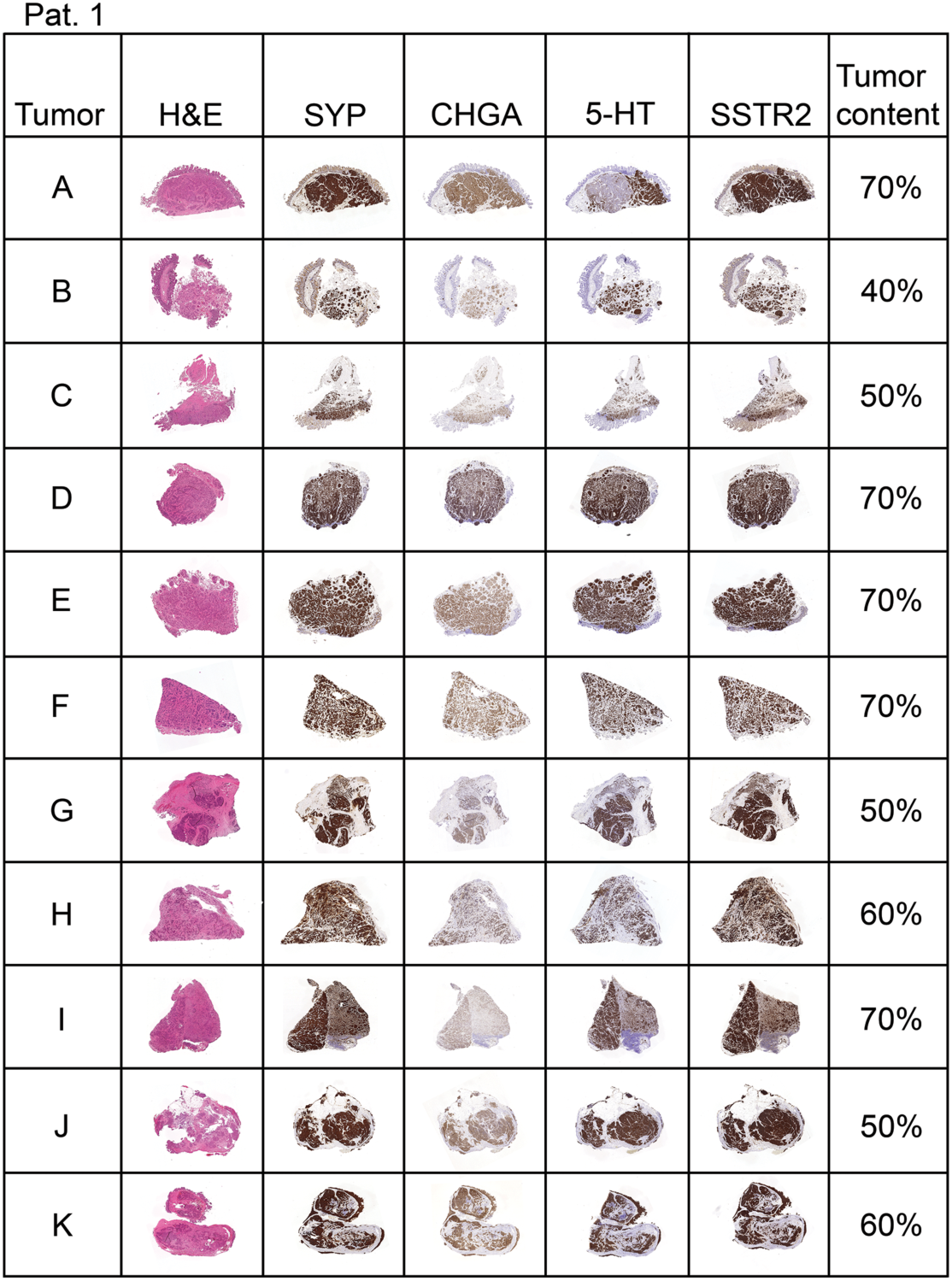
Immunohistochemical analysis of all sequenced samples from Patient 1. All tumors display similar morphology, and all express the neuroendocrine markers, SYP (synaptophysin), CHGA (chromogranin A) and 5-HT (serotonin) as well as the clinically relevant SSTR2 (somatostatin receptor 2). The assessment of tumor cell content was performed visually on hematoxylin and eosin-stained (H&E) sections of the tumors. The letter in each row indicates the sample ID.

**Supplementary Figure 2.**
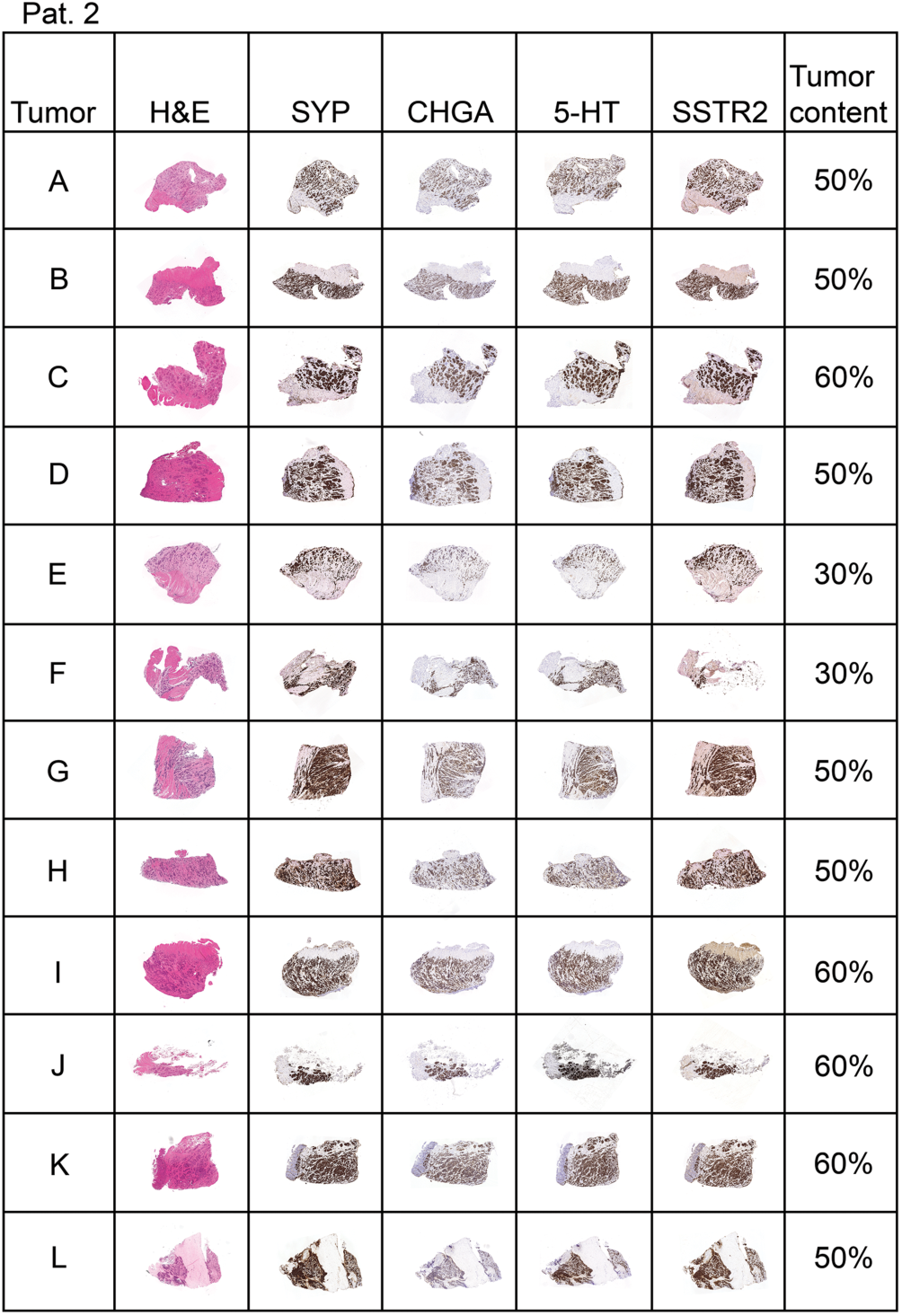
Immunohistochemical analysis of all sequenced samples from Patient 2. All tumors display similar morphology, and all express the neuroendocrine markers, SYP (synaptophysin), CHGA (chromogranin A) and 5-HT (serotonin) as well as the clinically relevant SSTR2 (somatostatin receptor 2). The assessment of tumor cell content was performed visually on hematoxylin and eosin-stained (H&E) sections of the tumors. The letter in each row indicates the sample ID.

**Supplementary Figure 3.**
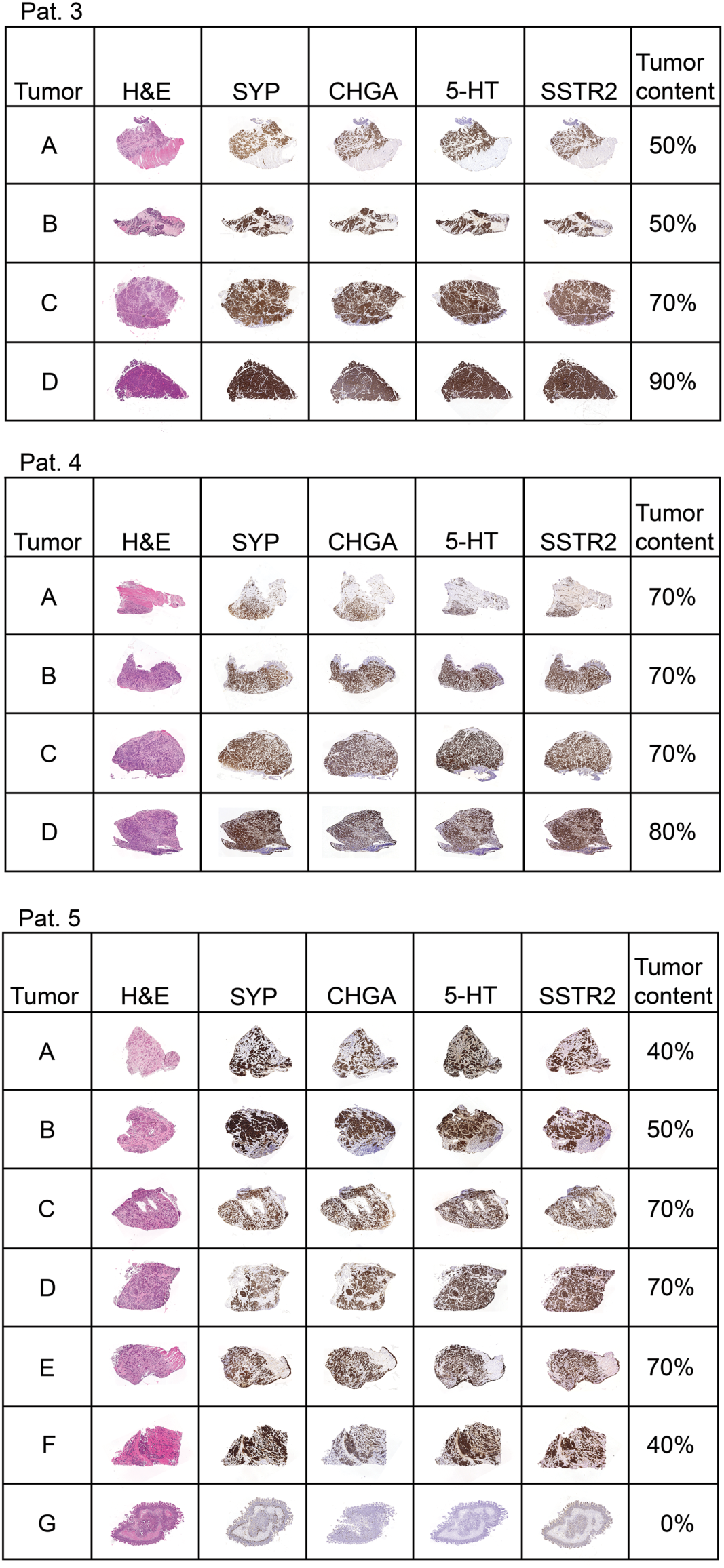
Immunohistochemical analysis of all sequenced samples from Patient 3-5. All tumors display similar morphology, and all express the neuroendocrine markers, SYP (synaptophysin), CHGA (chromogranin A) and 5-HT (serotonin) as well as the clinically relevant SSTR2 (somatostatin receptor 2). The assessment of tumor cell content was performed visually on hematoxylin and eosin-stained (H&E) sections of the tumors. The letter in each row indicates the sample ID. Sample 5G is from normal intestinal mucosa.

**Supplementary Figure 4.**
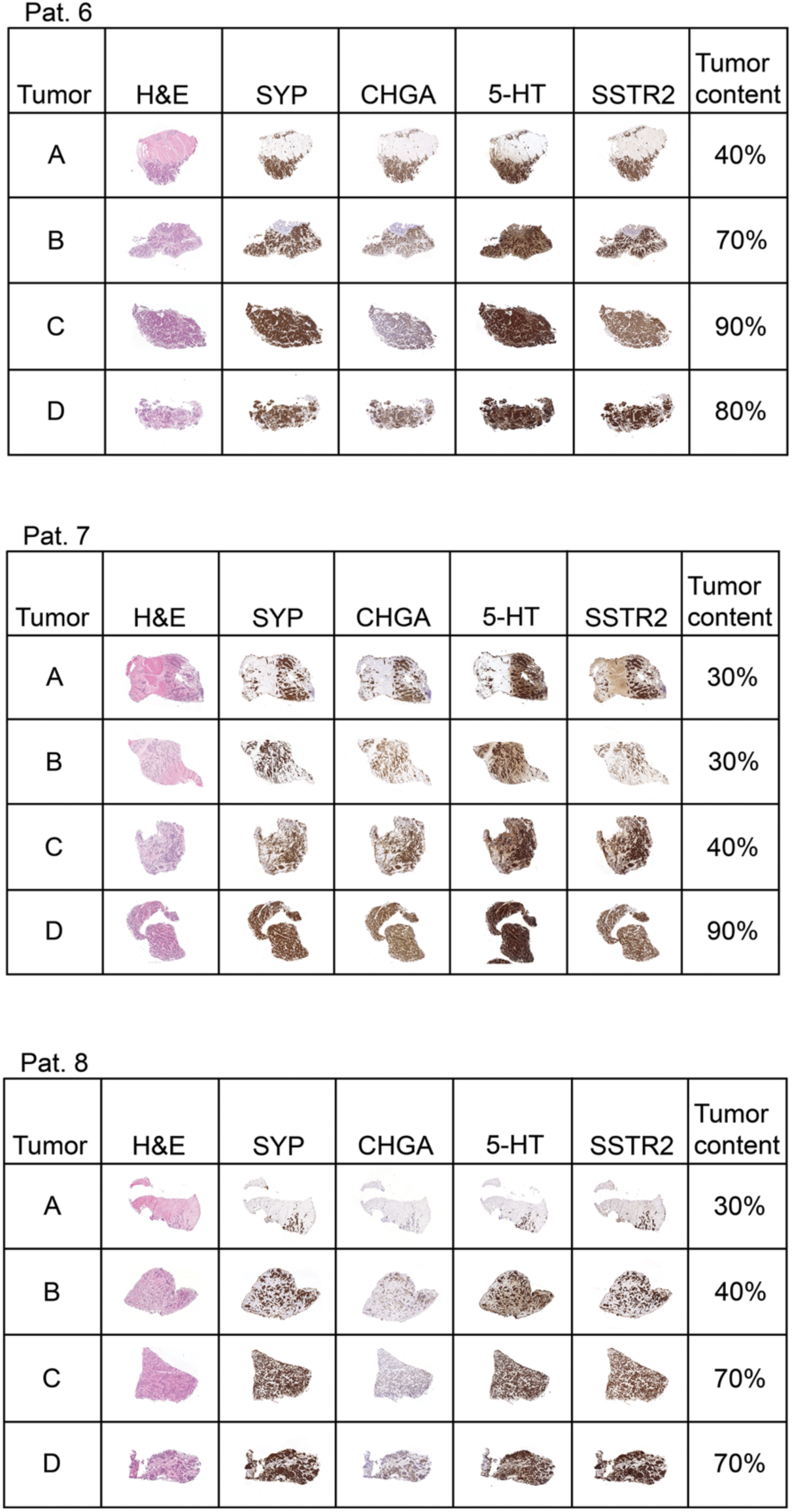
Immunohistochemical analysis of all sequenced samples from Patient 6-8. All tumors display similar morphology, and all express the neuroendocrine markers, SYP (synaptophysin), CHGA (chromogranin A) and 5-HT (serotonin) as well as the clinically relevant SSTR2 (somatostatin receptor 2). The assessment of tumor cell content was performed visually on hematoxylin and eosin-stained (H&E) sections of the tumors. The letter in each row indicates the sample ID.

**Supplementary Figure 5.**
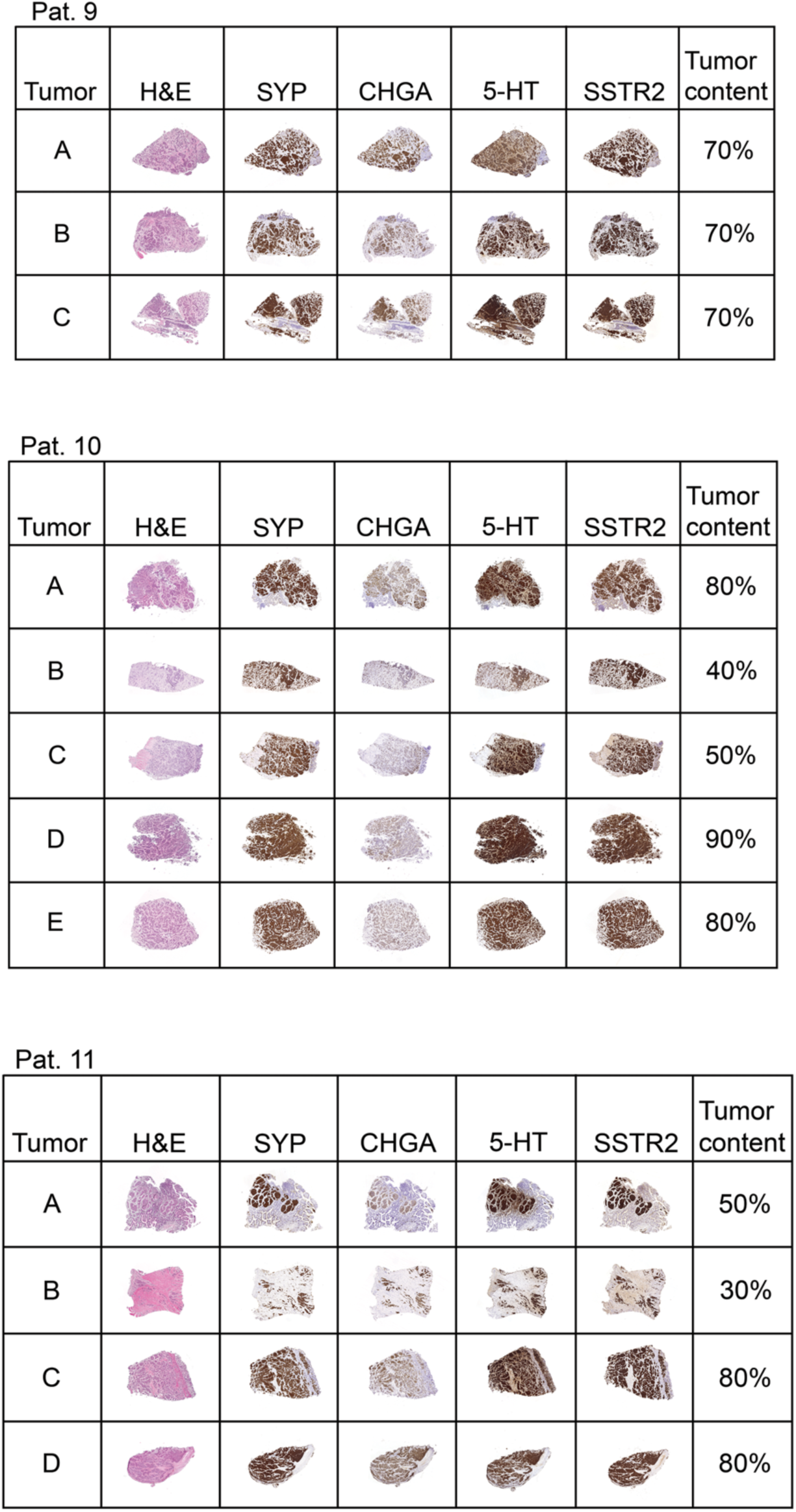
Immunohistochemical analysis of all sequenced samples from Patient 9-11. All tumors display similar morphology, and all express the neuroendocrine markers, SYP (synaptophysin), CHGA (chromogranin A) and 5-HT (serotonin) as well as the clinically relevant SSTR2 (somatostatin receptor 2). The assessment of tumor cell content was performed visually on hematoxylin and eosin-stained (H&E) sections of the tumors. The letter in each row indicates the sample ID.

**Supplementary Figure 6.**
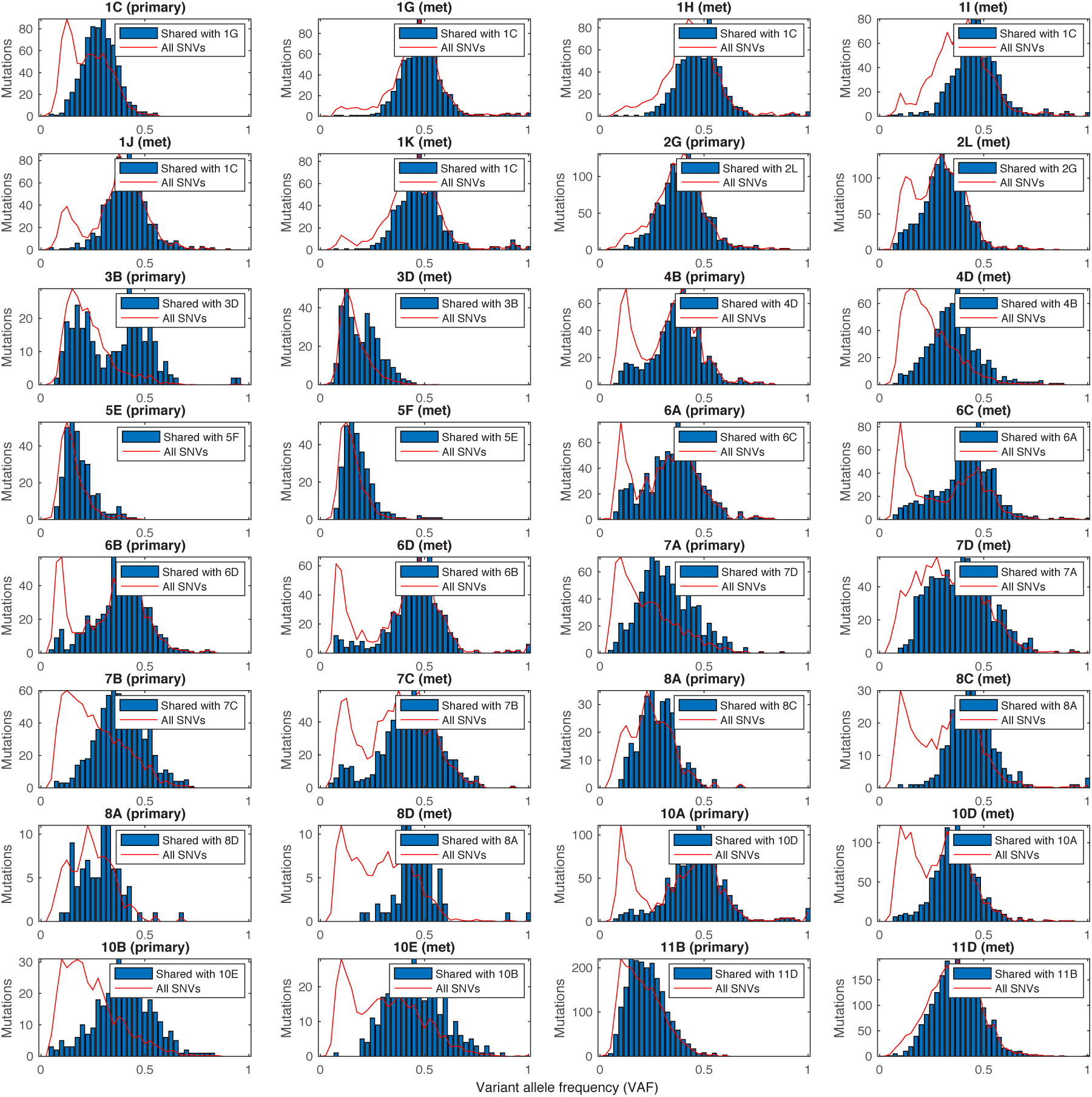
Analysis of variant allele frequency (VAF) distributions of primary-metastasis shared SNVs. VAF distribution plots of SNVs shared between primary tumors and corresponding metastases in all relevant samples. The overall VAF distribution for all high-confidence mutations in each sample is shown as reference (red; normalized to fit y-axis scale). More detailed views and discussions of Patient 3 and Patient 7 are provided in **Supplementary Figs. 14 and 15**.

**Supplementary Figure 7.**
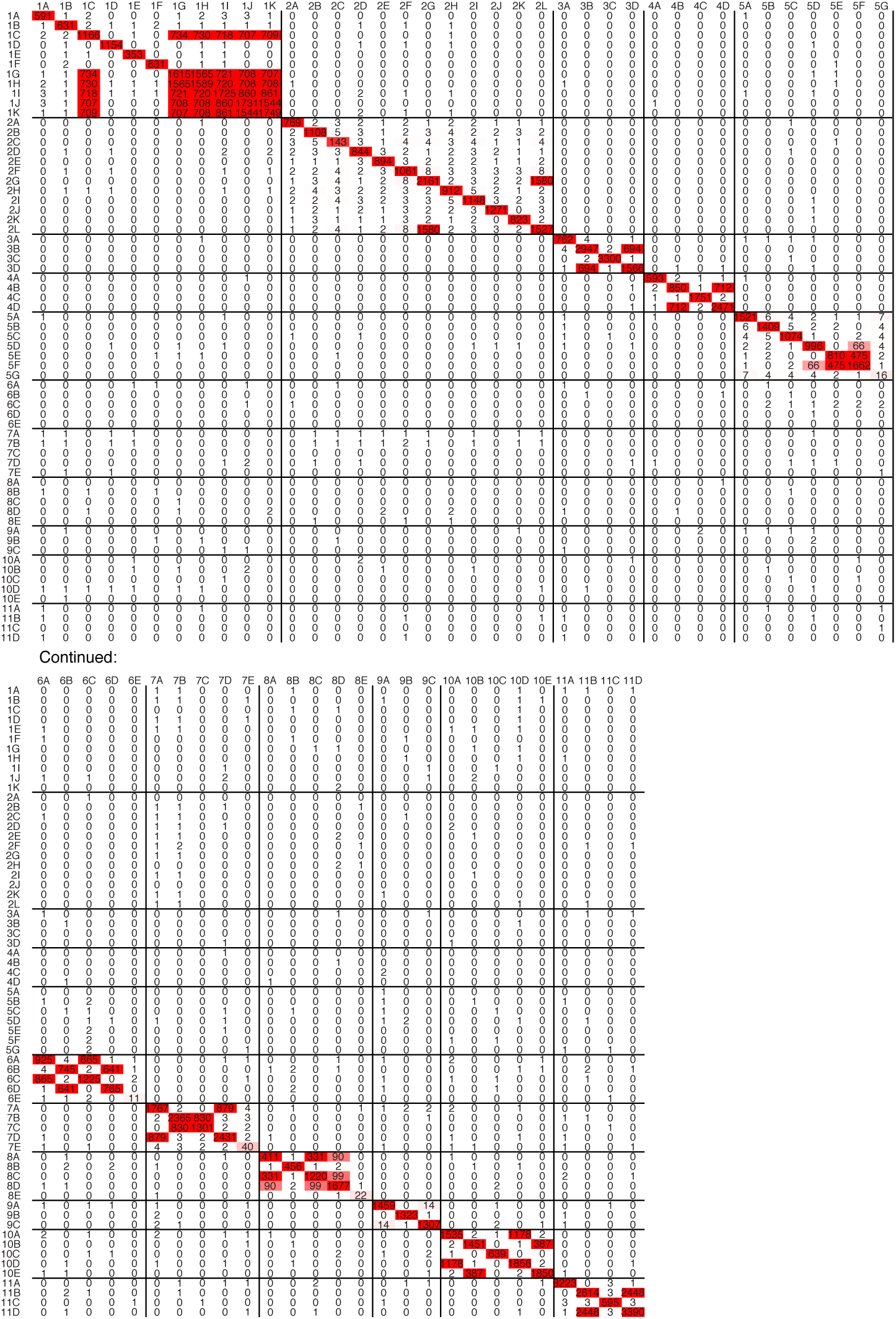
High-sensitivity search for shared variants by whitelisting of SNVs called at high confidence in at least one sample. A two-step procedure was used, where any somatic SNVs called at high confidence (detected by both Mutect and VarScan using various filters including a strand filter as described in Materials and Methods) in at least one sample were “whitelisted”. This was followed by a more sensitive search for presence of the whitelisted variants using VarScan without stringent filters (e.g. exclusion of strand filter, see Materials and Methods). The number and letter in each row/column label indicates the patient number and sample ID, respectively.

**Supplementary Figure 8.**
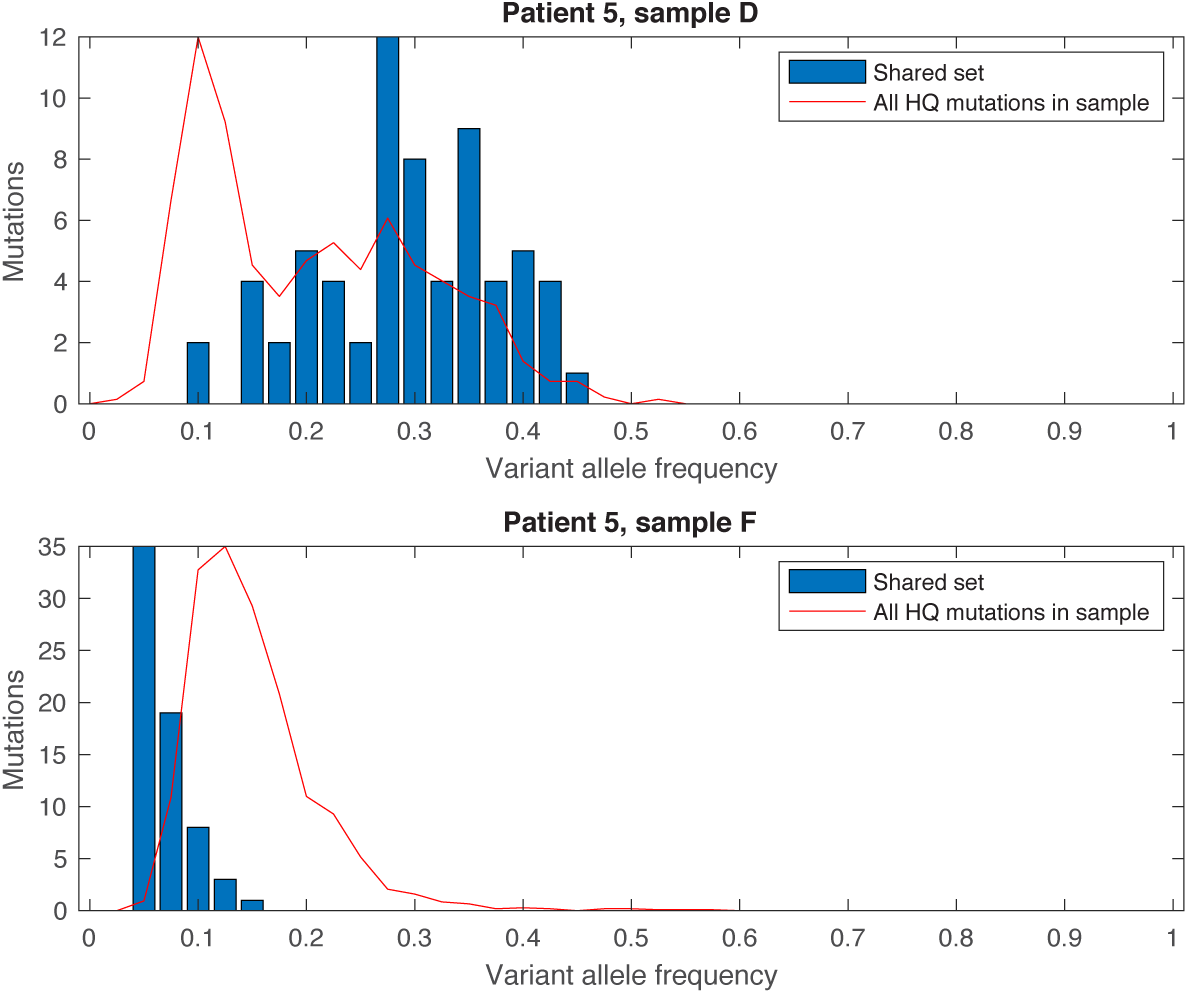
Variant allele frequency (VAF) distributions indicate low-level contamination between samples D and F from Patient 5. Samples D (primary tumor) and F (lymph node metastasis) in Patient 5 shared 8 SNVs in the main phylogenetic analysis (main Fig. 2). A high-sensitivity search for shared variants (**Supplementary Fig. 7**) revealed 58 additional shared variants called at high confidence in at least one of the samples while being detectable using relaxed filters (e.g. exclusion of strand filter, see Materials and Methods) in the other sample. These 66 shared SNVs were present at high VAFs in sample D (top histogram) while being highly subclonal in sample F (bottom histogram). This supports that the metastasis sample (F) was contaminated with DNA from D, for example during sample handling or, alternatively, that trace amounts of material from D reached F through metastatic spread. Overall VAF distributions for all high-quality mutations in each sample is shown for comparison (red; normalized to fit y-axis scale).

**Supplementary Figure 9.**
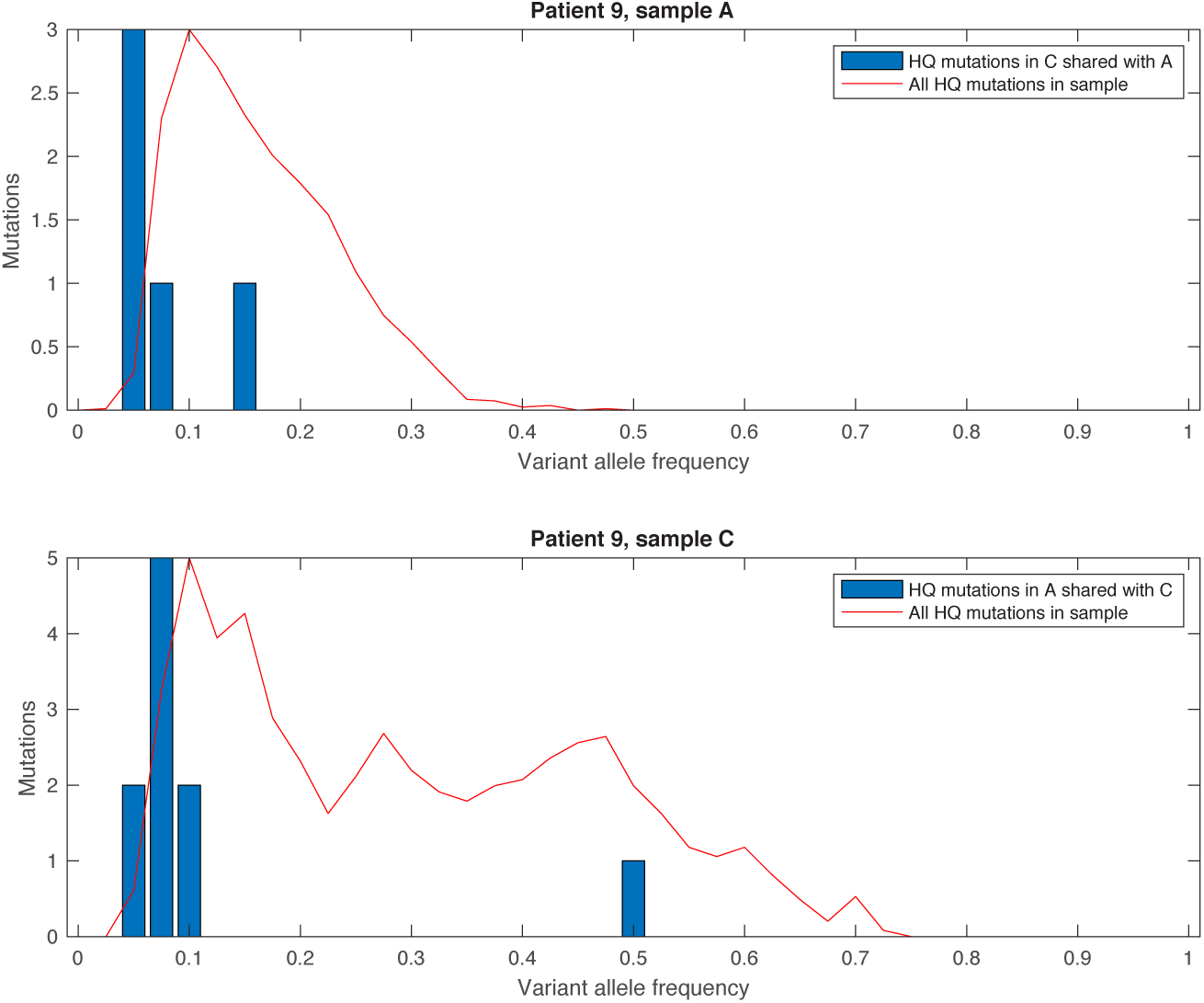
Variant allele frequency (VAF) distributions indicate low-level contamination between samples A and C from Patient 9. A high-sensitivity search for subclonal shared SNVs (**Supplementary Fig. 6**) revealed 14 variants shared between samples A (primary tumor) and C (lymph node metastasis) in Patient 9, all present at high confidence in at least one of the samples while being detectable using relaxed filters (e.g. exclusion of strand filter, see Materials and Methods) in the other. The graphs show that variants present at high confidence in A was generally found at low VAF in C, and vice versa. A likely explanation is sample contamination, although transfer of trace amounts of tumor material through metastatic spread could in principle also explain these patterns. Overall VAF distributions for all high-quality mutations in each sample is shown for comparison (red; normalized to fit y-axis scale).

**Supplementary Figure 10.**
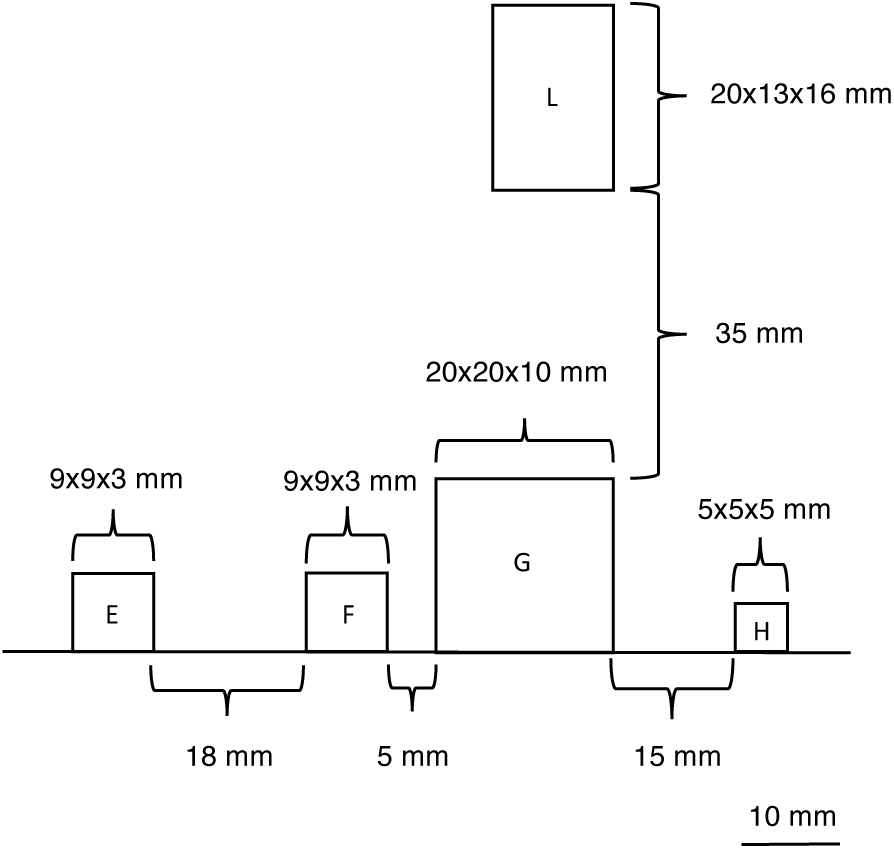
Physical positioning of tumors F, G and L in Patient 2.

**Supplementary Figure 11.**
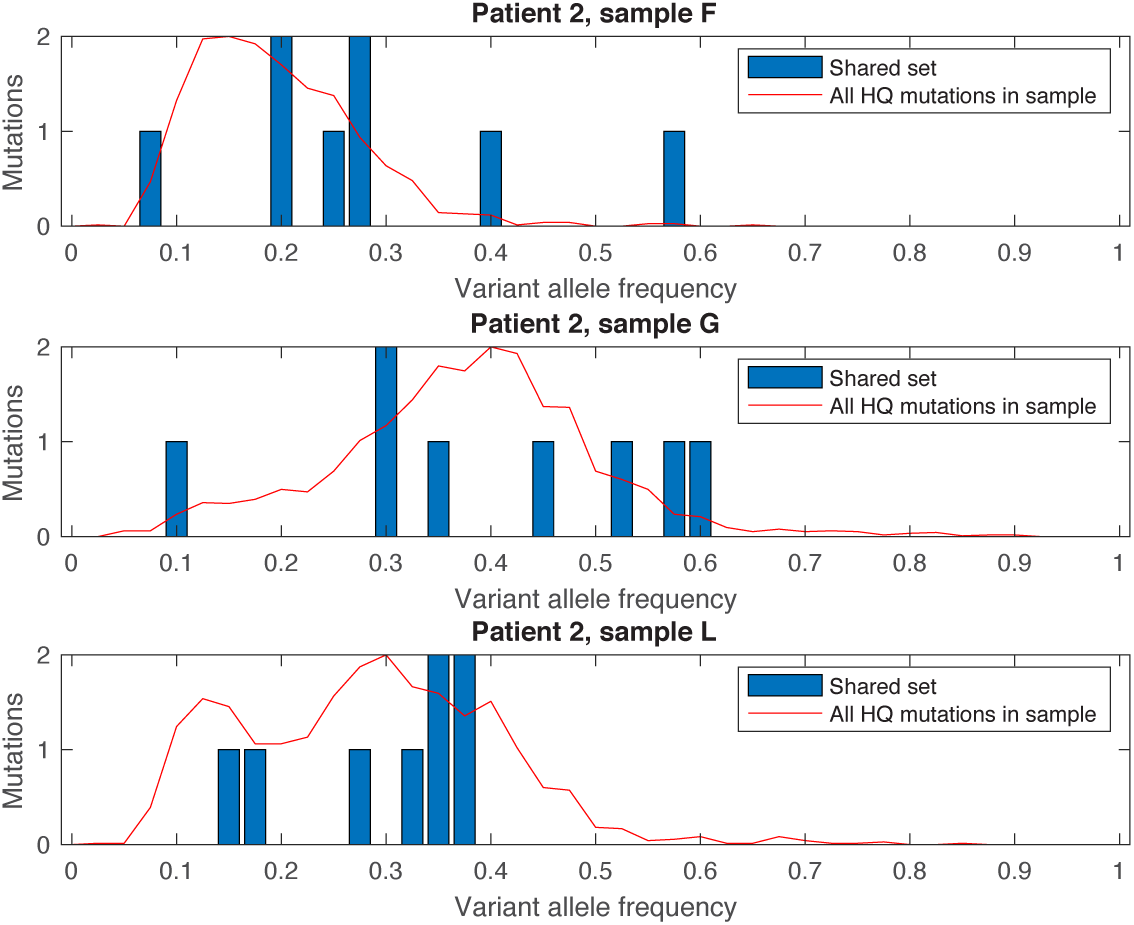
Variant allele frequency (VAF) distributions for 8 variants found in samples F, G and L from Patient 2. A set of 8 SNVs was shared between samples F, G and L (lymph node metastasis) in Patient 2 (main **Fig. 2**). This number stayed constant when using a high-sensitivity approach to uncover subclonal shared SNVs (**Supplementary Fig. 7**). Manual inspection in IGV supported that the shared calls were mostly of high quality, with only one having characteristics of being a false positive call (**Supplementary Table S3**). The shared SNVs showed normal VAF distributions (comparable to other mutations) in all samples. The overall VAF distribution for all high-quality mutations in each sample is shown for comparison (red; normalized to fit y-axis scale).

**Supplementary Figure 12.**
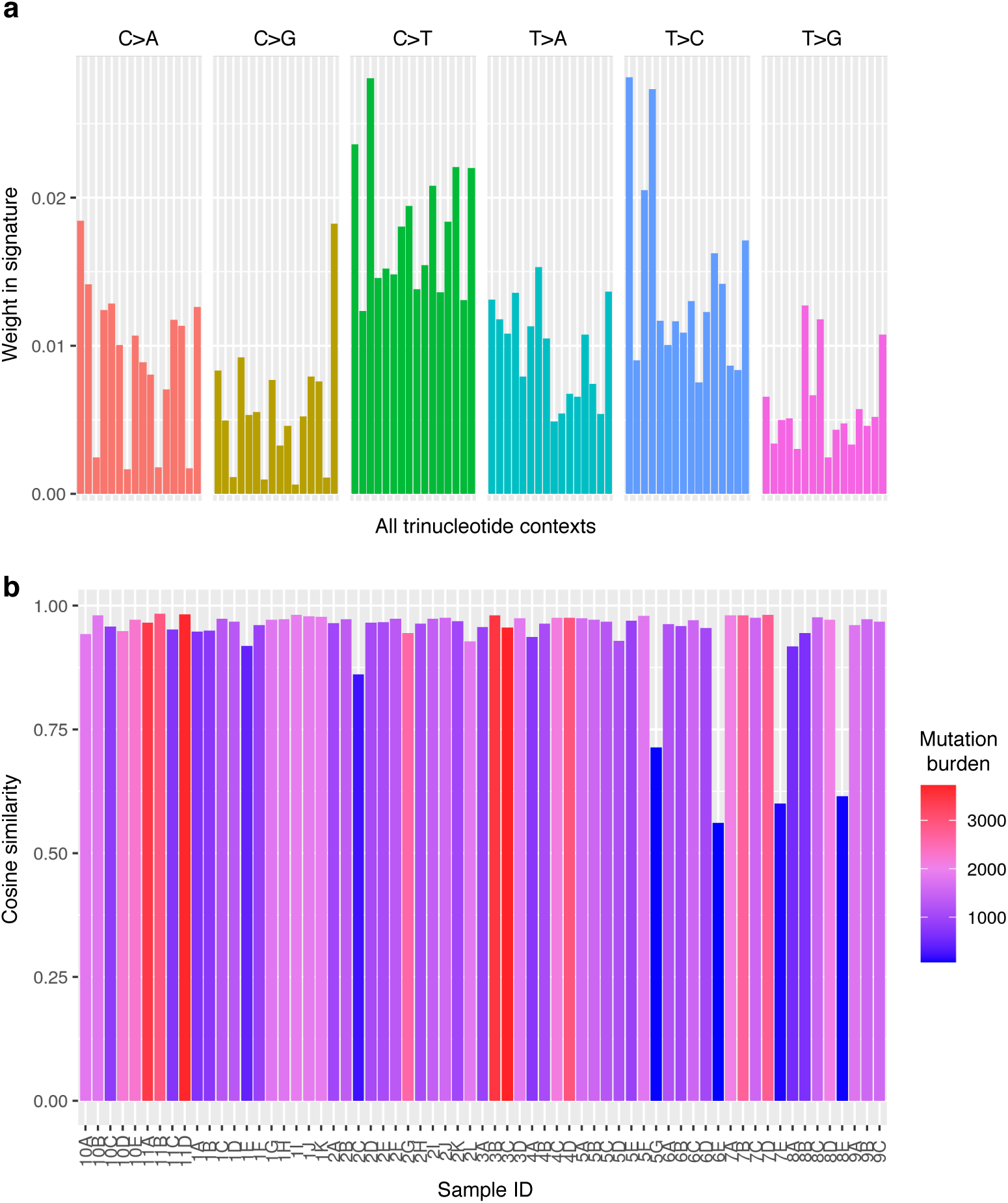
Highly coherent trinucleotide substitution profile across the cohort. (**a**) Average trinucleotide substitution profile in the cohort. (**b**) Cosine similarity of per-sample trinucleotide profiles and the cohort average shown in (a). Most samples showed high similarity, and deviation from the average profile was associated with reduced mutation burden, thus reducing the ability to accurately determine the signature. This is most clearly seen for the normal mucosa tissue samples, which all have low mutation counts.

**Supplementary Figure 13.**
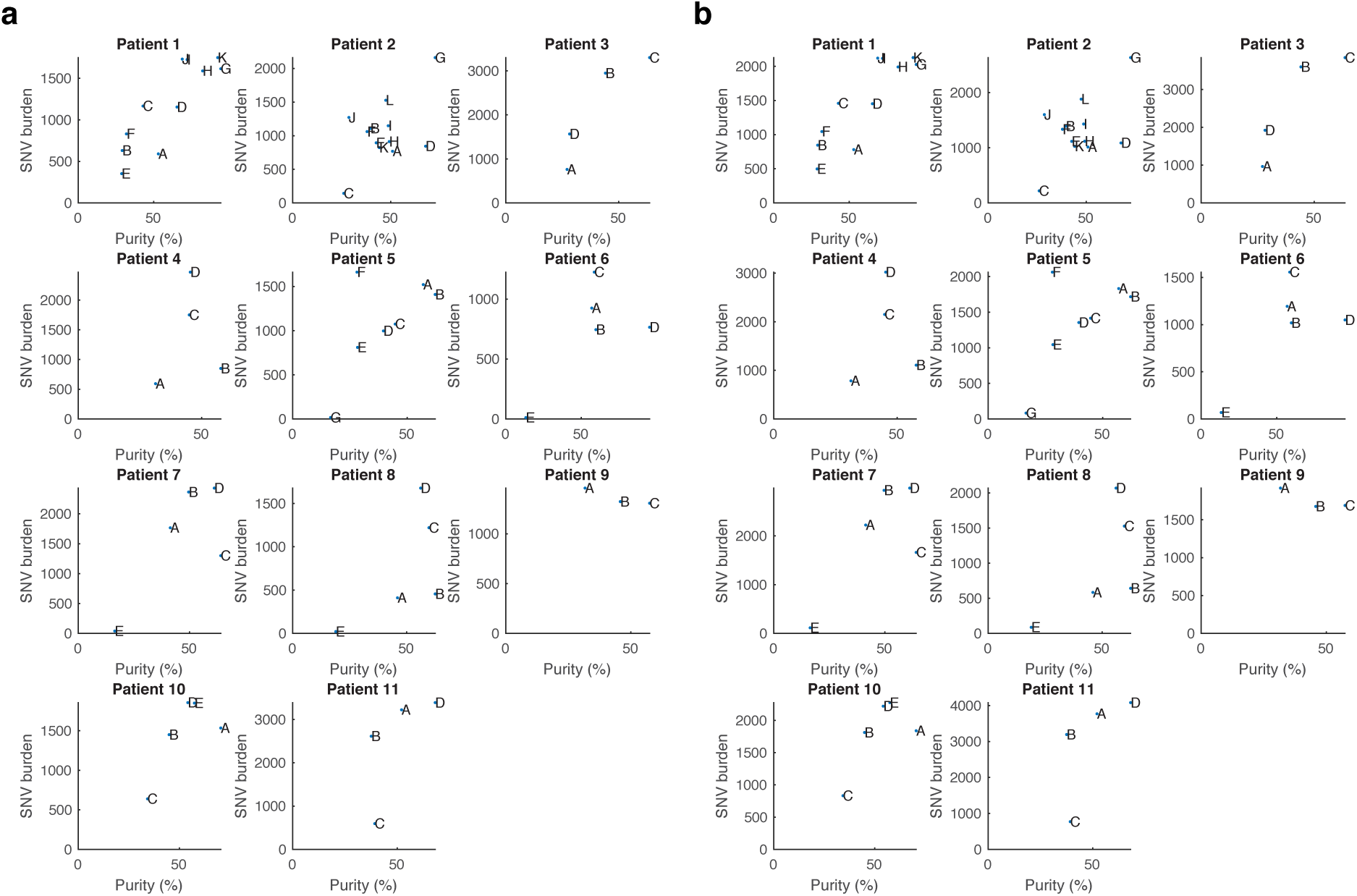
Positive correlations between per-sample estimated purity and average variant allele frequency (VAF). Tumor purity, as determined by PurBayes, correlated positively with the SNV burden in each patient. (**a**) Results using SNV calls used for phylogenetic analyses, which were stringently filtered for known population variants (dbSNP150 and ENSEMBL variants). (**b**) Results using relaxed population variant filtering (dbSNP138), as used for driver mutations and mutational signature analyses, were nearly identical.

**Supplementary Figure 14.**
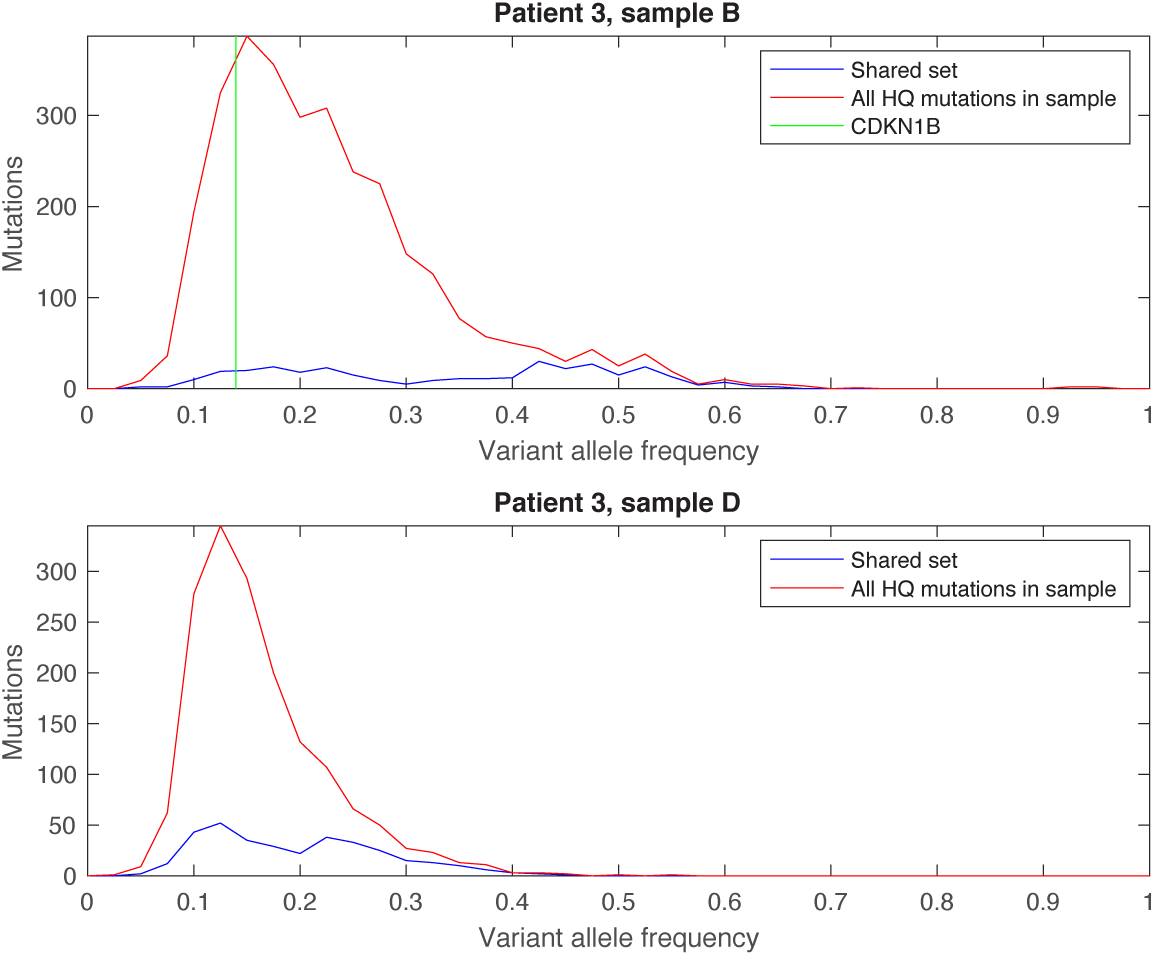
Variant allele frequency (VAF)-based analysis of a *CDKN1B* mutation in Patient 3. A *CDKN1B* SNV present in the metastatic primary tumor B is one of a large number of private variants (not shared with the metastasis D) present at relatively low VAF compared to shared variants, which showed higher mean VAF and bimodality in the primary tumor B. This is indicative of subclonality, early whole genome duplication, or a combination of both.

**Supplementary Figure 15.**
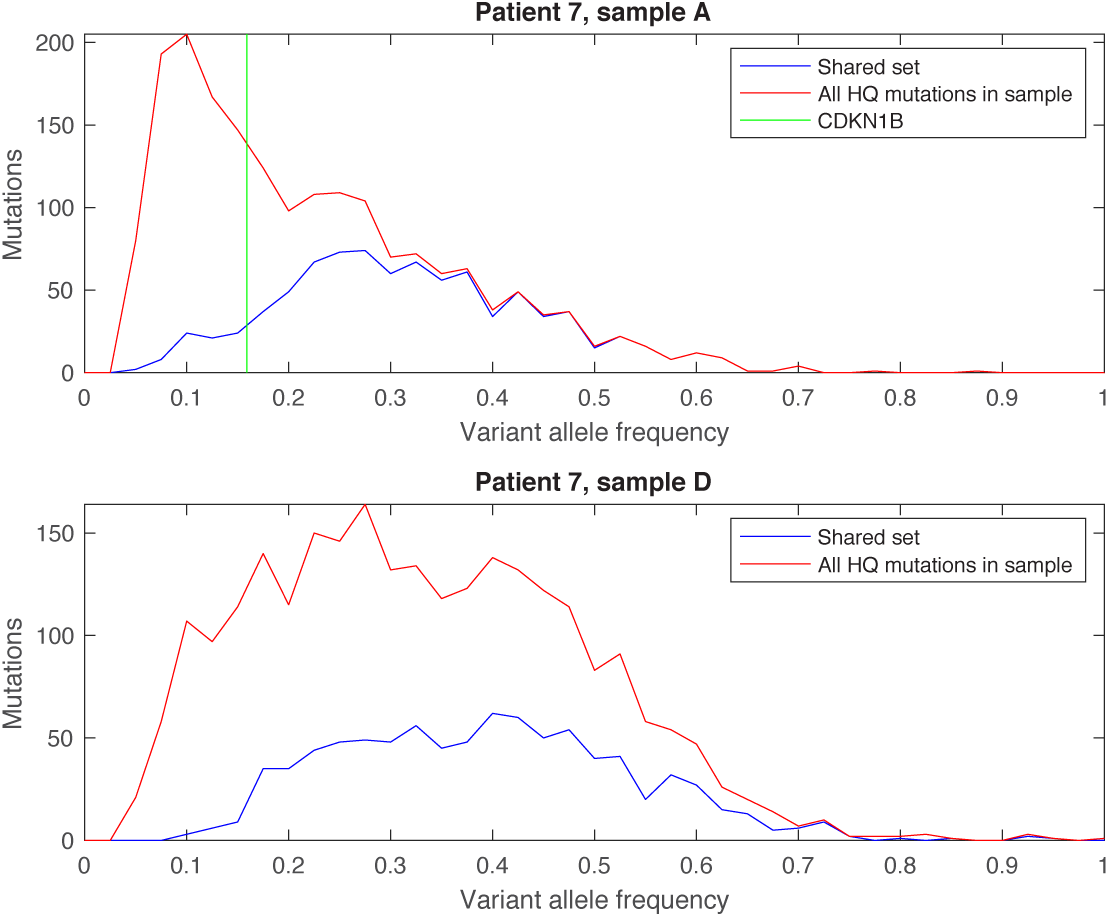
Variant allele frequency (VAF)-based analysis of a *CDKN1B* mutation in Patient 7. A *CDKN1B* SNV present in the metastatic primary tumor A is one of a many private variants (not shared with the metastasis D) present at relatively low VAF compared to shared variants.

**Supplementary Figure 16.**
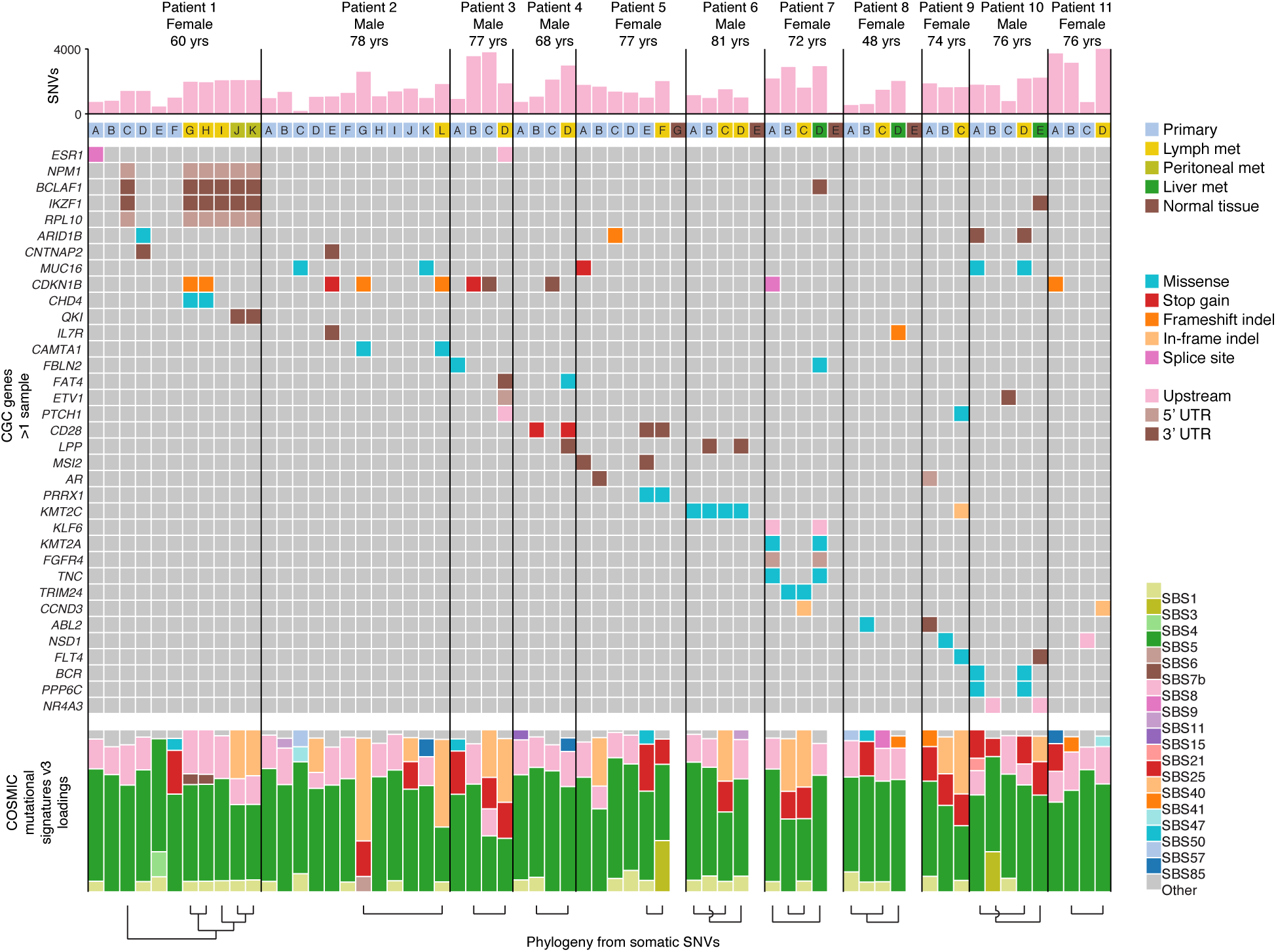
Overview of potential driver mutations including regulatory (upstream and UTR) mutations for all 11 patients. All CGC genes with non-synonymous mutations in more than one sample are shown. Upstream and UTR mutations are shown as reported by AnnoVar. Phylogenetic relationships previously inferred from somatic SNVs (main **Fig. 1-2**) are indicated at the bottom. Somatic SNV burdens, potential driver mutations (Cancer Gene Census genes mutated in >1 sample) and mutational signature loadings in this figure were determined using less rigorous population variant filtering compared to the phylogenetic analyses (see Materials and Methods). CGC, Cancer Gene Census.

**Supplementary Figure 17.**
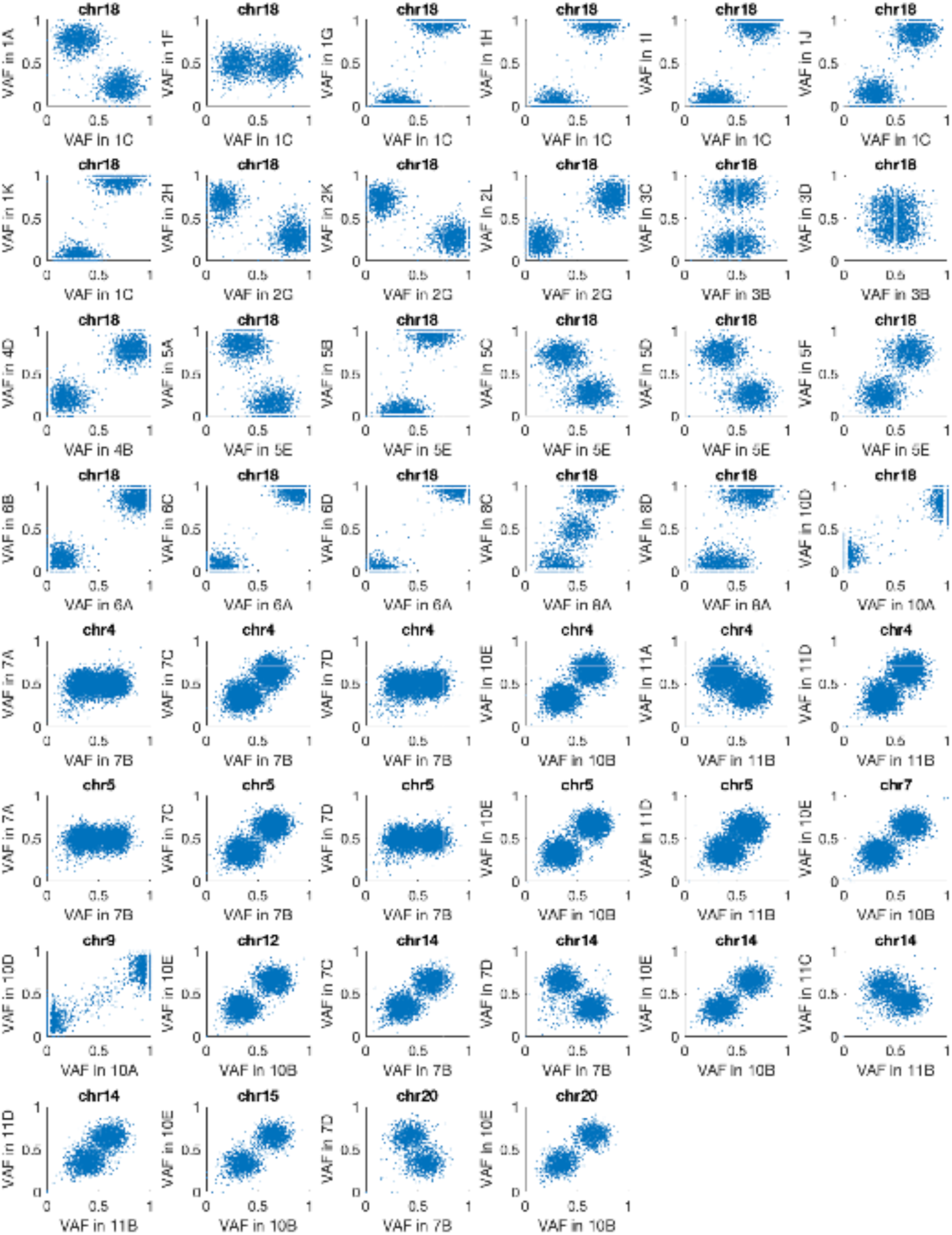
Chromosomal phasing of whole-chromosome copy number events based on germline heterozygous SNPs. Heterozygous germline SNPs on chr18 (read coverage 20 and VAF ranging from 0.25 to 0.75 in the blood normal) were used to determine what chromosome homolog was altered. The scatter plots compare VAFs for individual SNPs in a reference sample compared to other relevant samples. The data was down-sampled to 20% to reduce dot density in the plots. The related samples 7A and 7D lacked VAF bimodality on chr4/5 indicative of biallelic gain, further supported by high copy number amplitude in these samples. Lack of bimodality on chr18 in the related samples 3B and 3D may be explained by a biallelic deletion on a whole-genome duplication background (notably the unrelated sample, 3C, deviates and shows bimodality). Alternative explanations include a homozygous deletion (unlikely due to low-level amplitude change in 3D) or a false copy number call (unlikely due to strong amplitude change in 3B).

**Supplementary Figure 18.**
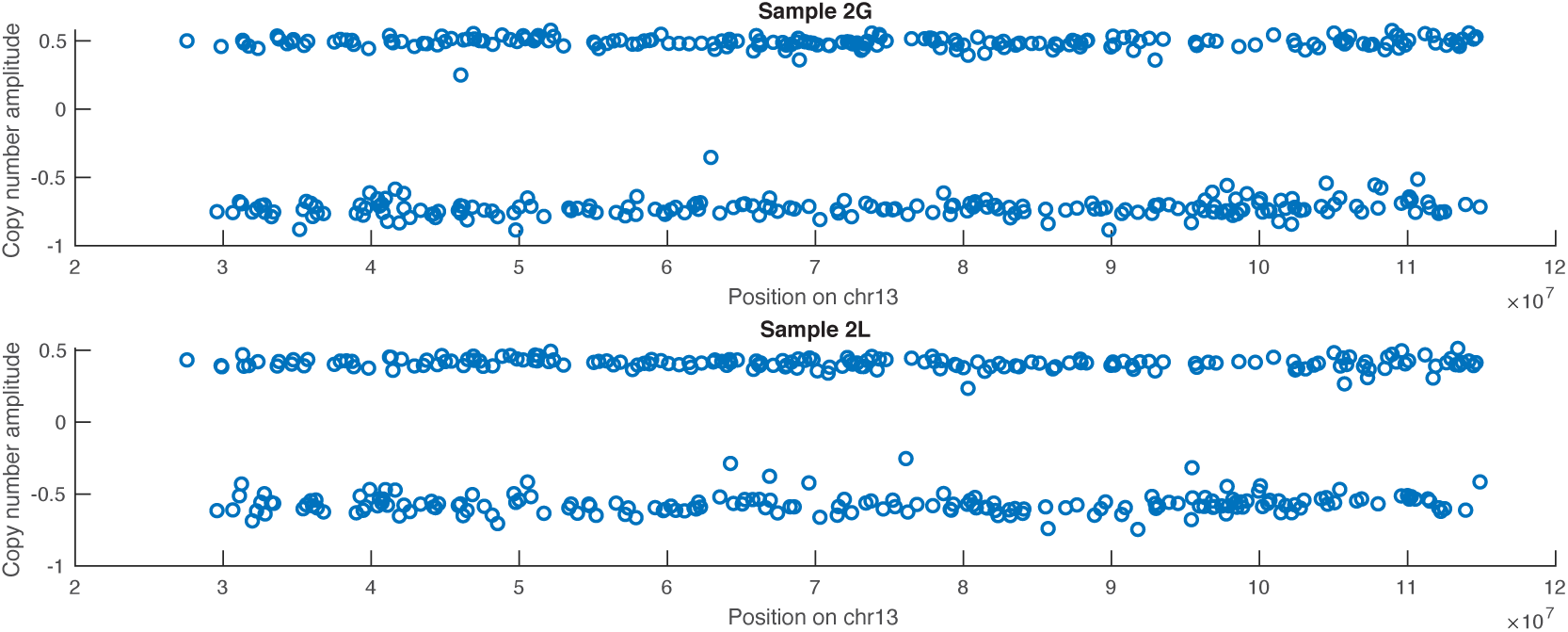
Clustered copy number alterations on chromosome 13 in Patient 2 compatible with chromothripsis. CNAs on chromosome 13 in Patient 2 (samples G and L) oscillated between two distinct amplitude states, which is a hallmark of chromothripsis.

**Supplementary Figure 19.**
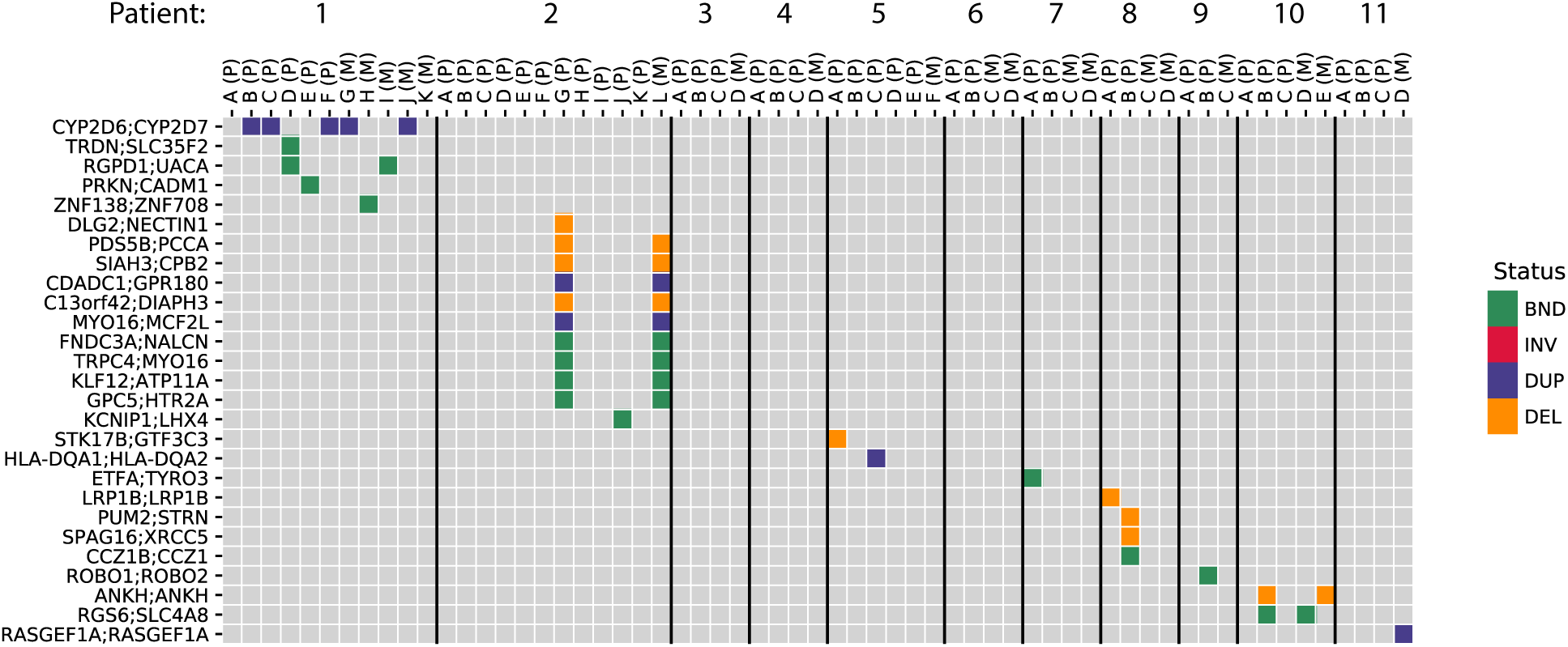
Analysis of somatic genomic structural alterations. In-frame fusion gene pairs colored by alteration type, with translocations (BND) shown in green, inversions in red, duplications in purple, and deletions in orange.

**Table S1.**
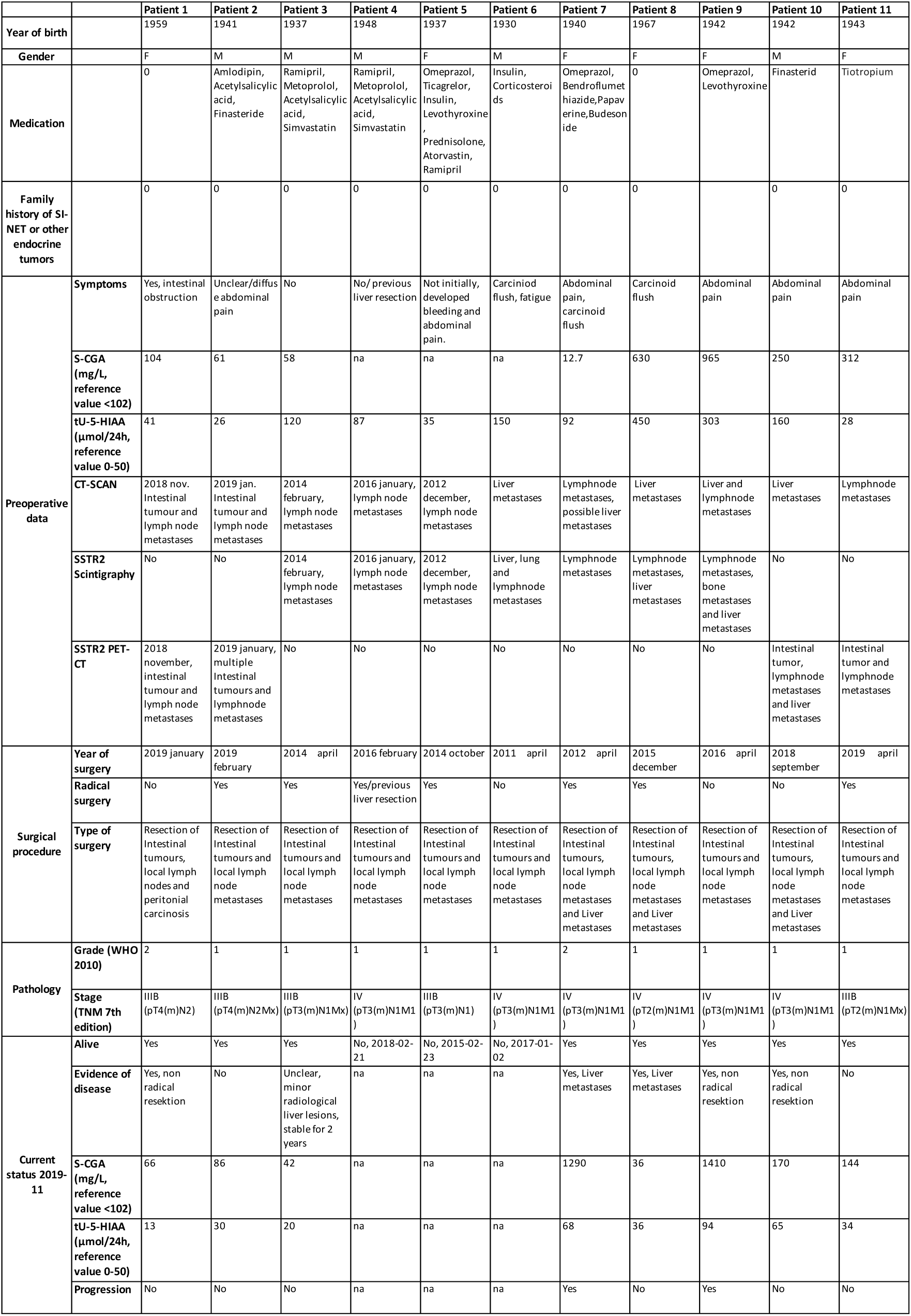
Patient characteristics.

**Table S2.**
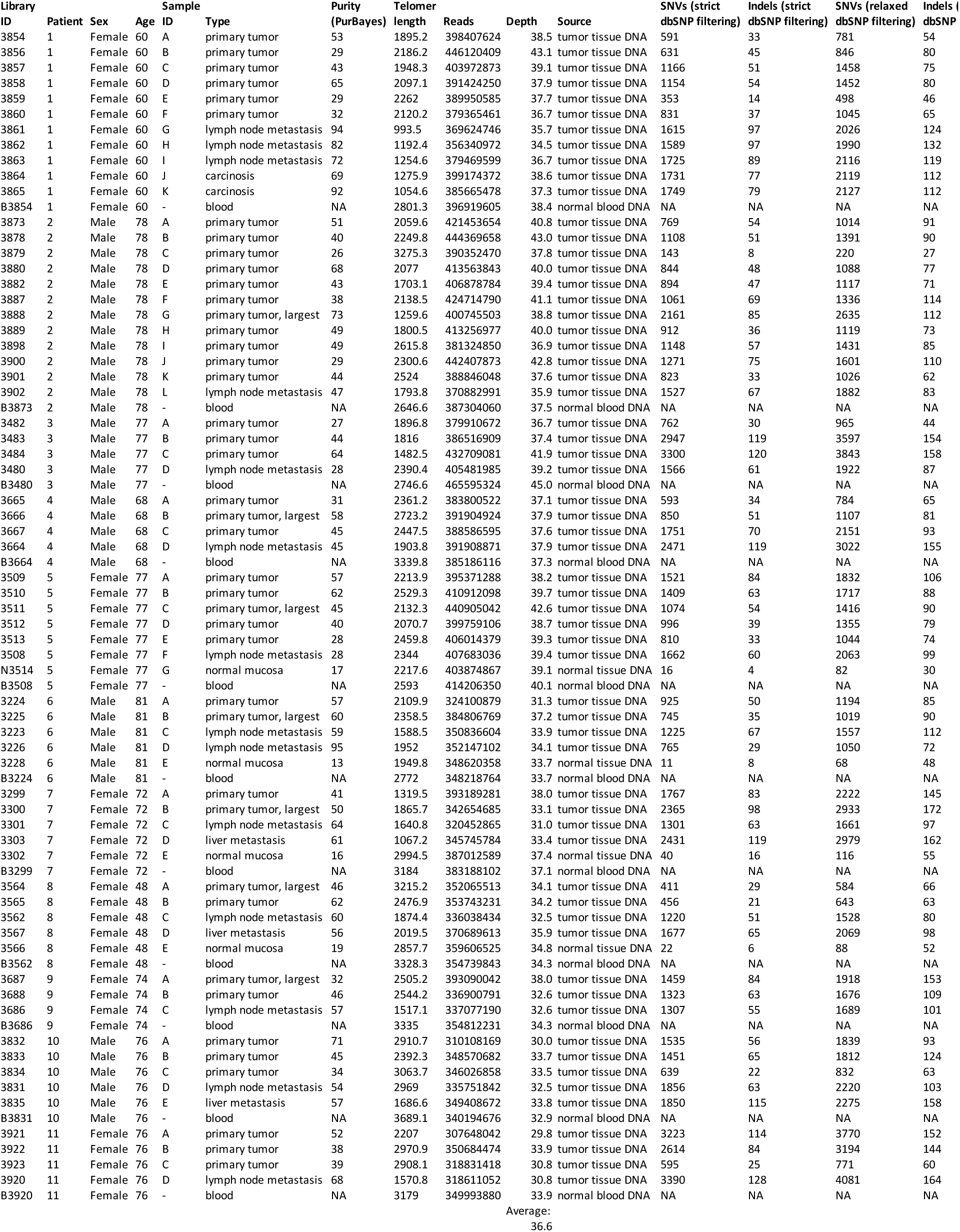
Sample overview and WGS statistics.

Sample overview and WGS statistics
| Library ID | Patient | Sex | Age | Sample ID | Type | Purity (PurBayes) | Telomer length | Reads | Depth | Source | SNVs (strict dbSNP filtering) | Indels (strict dbSNP filtering) | SNVs (relaxed dbSNP filtering) | Indels (relaxed dbSNP filtering) |
| --- | --- | --- | --- | --- | --- | --- | --- | --- | --- | --- | --- | --- | --- | --- |
| 3854 | 1 | Female | 60 | A | primary tumor | 53 | 1895.2 | 398407624 | 38.5 | tumor tissue DNA | 591 | 33 | 781 | 54 |
| 3856 | 1 | Female | 60 | B | primary tumor | 29 | 2186.2 | 446120409 | 43.1 | tumor tissue DNA | 631 | 45 | 846 | 80 |
| 3857 | 1 | Female | 60 | C | primary tumor | 43 | 1948.3 | 403972873 | 39.1 | tumor tissue DNA | 1166 | 51 | 1458 | 75 |
| 3858 | 1 | Female | 60 | D | primary tumor | 65 | 2097.1 | 391424250 | 37.9 | tumor tissue DNA | 1154 | 54 | 1452 | 80 |
| 3859 | 1 | Female | 60 | E | primary tumor | 29 | 2262 | 389950585 | 37.7 | tumor tissue DNA | 353 | 14 | 498 | 46 |
| 3860 | 1 | Female | 60 | F | primary tumor | 32 | 2120.2 | 379365461 | 36.7 | tumor tissue DNA | 831 | 37 | 1045 | 65 |
| 3861 | 1 | Female | 60 | G | lymph node metastasis | 94 | 993.5 | 369624746 | 35.7 | tumor tissue DNA | 1615 | 97 | 2026 | 124 |
| 3862 | 1 | Female | 60 | H | lymph node metastasis | 82 | 1192.4 | 356340972 | 34.5 | tumor tissue DNA | 1589 | 97 | 1990 | 132 |
| 3863 | 1 | Female | 60 | I | lymph node metastasis | 72 | 1254.6 | 379469599 | 36.7 | tumor tissue DNA | 1725 | 89 | 2116 | 119 |
| 3864 | 1 | Female | 60 | J | carcinosis | 69 | 1275.9 | 399174372 | 38.6 | tumor tissue DNA | 1731 | 77 | 2119 | 112 |
| 3865 | 1 | Female | 60 | K | carcinosis | 92 | 1054.6 | 385665478 | 37.3 | tumor tissue DNA | 1749 | 79 | 2127 | 112 |
| B3854 | 1 | Female | 60 | - | blood | NA | 2801.3 | 396919605 | 38.4 | normal blood DNA | NA | NA | NA | NA |
| 3873 | 2 | Male | 78 | A | primary tumor | 51 | 2059.6 | 421453654 | 40.8 | tumor tissue DNA | 769 | 54 | 1014 | 91 |
| 3878 | 2 | Male | 78 | B | primary tumor | 40 | 2249.8 | 444369658 | 43.0 | tumor tissue DNA | 1108 | 51 | 1391 | 90 |
| 3879 | 2 | Male | 78 | C | primary tumor | 26 | 3275.3 | 390352470 | 37.8 | tumor tissue DNA | 143 | 8 | 220 | 27 |
| 3880 | 2 | Male | 78 | D | primary tumor | 68 | 2077 | 413563843 | 40.0 | tumor tissue DNA | 844 | 48 | 1088 | 77 |
| 3882 | 2 | Male | 78 | E | primary tumor | 43 | 1703.1 | 406878784 | 39.4 | tumor tissue DNA | 894 | 47 | 1117 | 71 |
| 3887 | 2 | Male | 78 | F | primary tumor | 38 | 2138.5 | 424714790 | 41.1 | tumor tissue DNA | 1061 | 69 | 1336 | 114 |
| 3888 | 2 | Male | 78 | G | primary tumor, largest | 73 | 1259.6 | 400745503 | 38.8 | tumor tissue DNA | 2161 | 85 | 2635 | 112 |
| 3889 | 2 | Male | 78 | H | primary tumor | 49 | 1800.5 | 413256977 | 40.0 | tumor tissue DNA | 912 | 36 | 1119 | 73 |
| 3898 | 2 | Male | 78 | I | primary tumor | 49 | 2615.8 | 381324850 | 36.9 | tumor tissue DNA | 1148 | 57 | 1431 | 85 |
| 3900 | 2 | Male | 78 | J | primary tumor | 29 | 2300.6 | 442407873 | 42.8 | tumor tissue DNA | 1271 | 75 | 1601 | 110 |
| 3901 | 2 | Male | 78 | K | primary tumor | 44 | 2524 | 388846048 | 37.6 | tumor tissue DNA | 823 | 33 | 1026 | 62 |
| 3902 | 2 | Male | 78 | L | lymph node metastasis | 47 | 1793.8 | 370882991 | 35.9 | tumor tissue DNA | 1527 | 67 | 1882 | 83 |
| B3873 | 2 | Male | 78 | - | blood | NA | 2646.6 | 387304060 | 37.5 | normal blood DNA | NA | NA | NA | NA |
| 3482 | 3 | Male | 77 | A | primary tumor | 27 | 1896.8 | 379910672 | 36.7 | tumor tissue DNA | 762 | 30 | 965 | 44 |
| 3483 | 3 | Male | 77 | B | primary tumor | 44 | 1816 | 386516909 | 37.4 | tumor tissue DNA | 2947 | 119 | 3597 | 154 |
| 3484 | 3 | Male | 77 | C | primary tumor | 64 | 1482.5 | 432709081 | 41.9 | tumor tissue DNA | 3300 | 120 | 3843 | 158 |
| 3480 | 3 | Male | 77 | D | lymph node metastasis | 28 | 2390.4 | 405481985 | 39.2 | tumor tissue DNA | 1566 | 61 | 1922 | 87 |
| B3480 | 3 | Male | 77 | - | blood | NA | 2746.6 | 465595324 | 45.0 | normal blood DNA | NA | NA | NA | NA |
| 3665 | 4 | Male | 68 | A | primary tumor | 31 | 2361.2 | 383800522 | 37.1 | tumor tissue DNA | 593 | 34 | 784 | 65 |
| 3666 | 4 | Male | 68 | B | primary tumor, largest | 58 | 2723.2 | 391904924 | 37.9 | tumor tissue DNA | 850 | 51 | 1107 | 81 |
| 3667 | 4 | Male | 68 | C | primary tumor | 45 | 2447.5 | 388586595 | 37.6 | tumor tissue DNA | 1751 | 70 | 2151 | 93 |
| 3664 | 4 | Male | 68 | D | lymph node metastasis | 45 | 1903.8 | 391908871 | 37.9 | tumor tissue DNA | 2471 | 119 | 3022 | 155 |
| B3664 | 4 | Male | 68 | - | blood | NA | 3339.8 | 385186116 | 37.3 | normal blood DNA | NA | NA | NA | NA |
| 3509 | 5 | Female | 77 | A | primary tumor | 57 | 2213.9 | 395371288 | 38.2 | tumor tissue DNA | 1521 | 84 | 1832 | 106 |
| 3510 | 5 | Female | 77 | B | primary tumor | 62 | 2529.3 | 410912098 | 39.7 | tumor tissue DNA | 1409 | 63 | 1717 | 88 |
| 3511 | 5 | Female | 77 | C | primary tumor, largest | 45 | 2132.3 | 440905042 | 42.6 | tumor tissue DNA | 1074 | 54 | 1416 | 90 |
| 3512 | 5 | Female | 77 | D | primary tumor | 40 | 2070.7 | 399759106 | 38.7 | tumor tissue DNA | 996 | 39 | 1355 | 79 |
| 3513 | 5 | Female | 77 | E | primary tumor | 28 | 2459.8 | 406014379 | 39.3 | tumor tissue DNA | 810 | 33 | 1044 | 74 |
| 3508 | 5 | Female | 77 | F | lymph node metastasis | 28 | 2344 | 407683036 | 39.4 | tumor tissue DNA | 1662 | 60 | 2063 | 99 |
| N3514 | 5 | Female | 77 | G | normal mucosa | 17 | 2217.6 | 403874867 | 39.1 | normal tissue DNA | 16 | 4 | 82 | 30 |
| B3508 | 5 | Female | 77 | - | blood | NA | 2593 | 414206350 | 40.1 | normal blood DNA | NA | NA | NA | NA |
| 3224 | 6 | Male | 81 | A | primary tumor | 57 | 2109.9 | 324100879 | 31.3 | tumor tissue DNA | 925 | 50 | 1194 | 85 |
| 3225 | 6 | Male | 81 | B | primary tumor, largest | 60 | 2358.5 | 384806769 | 37.2 | tumor tissue DNA | 745 | 35 | 1019 | 90 |
| 3223 | 6 | Male | 81 | C | lymph node metastasis | 59 | 1588.5 | 350836604 | 33.9 | tumor tissue DNA | 1225 | 67 | 1557 | 112 |
| 3226 | 6 | Male | 81 | D | lymph node metastasis | 95 | 1952 | 352147102 | 34.1 | tumor tissue DNA | 765 | 29 | 1050 | 72 |
| 3228 | 6 | Male | 81 | E | normal mucosa | 13 | 1949.8 | 348620358 | 33.7 | normal tissue DNA | 11 | 8 | 68 | 48 |
| B3224 | 6 | Male | 81 | - | blood | NA | 2772 | 348218764 | 33.7 | normal blood DNA | NA | NA | NA | NA |
| 3299 | 7 | Female | 72 | A | primary tumor | 41 | 1319.5 | 393189281 | 38.0 | tumor tissue DNA | 1767 | 83 | 2222 | 145 |
| 3300 | 7 | Female | 72 | B | primary tumor, largest | 50 | 1865.7 | 342654685 | 33.1 | tumor tissue DNA | 2365 | 98 | 2933 | 172 |
| 3301 | 7 | Female | 72 | C | lymph node metastasis | 64 | 1640.8 | 320452865 | 31.0 | tumor tissue DNA | 1301 | 63 | 1661 | 97 |
| 3303 | 7 | Female | 72 | D | liver metastasis | 61 | 1067.2 | 345745784 | 33.4 | tumor tissue DNA | 2431 | 119 | 2979 | 162 |
| 3302 | 7 | Female | 72 | E | normal mucosa | 16 | 2994.5 | 387012589 | 37.4 | normal tissue DNA | 40 | 16 | 116 | 55 |
| B3299 | 7 | Female | 72 | - | blood | NA | 3184 | 383188102 | 37.1 | normal blood DNA | NA | NA | NA | NA |
| 3564 | 8 | Female | 48 | A | primary tumor, largest | 46 | 3215.2 | 352065513 | 34.1 | tumor tissue DNA | 411 | 29 | 584 | 66 |
| 3565 | 8 | Female | 48 | B | primary tumor | 62 | 2476.9 | 353743231 | 34.2 | tumor tissue DNA | 456 | 21 | 643 | 63 |
| 3562 | 8 | Female | 48 | C | lymph node metastasis | 60 | 1874.4 | 336038434 | 32.5 | tumor tissue DNA | 1220 | 51 | 1528 | 80 |
| 3567 | 8 | Female | 48 | D | liver metastasis | 56 | 2019.5 | 370689613 | 35.9 | tumor tissue DNA | 1677 | 65 | 2069 | 98 |
| 3566 | 8 | Female | 48 | E | normal mucosa | 19 | 2857.7 | 359606525 | 34.8 | normal tissue DNA | 22 | 6 | 88 | 52 |
| B3562 | 8 | Female | 48 | - | blood | NA | 3328.3 | 354739843 | 34.3 | normal blood DNA | NA | NA | NA | NA |
| 3687 | 9 | Female | 74 | A | primary tumor, largest | 32 | 2505.2 | 393090042 | 38.0 | tumor tissue DNA | 1459 | 84 | 1918 | 153 |
| 3688 | 9 | Female | 74 | B | primary tumor | 46 | 2544.2 | 336900791 | 32.6 | tumor tissue DNA | 1323 | 63 | 1676 | 109 |
| 3686 | 9 | Female | 74 | C | lymph node metastasis | 57 | 1517.1 | 337077190 | 32.6 | tumor tissue DNA | 1307 | 55 | 1689 | 101 |
| B3686 | 9 | Female | 74 | - | blood | NA | 3335 | 354812231 | 34.3 | normal blood DNA | NA | NA | NA | NA |
| 3832 | 10 | Male | 76 | A | primary tumor | 71 | 2910.7 | 310108169 | 30.0 | tumor tissue DNA | 1535 | 56 | 1839 | 93 |
| 3833 | 10 | Male | 76 | B | primary tumor | 45 | 2392.3 | 348570682 | 33.7 | tumor tissue DNA | 1451 | 65 | 1812 | 124 |
| 3834 | 10 | Male | 76 | C | primary tumor | 34 | 3063.7 | 346026858 | 33.5 | tumor tissue DNA | 639 | 22 | 832 | 63 |
| 3831 | 10 | Male | 76 | D | lymph node metastasis | 54 | 2969 | 335751842 | 32.5 | tumor tissue DNA | 1856 | 63 | 2220 | 103 |
| 3835 | 10 | Male | 76 | E | liver metastasis | 57 | 1686.6 | 349408672 | 33.8 | tumor tissue DNA | 1850 | 115 | 2275 | 158 |
| B3831 | 10 | Male | 76 | - | blood | NA | 3689.1 | 340194676 | 32.9 | normal blood DNA | NA | NA | NA | NA |
| 3921 | 11 | Female | 76 | A | primary tumor | 52 | 2207 | 307648042 | 29.8 | tumor tissue DNA | 3223 | 114 | 3770 | 152 |
| 3922 | 11 | Female | 76 | B | primary tumor | 38 | 2970.9 | 350684474 | 33.9 | tumor tissue DNA | 2614 | 84 | 3194 | 144 |
| 3923 | 11 | Female | 76 | C | primary tumor | 39 | 2908.1 | 318831418 | 30.8 | tumor tissue DNA | 595 | 25 | 771 | 60 |
| 3920 | 11 | Female | 76 | D | lymph node metastasis | 68 | 1570.8 | 318611052 | 30.8 | tumor tissue DNA | 3390 | 128 | 4081 | 164 |
| B3920 | 11 | Female | 76 | - | blood | NA | 3179 | 349993880 | 33.9 | normal blood DNA | NA | NA | NA | NA |

Average: 36.6

**Table S3.**
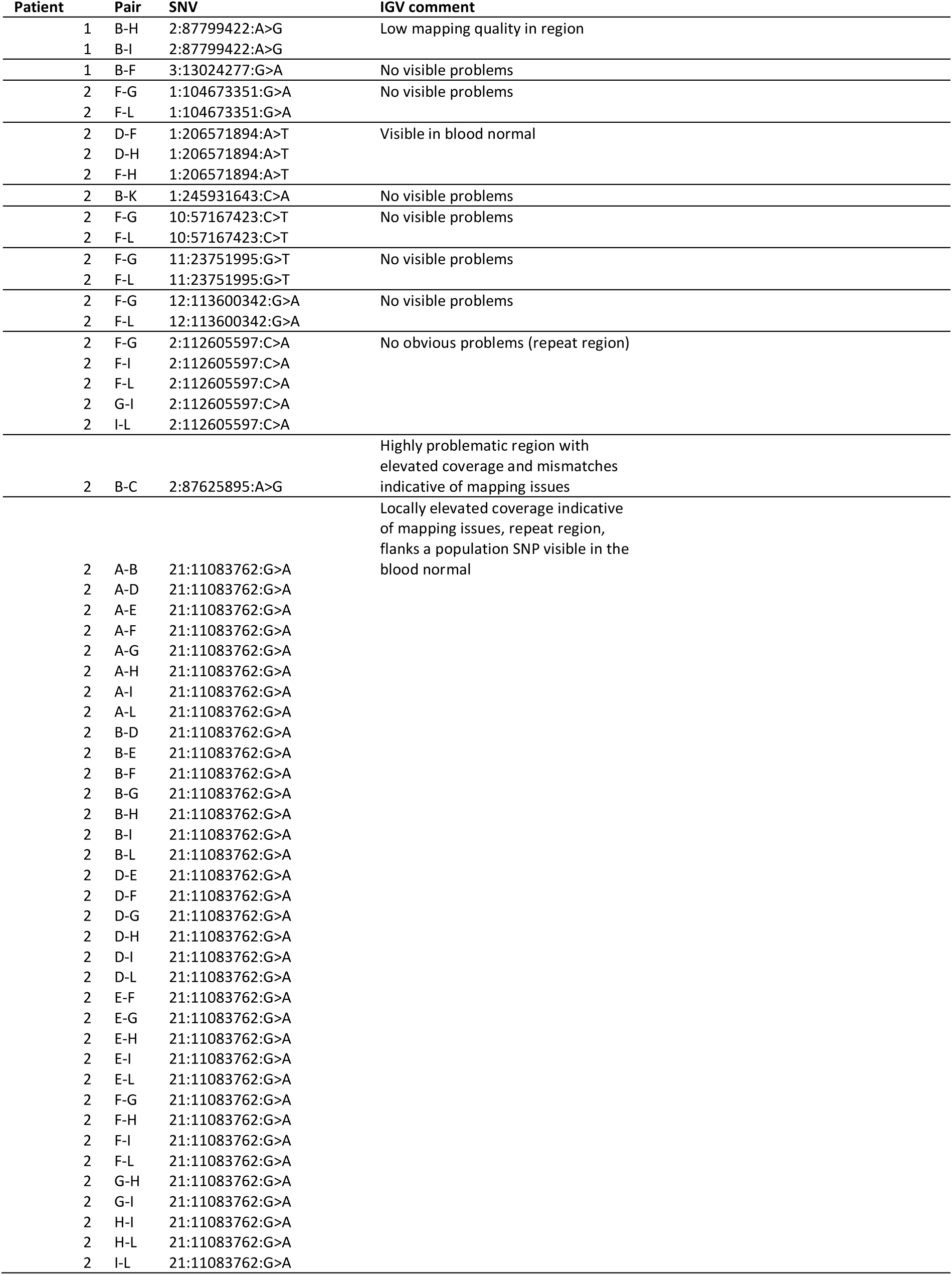
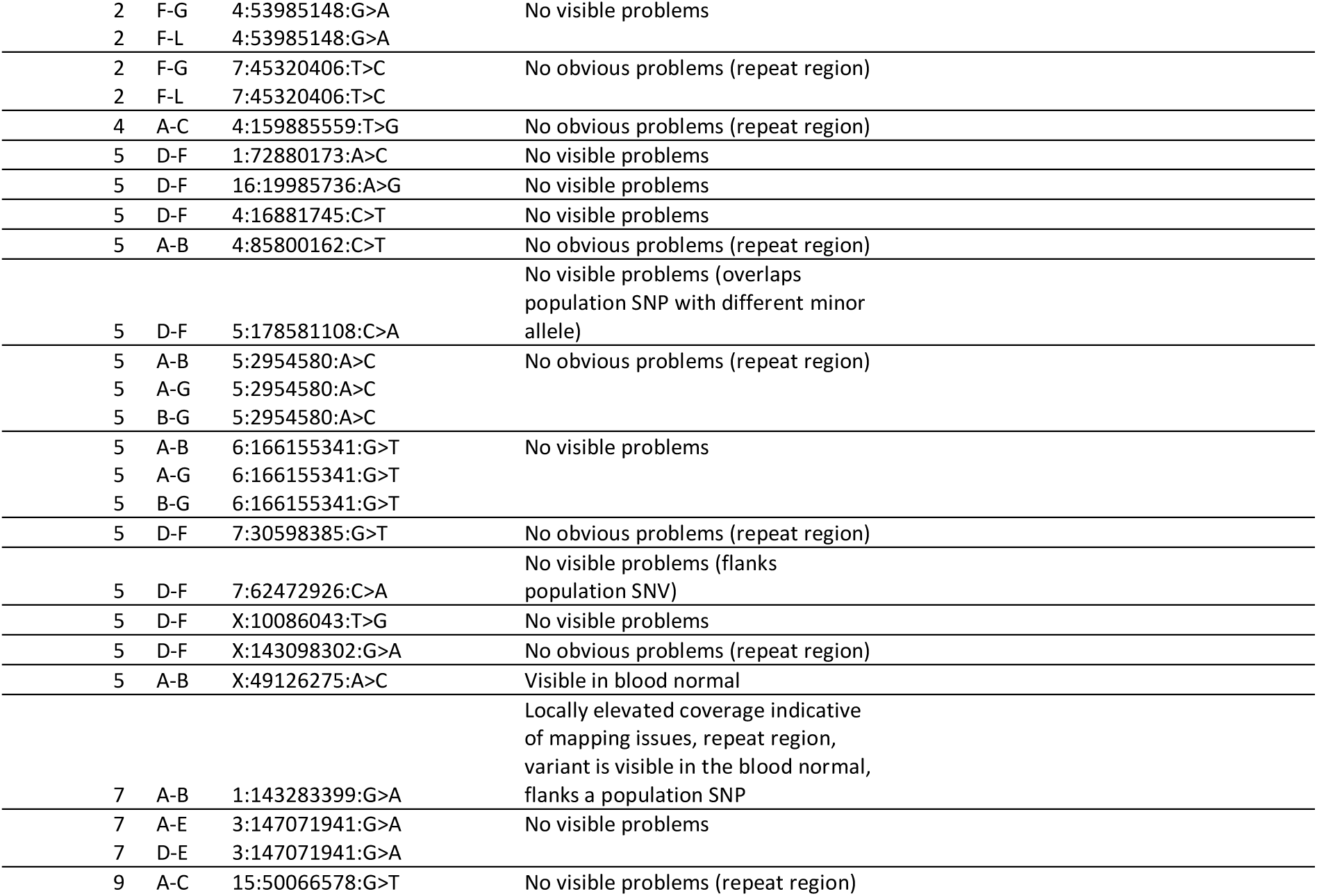
Manual inspection of 29 remaining spurious overlapping SNVs (appearing in pairs sharing <10 mutations) in high-confidence mutation calls.

**Table S4.**
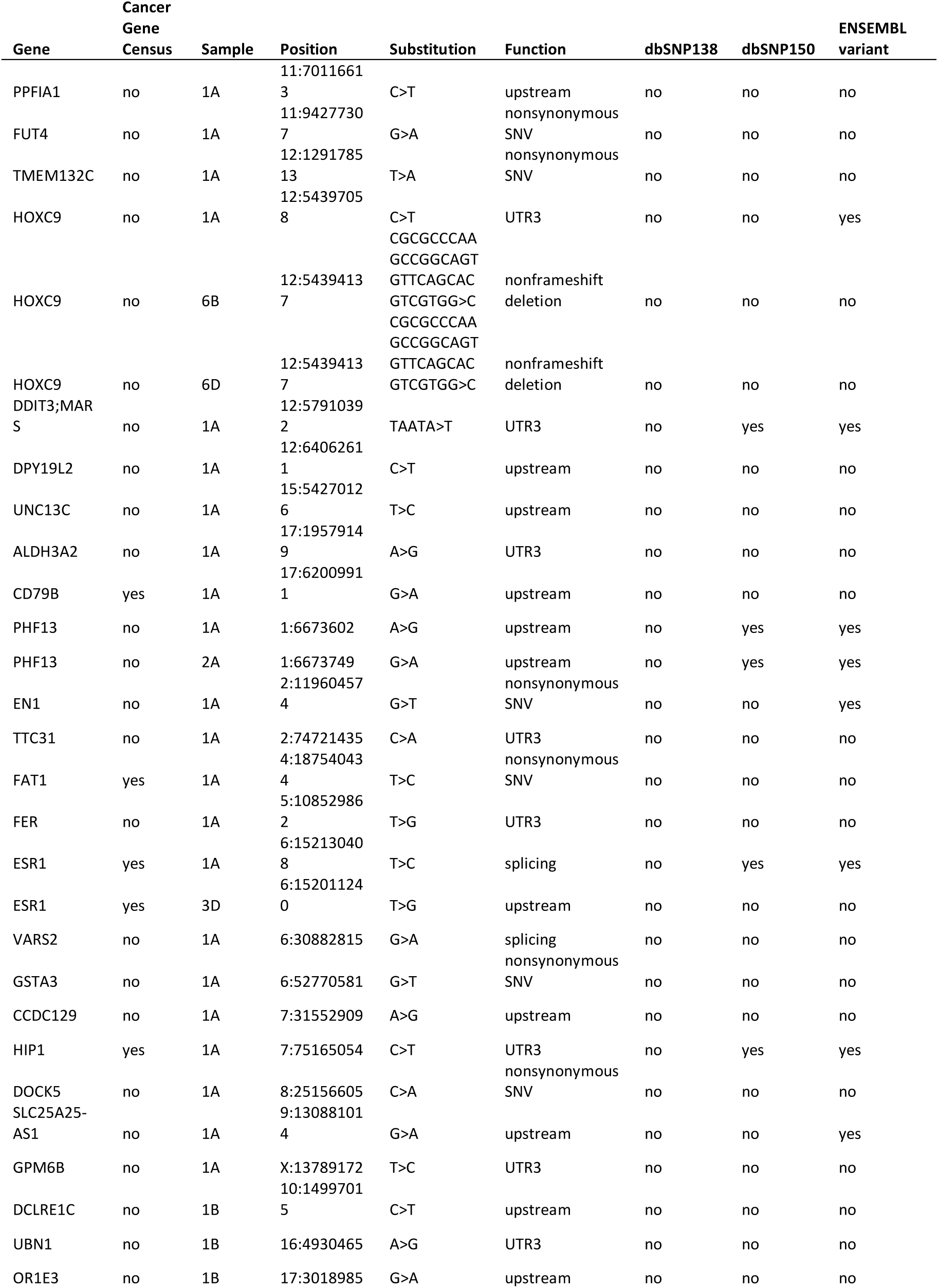
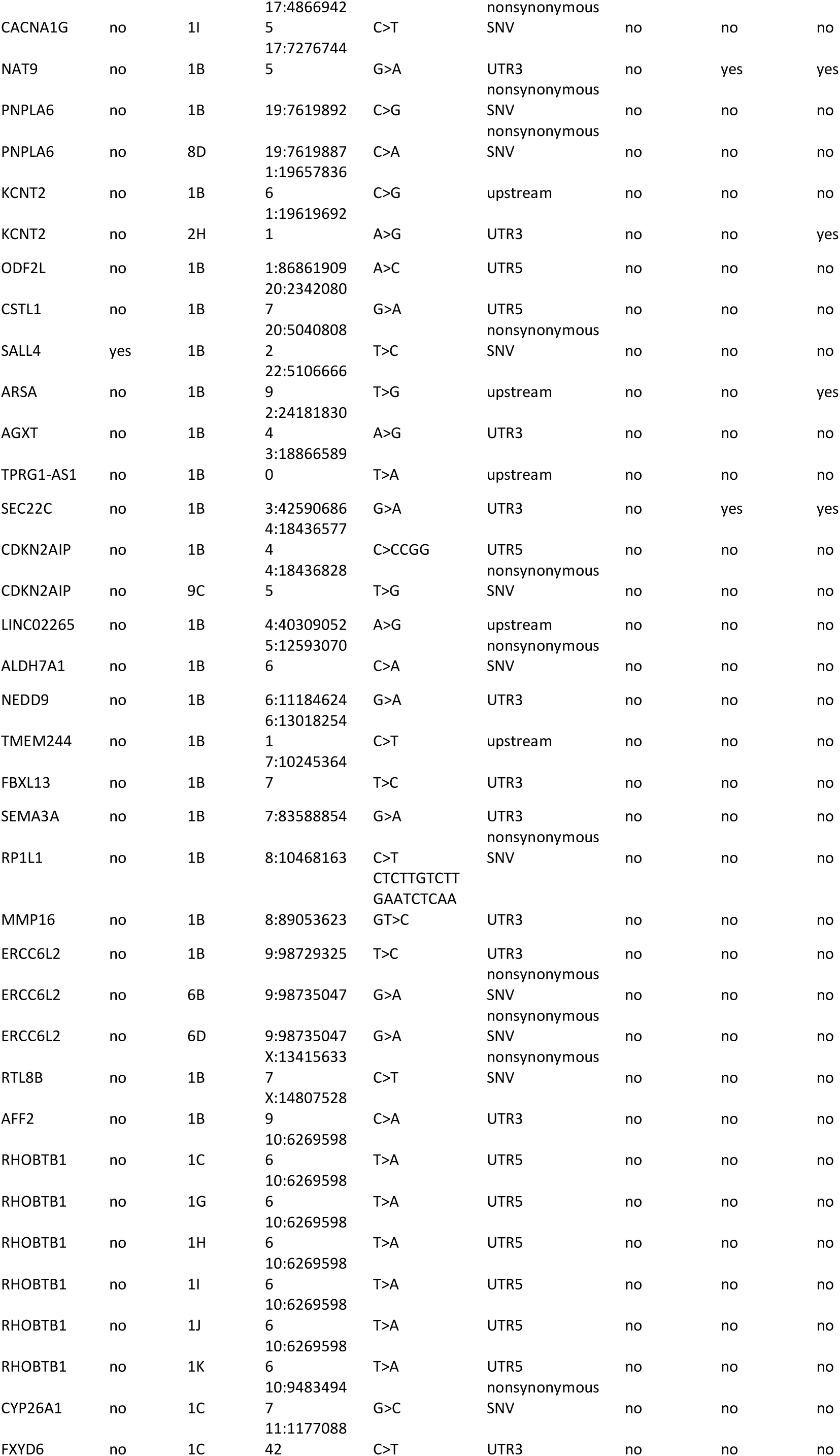
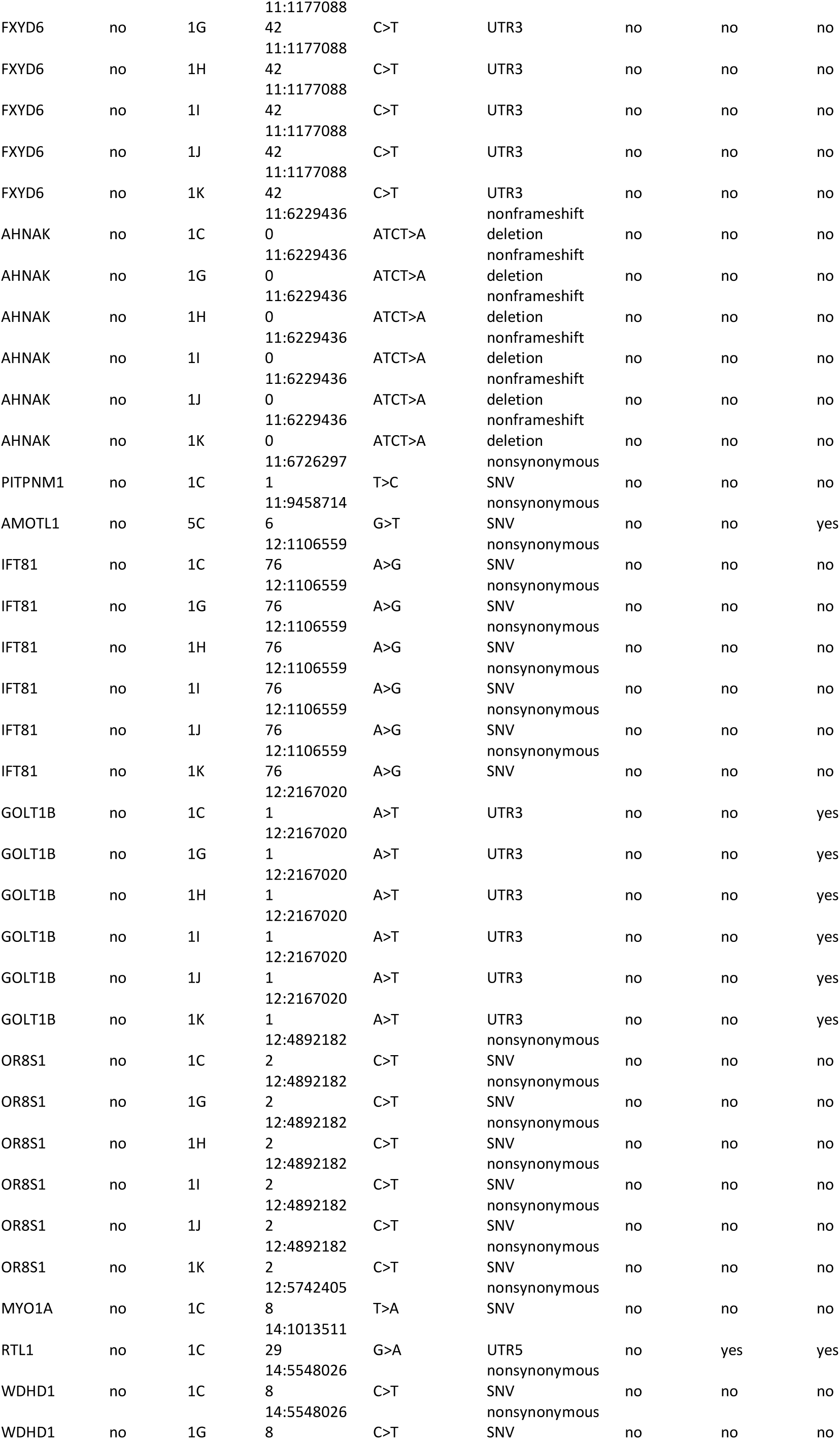
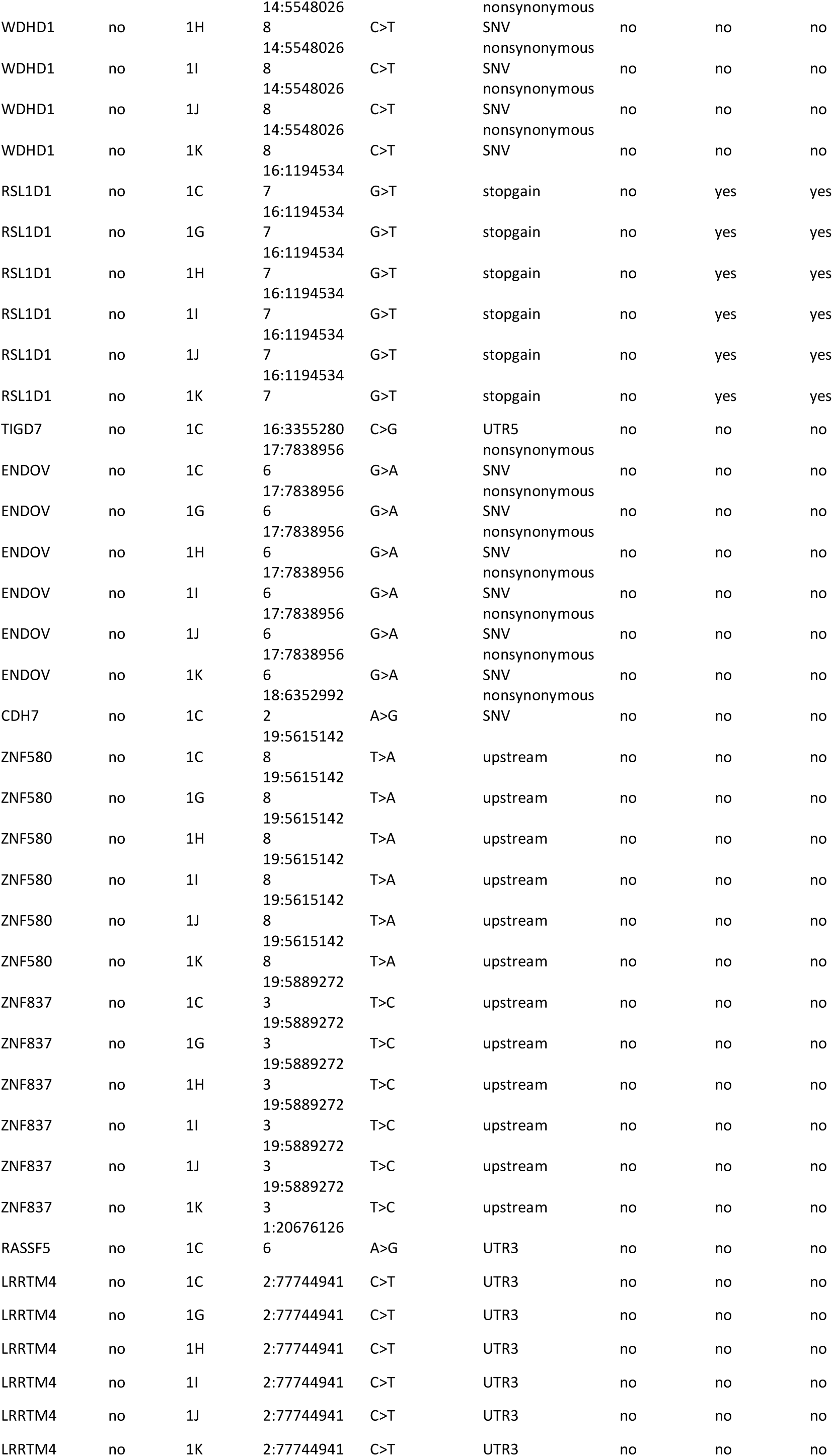
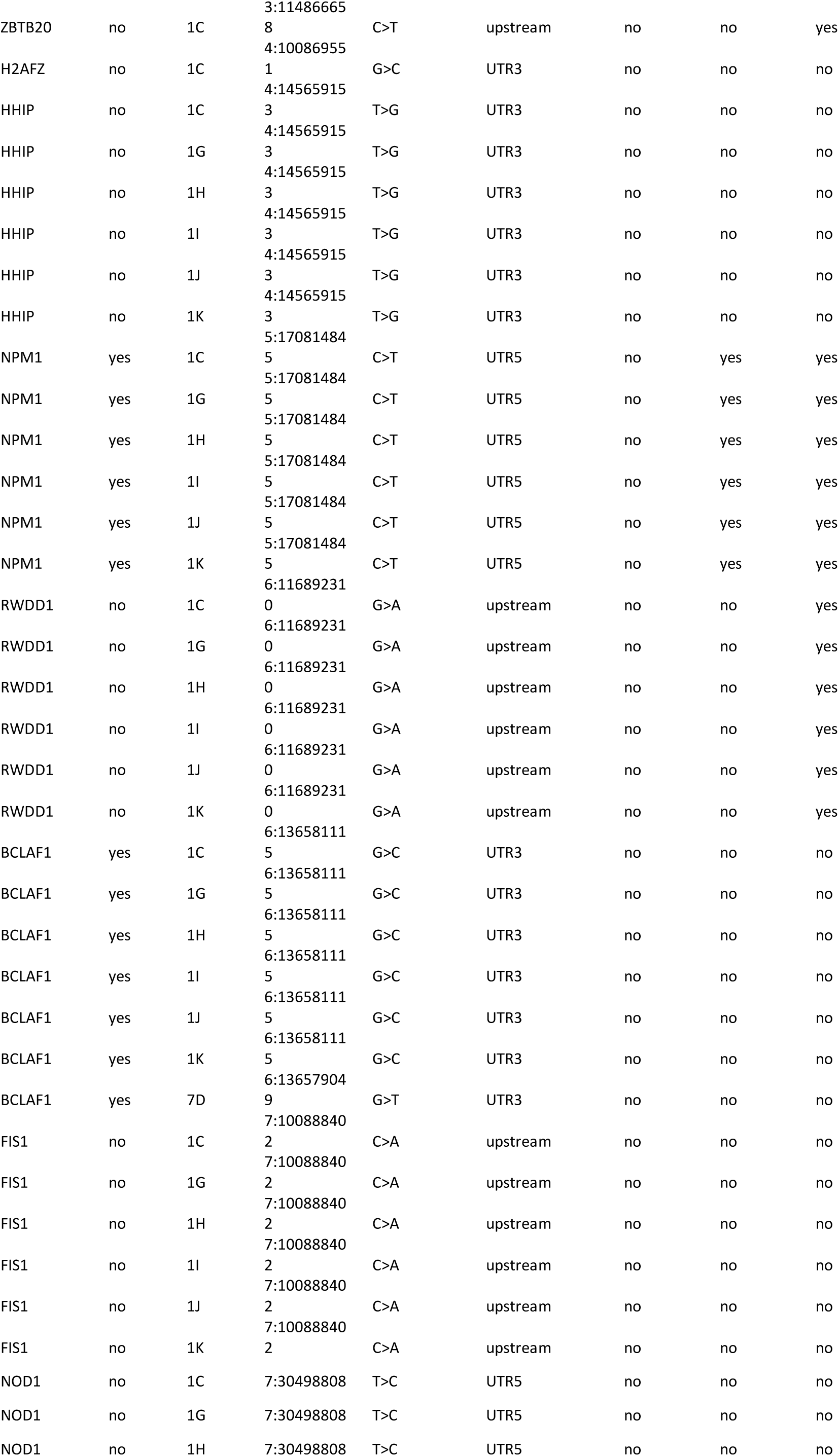
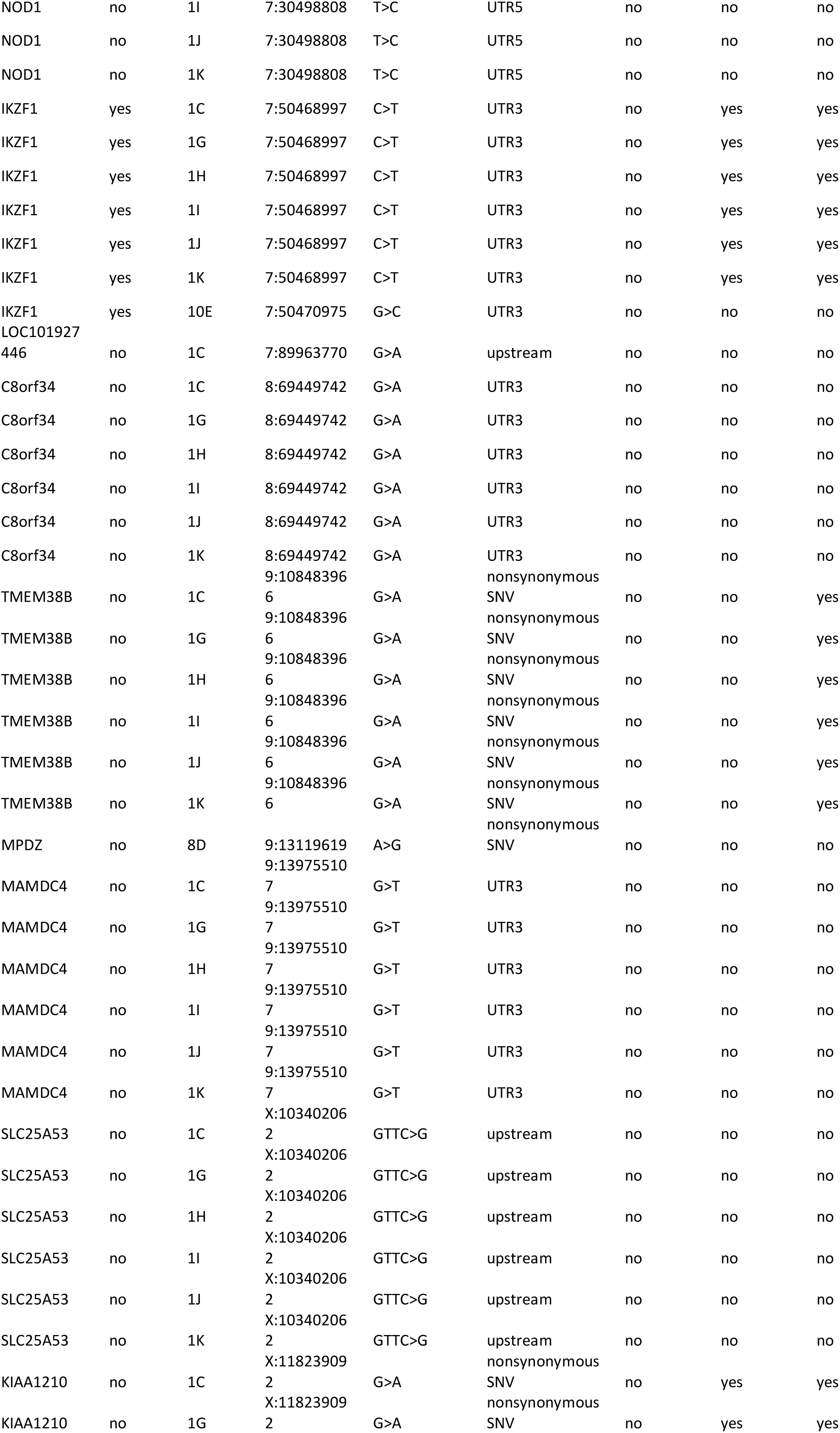
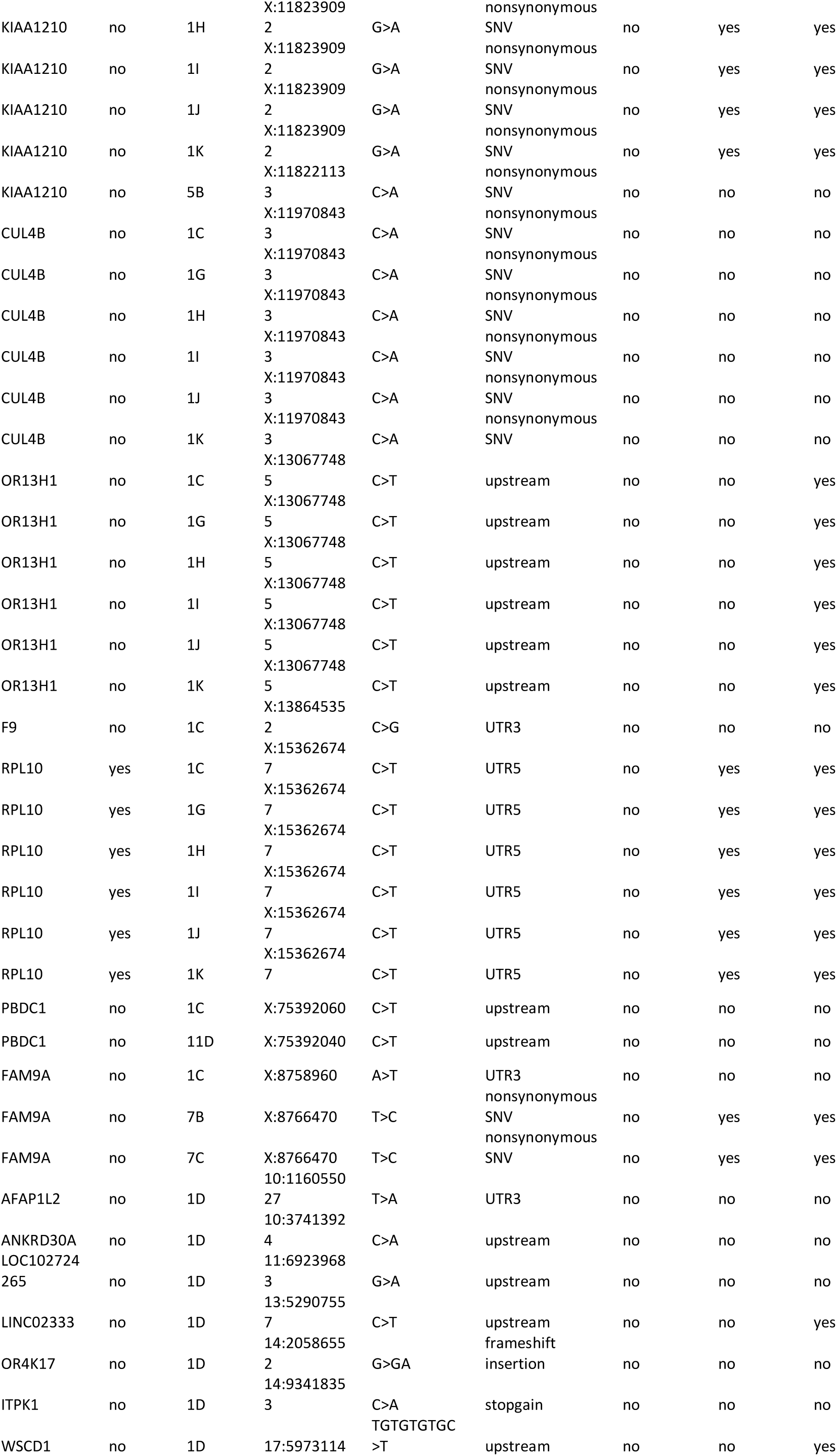
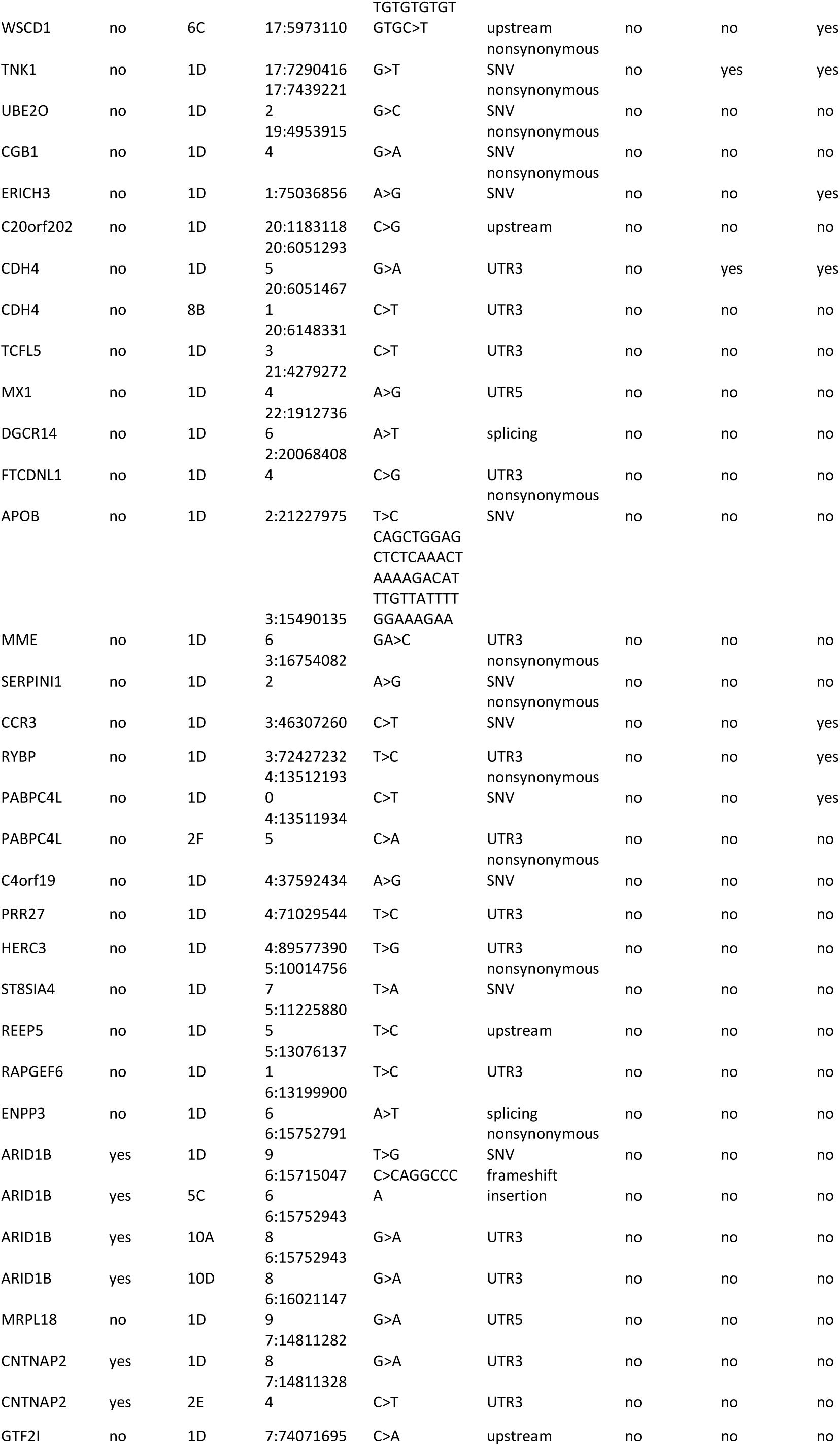
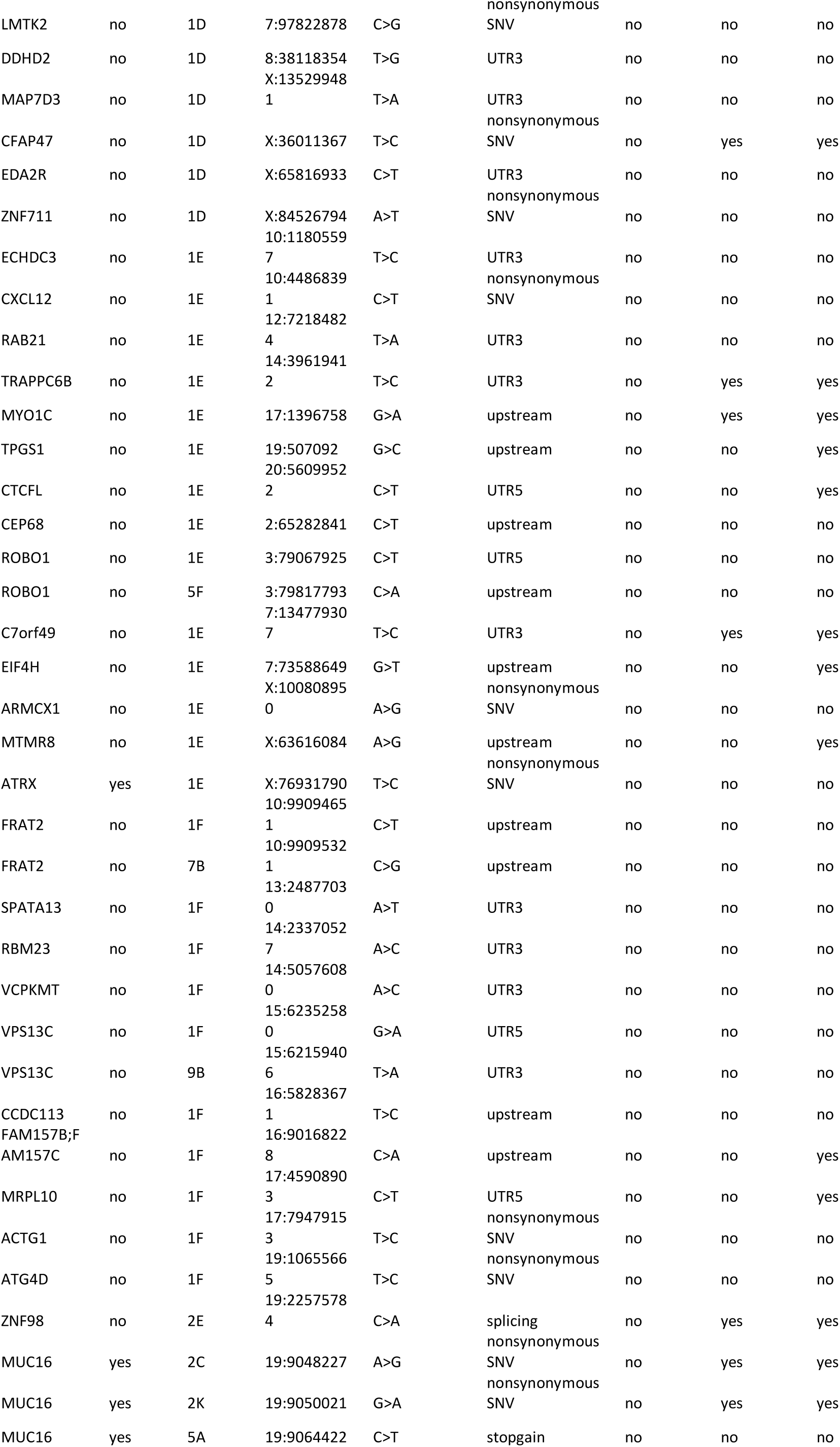
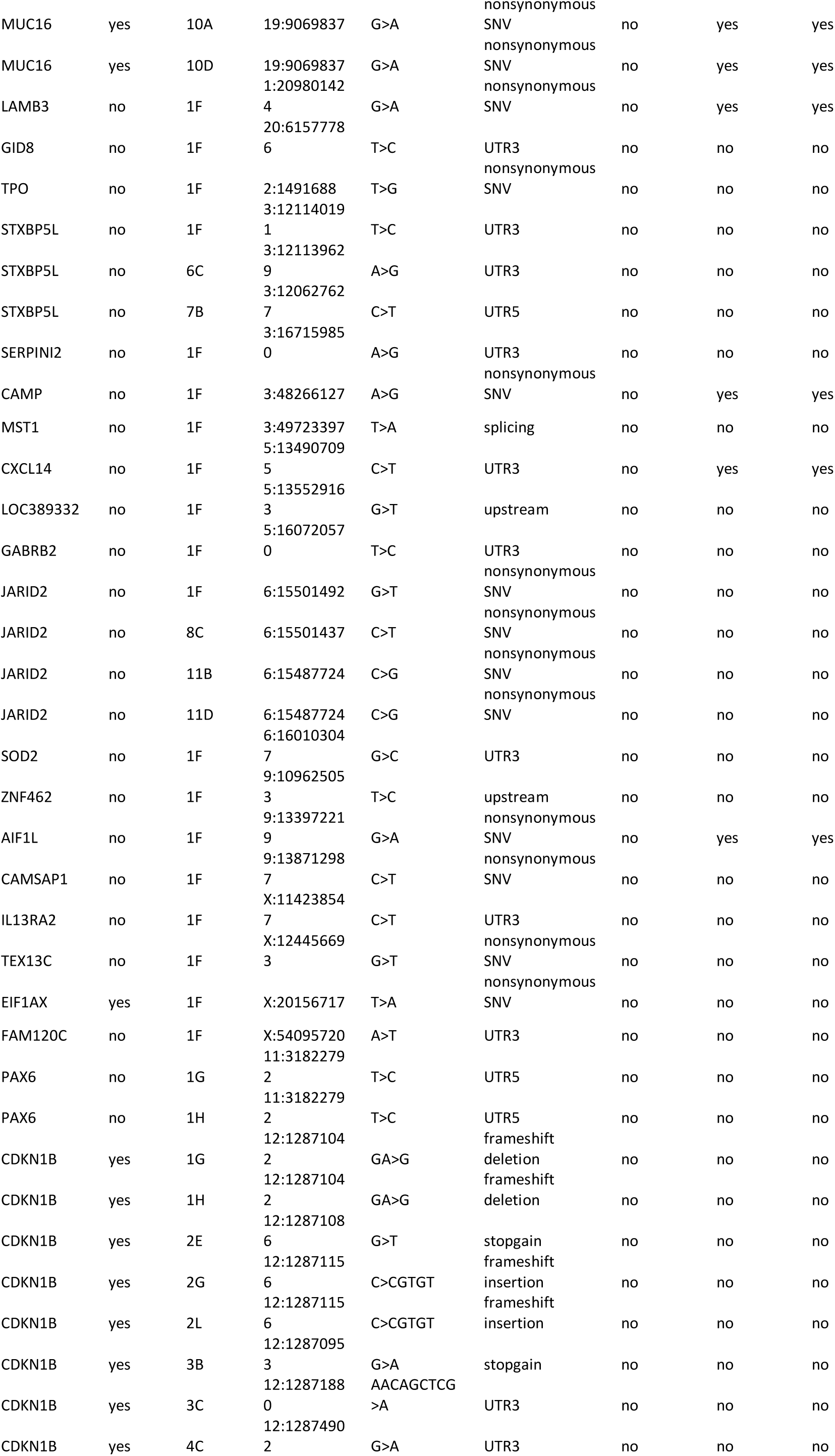
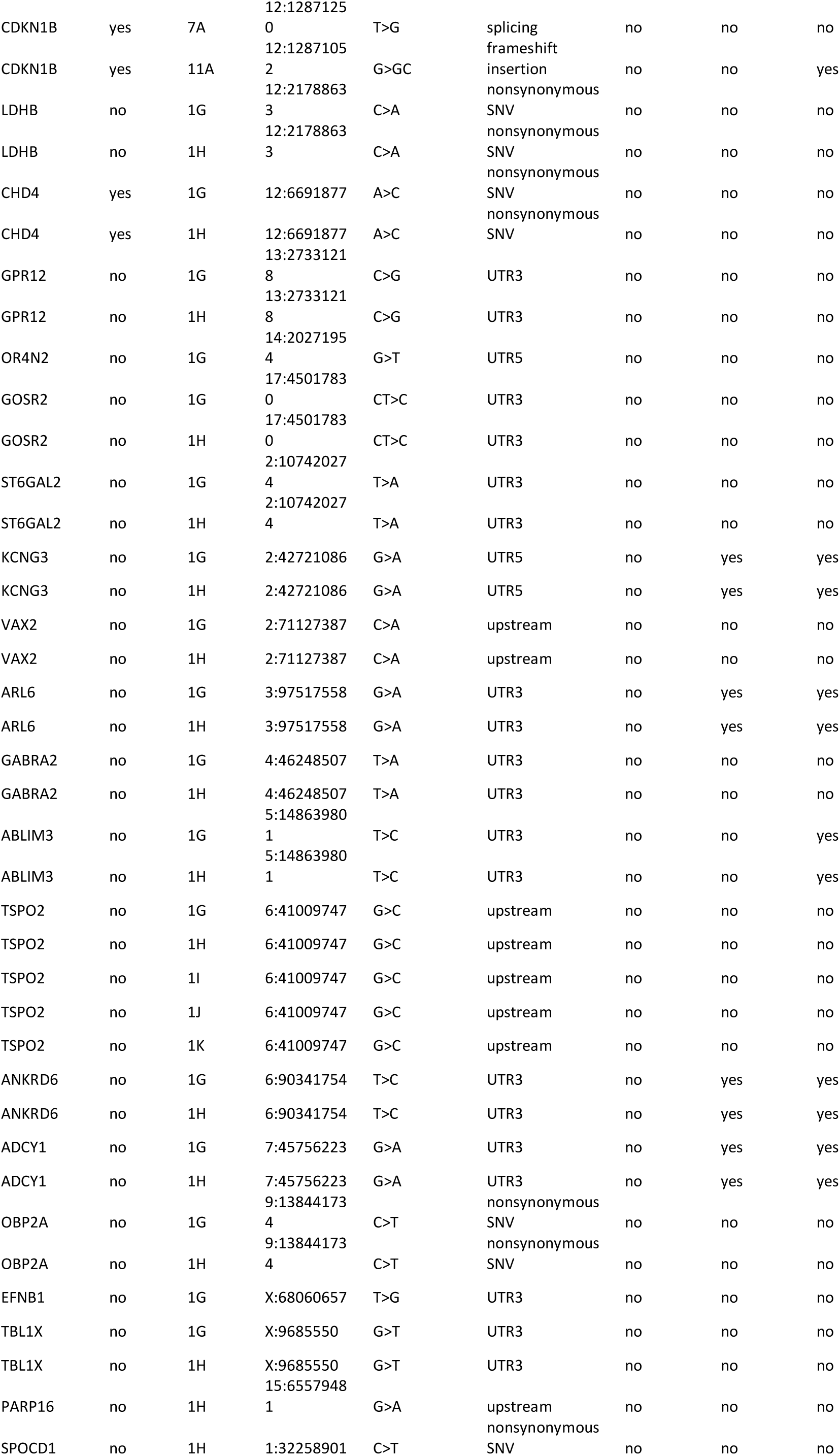
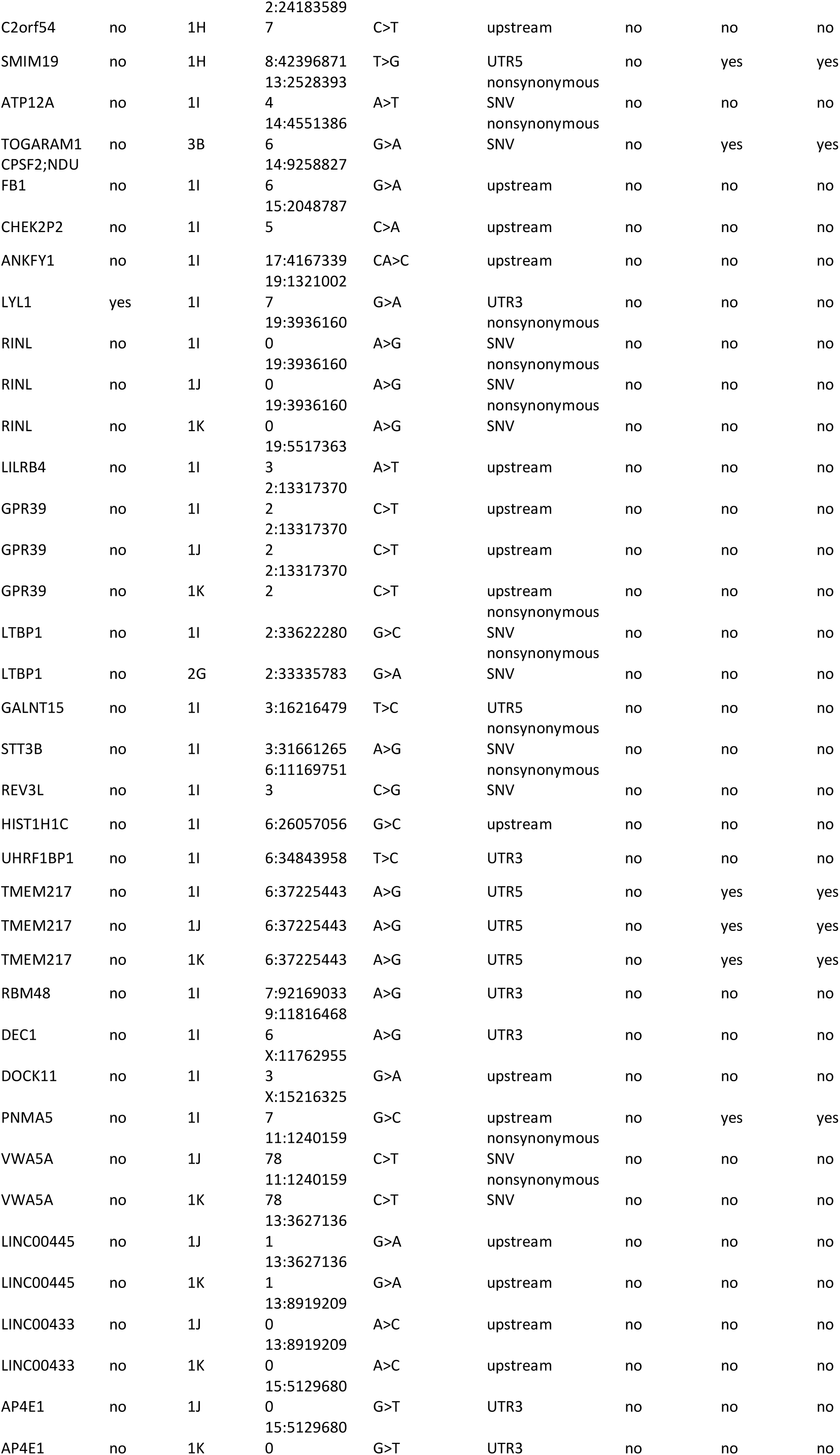
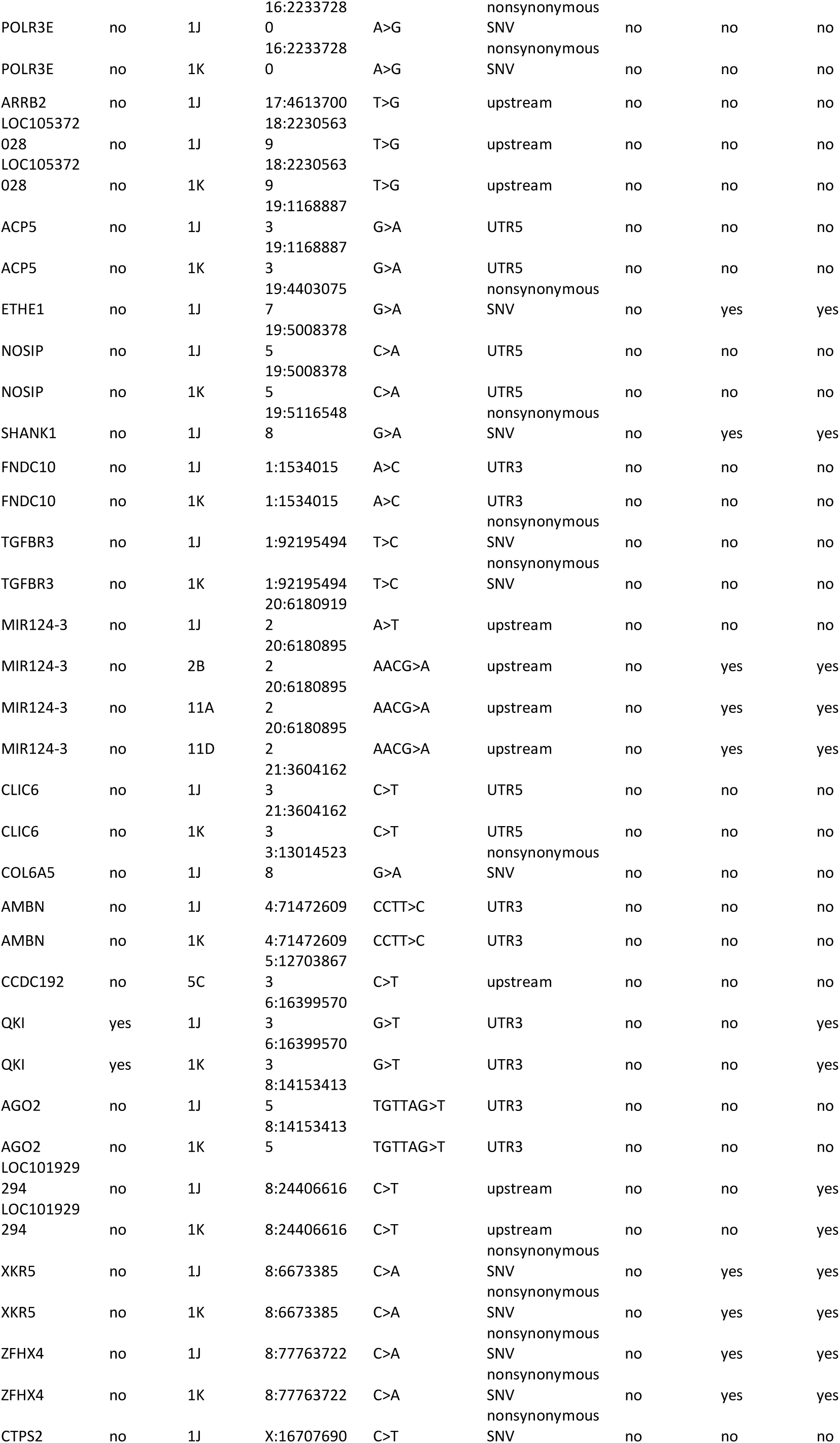
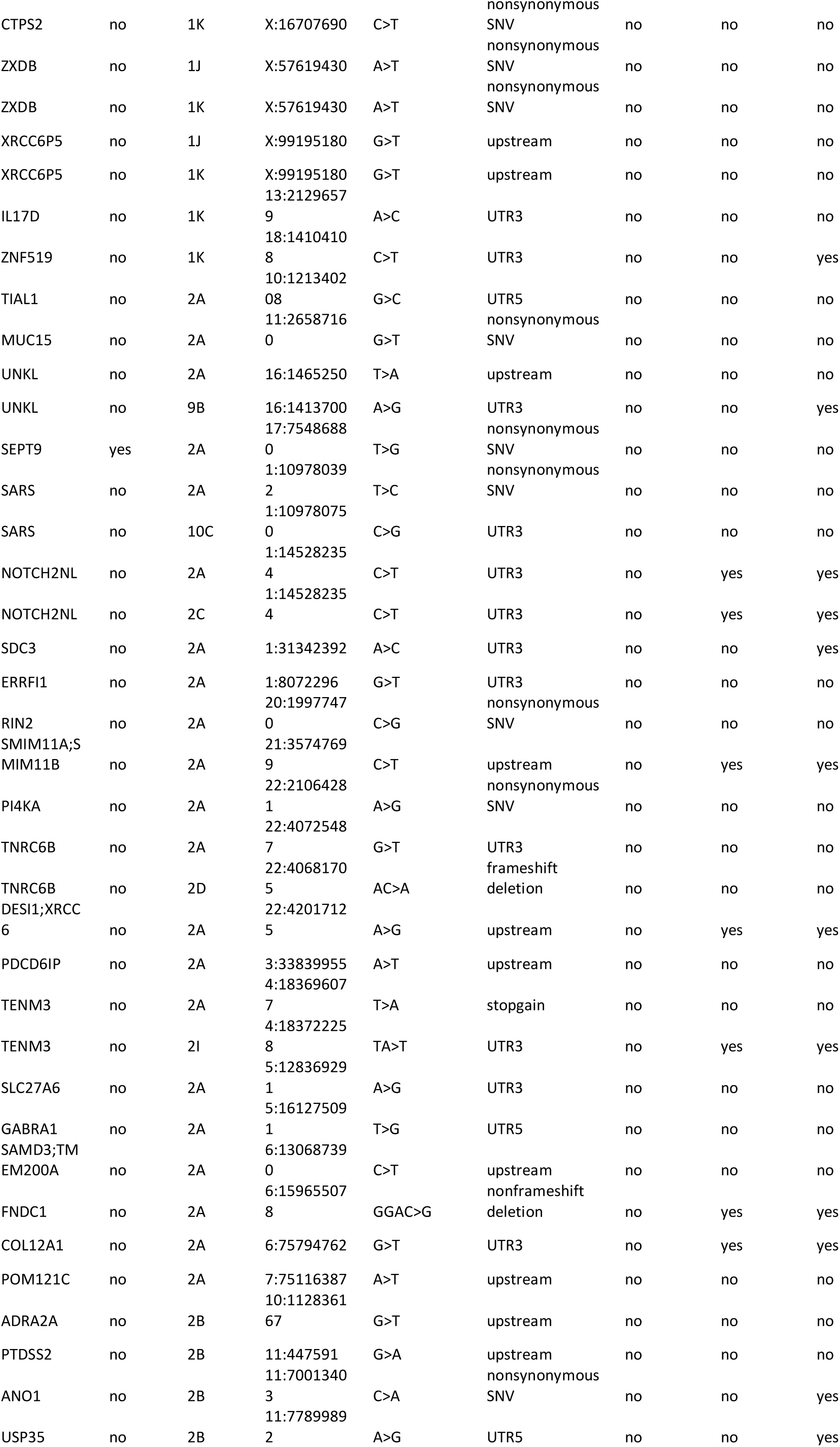
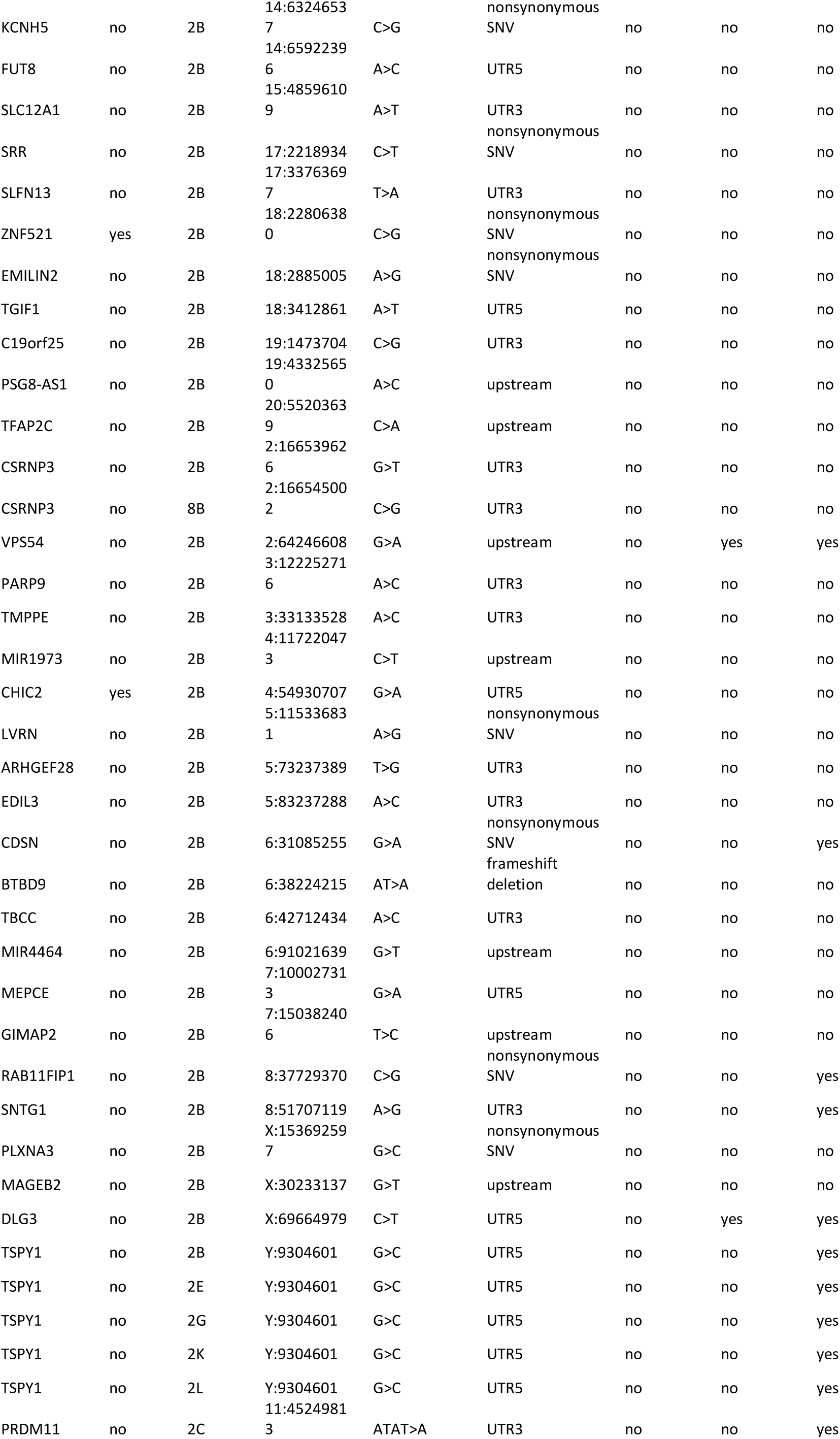
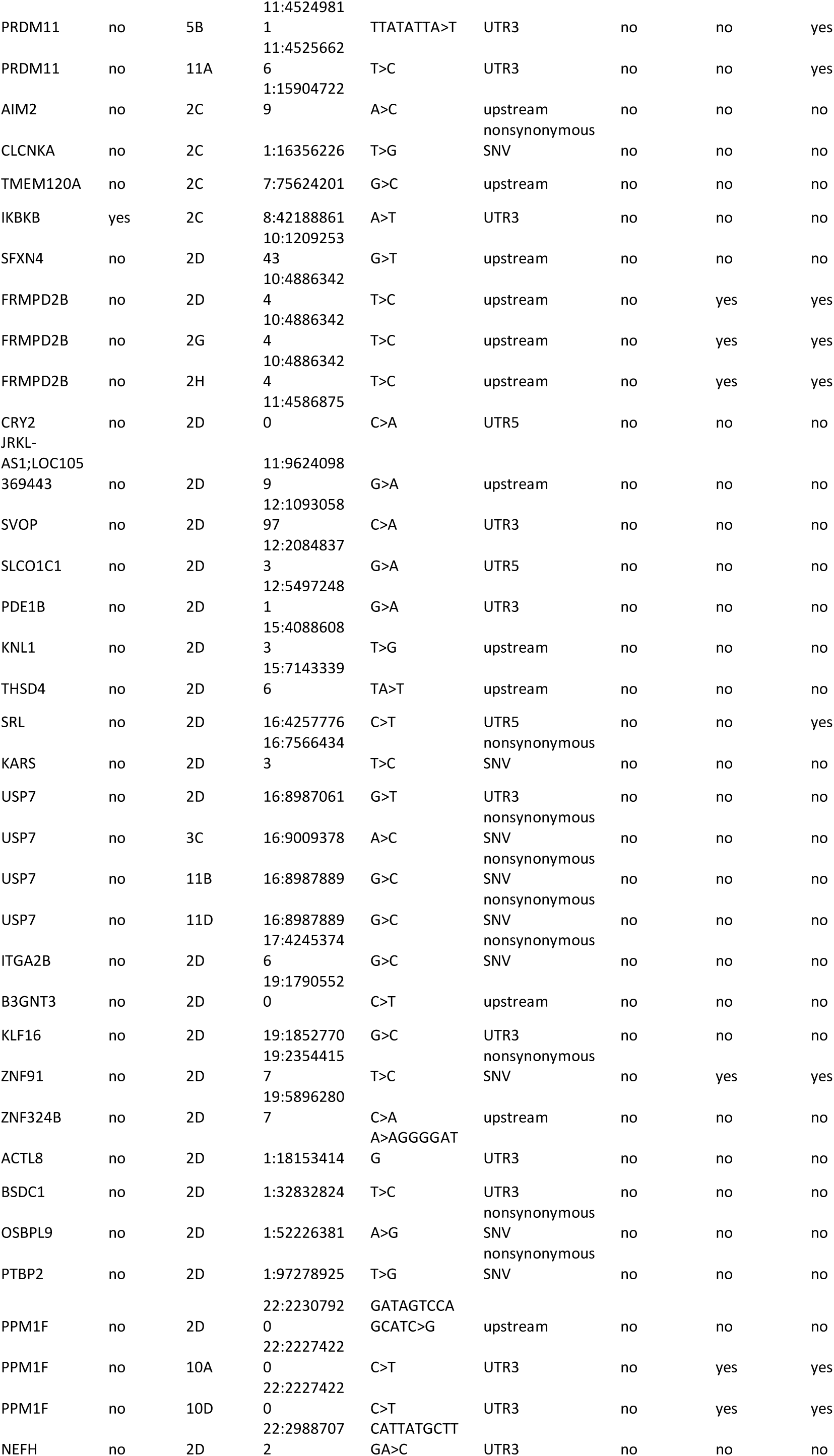
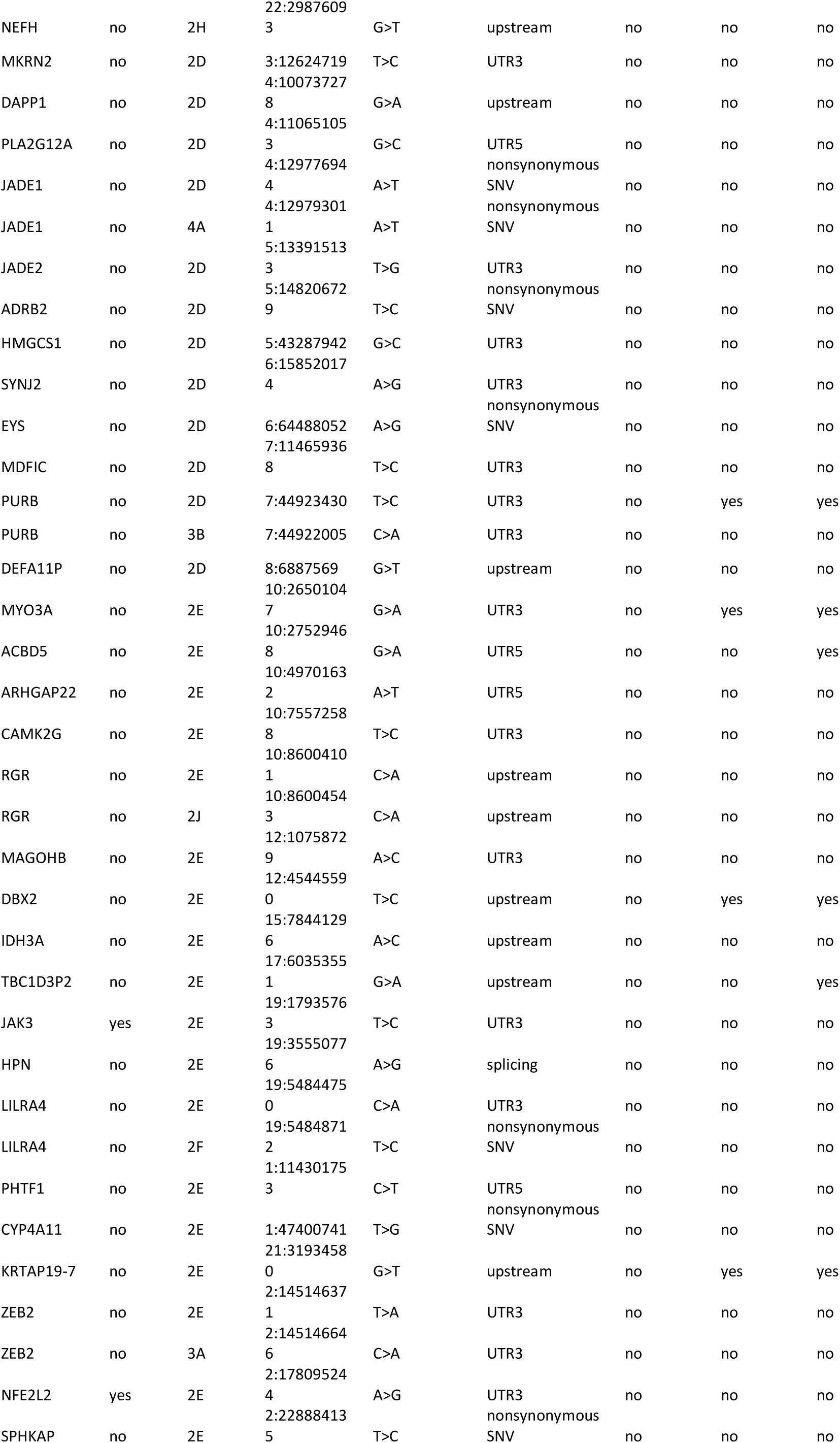
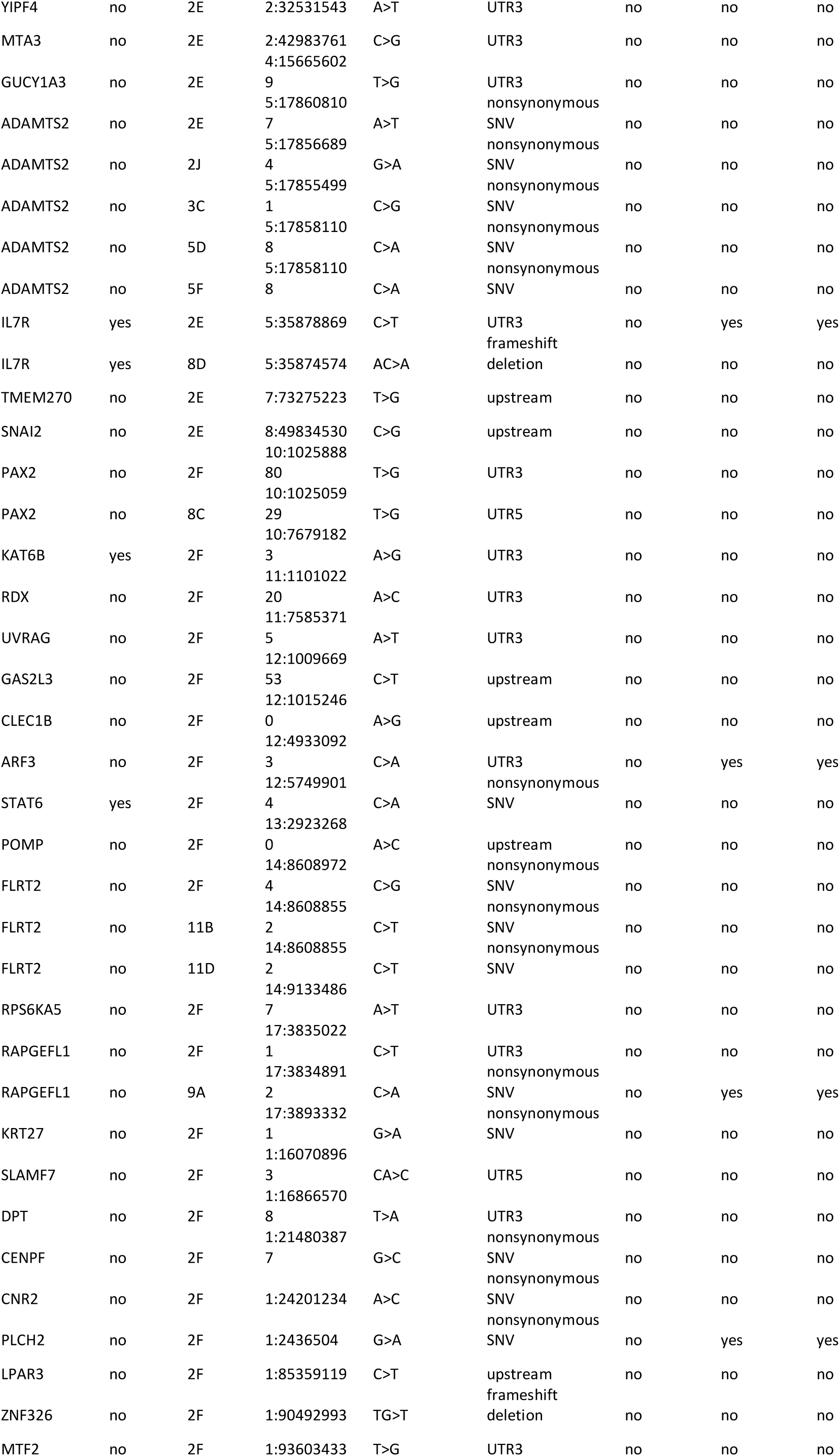
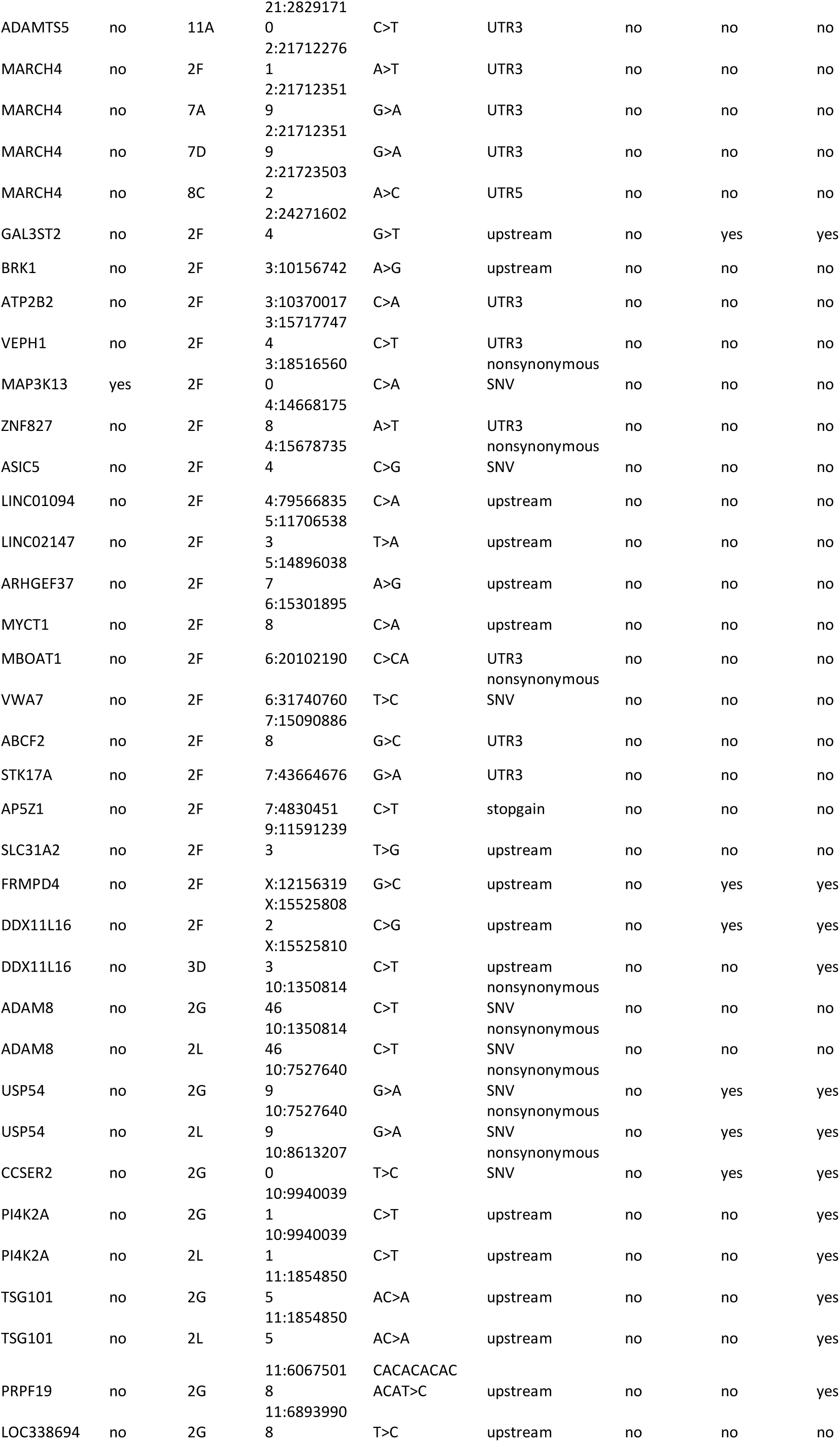
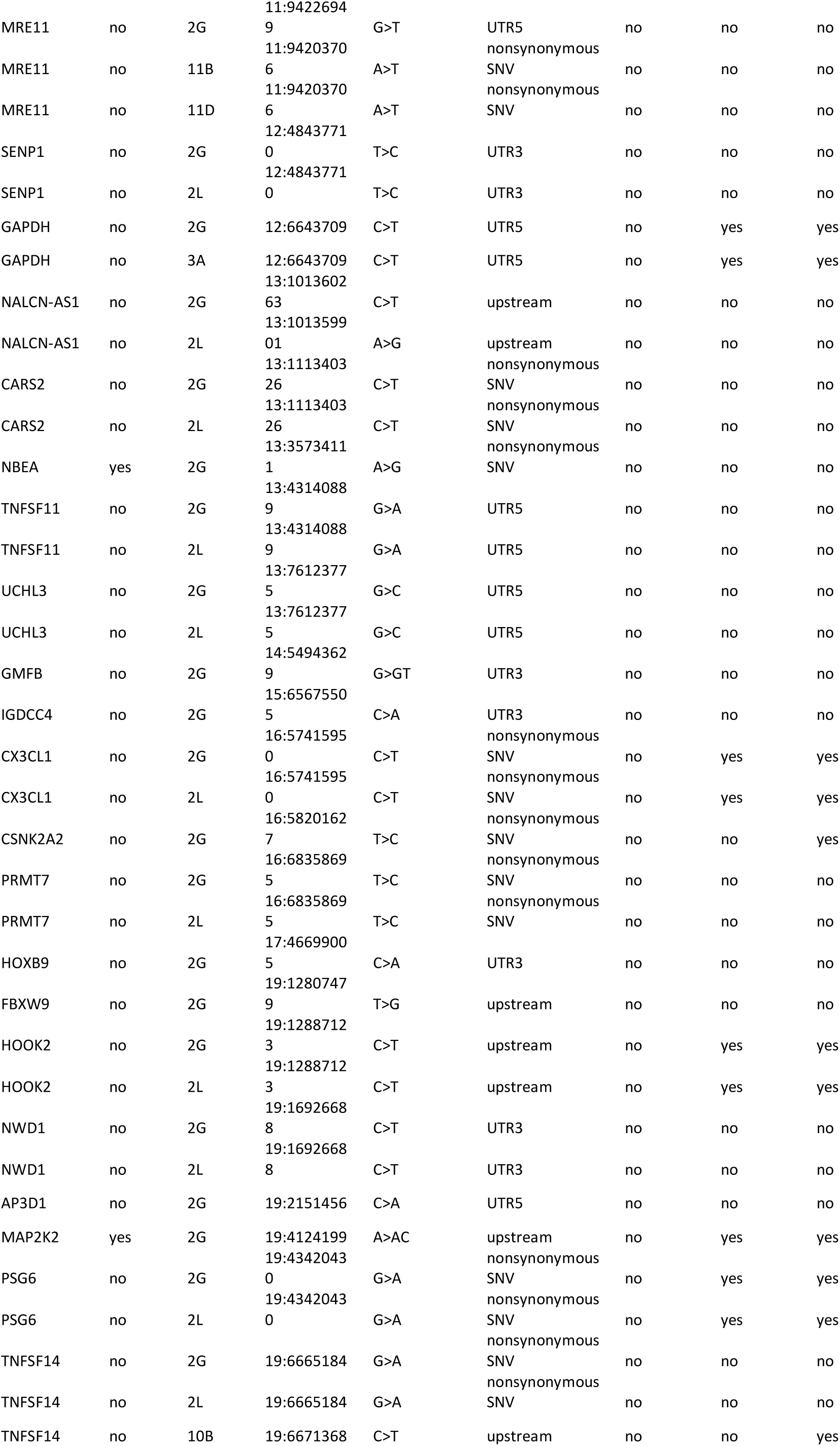
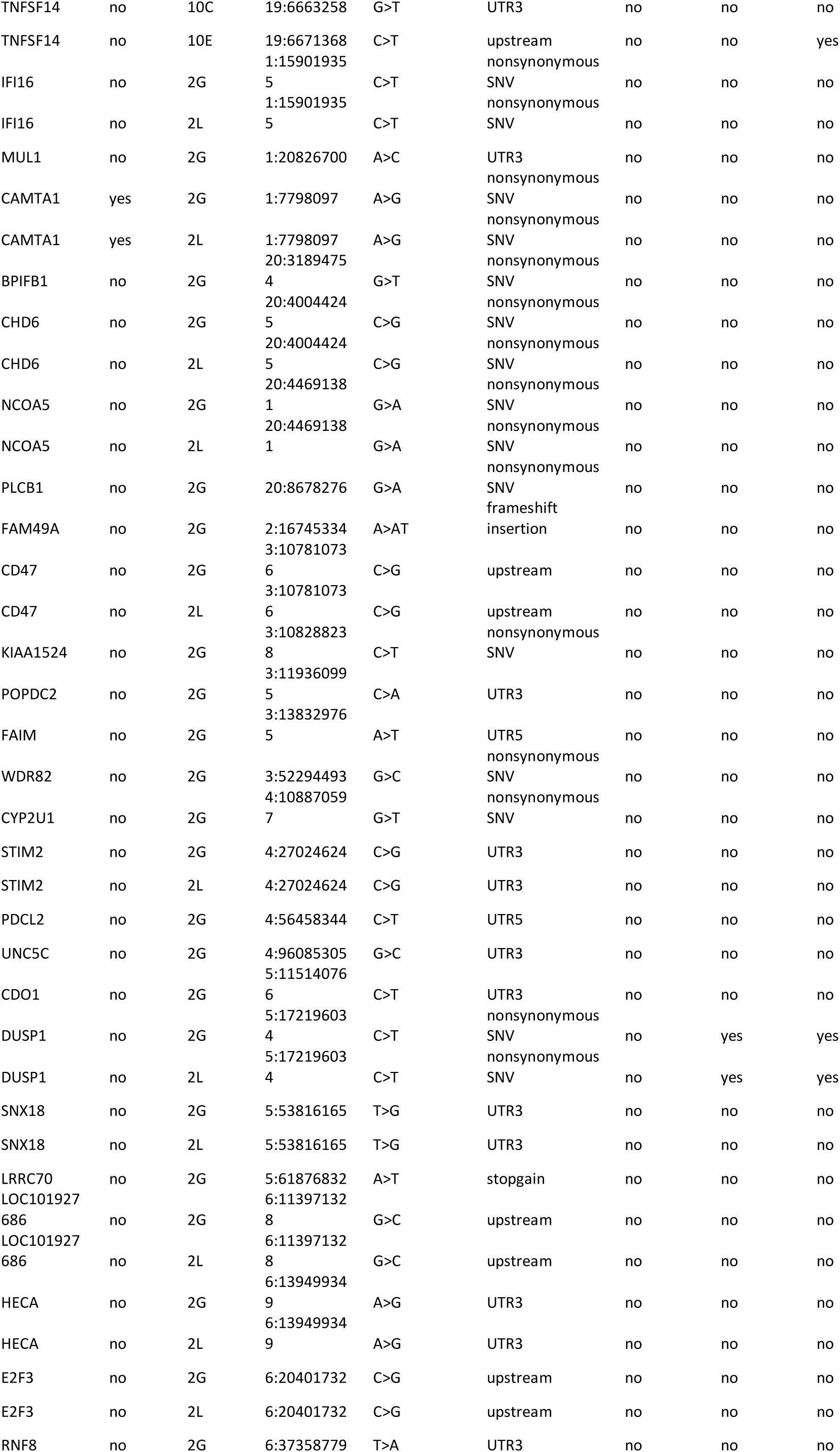
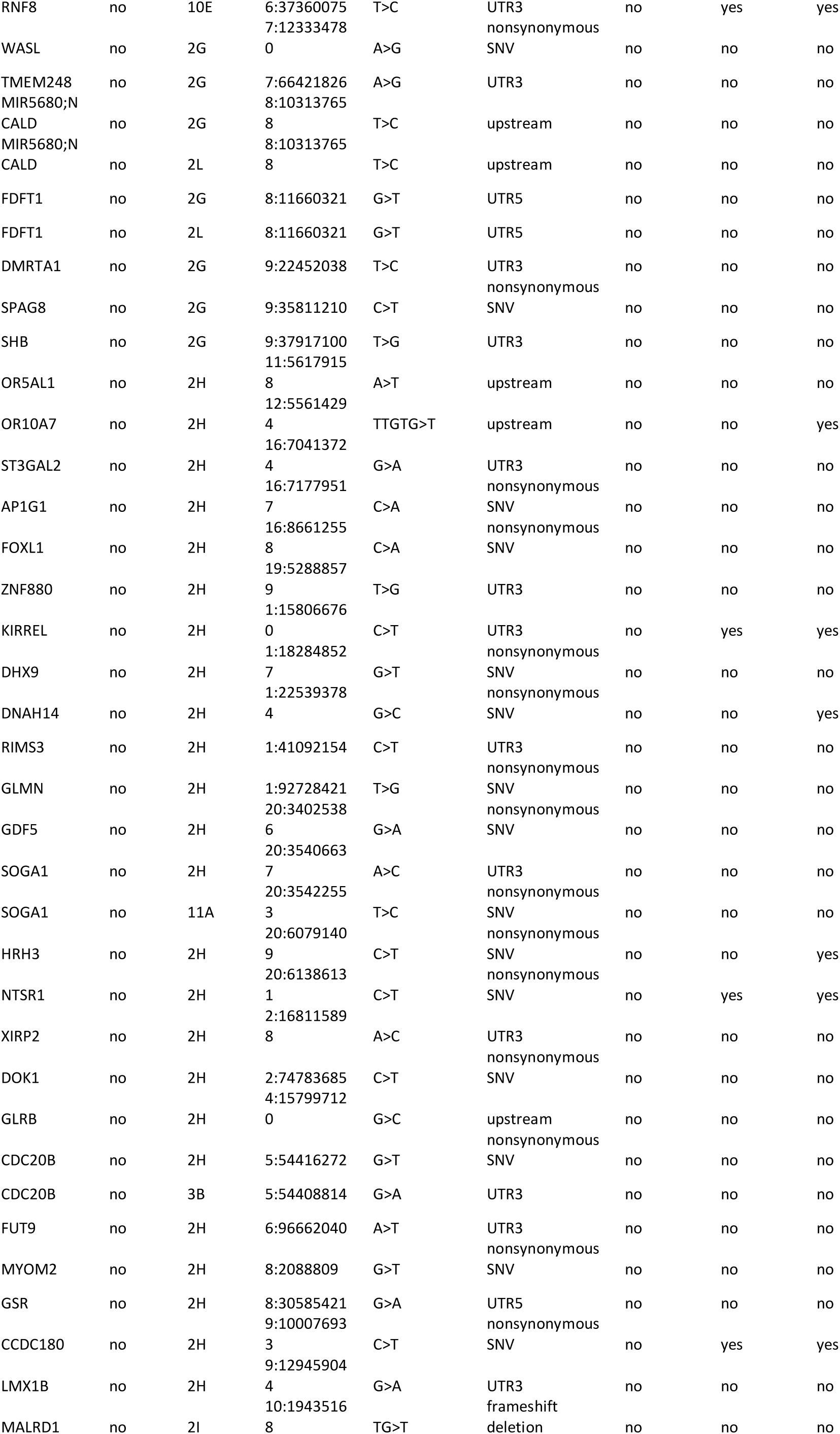
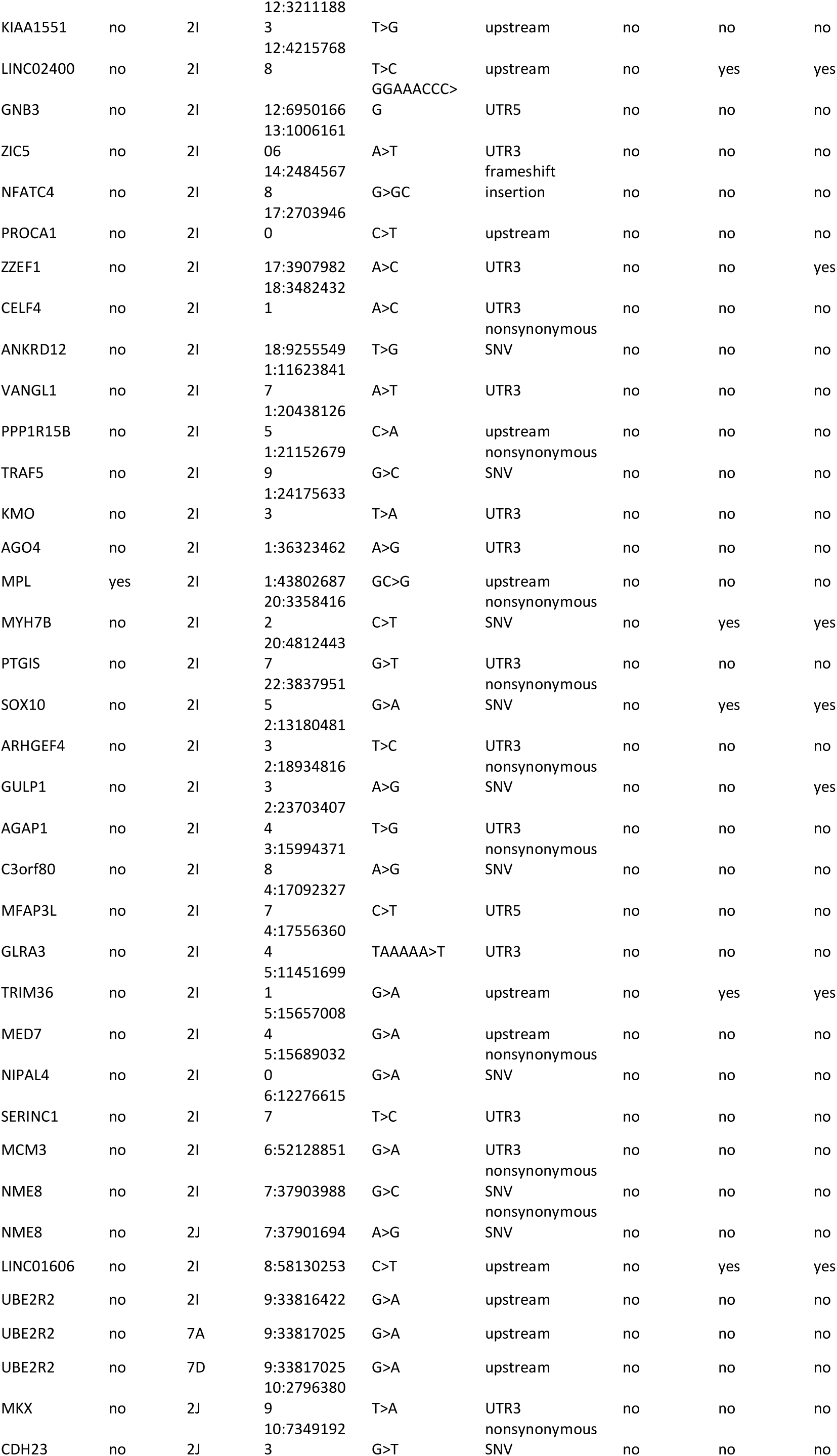
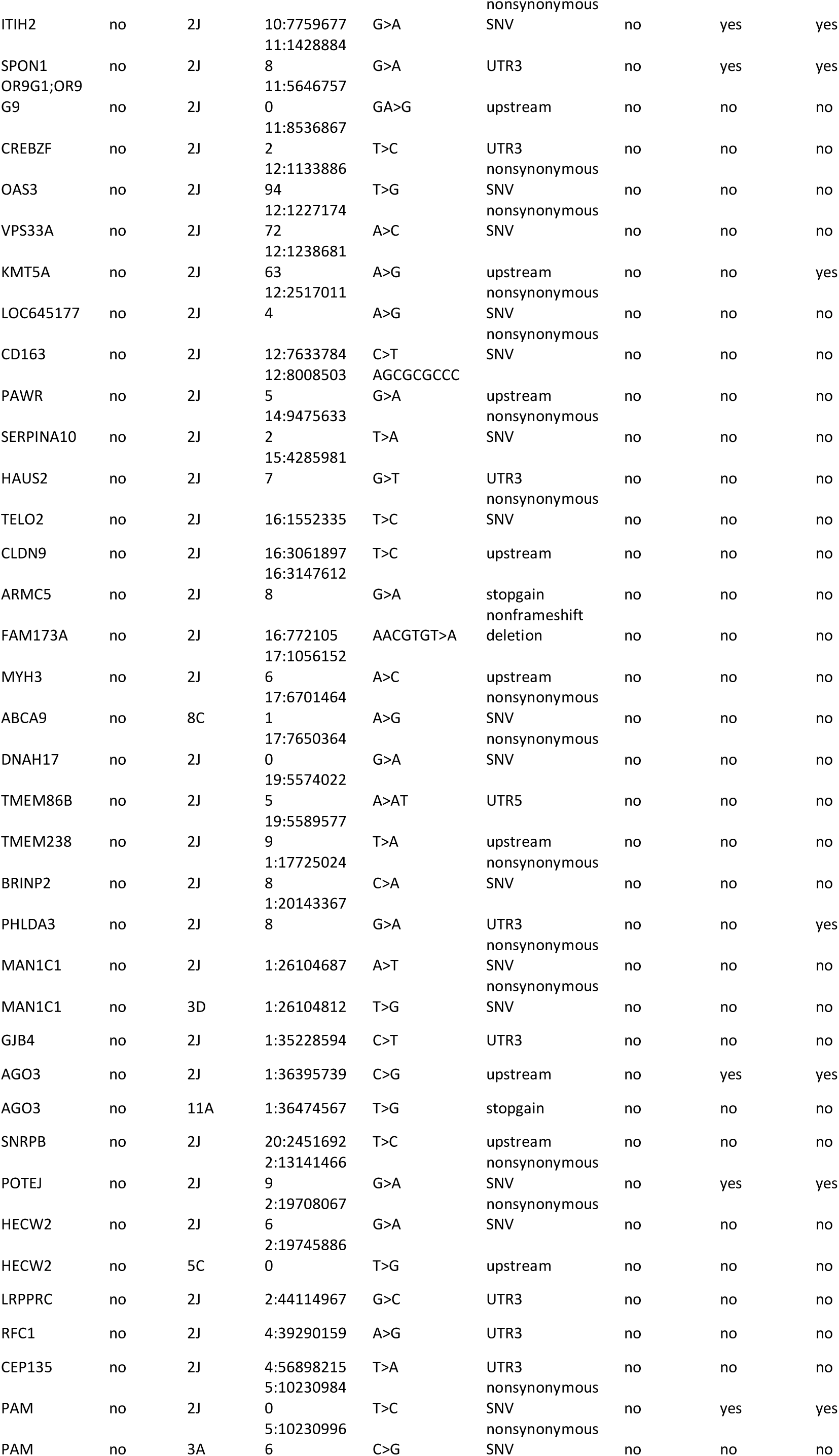
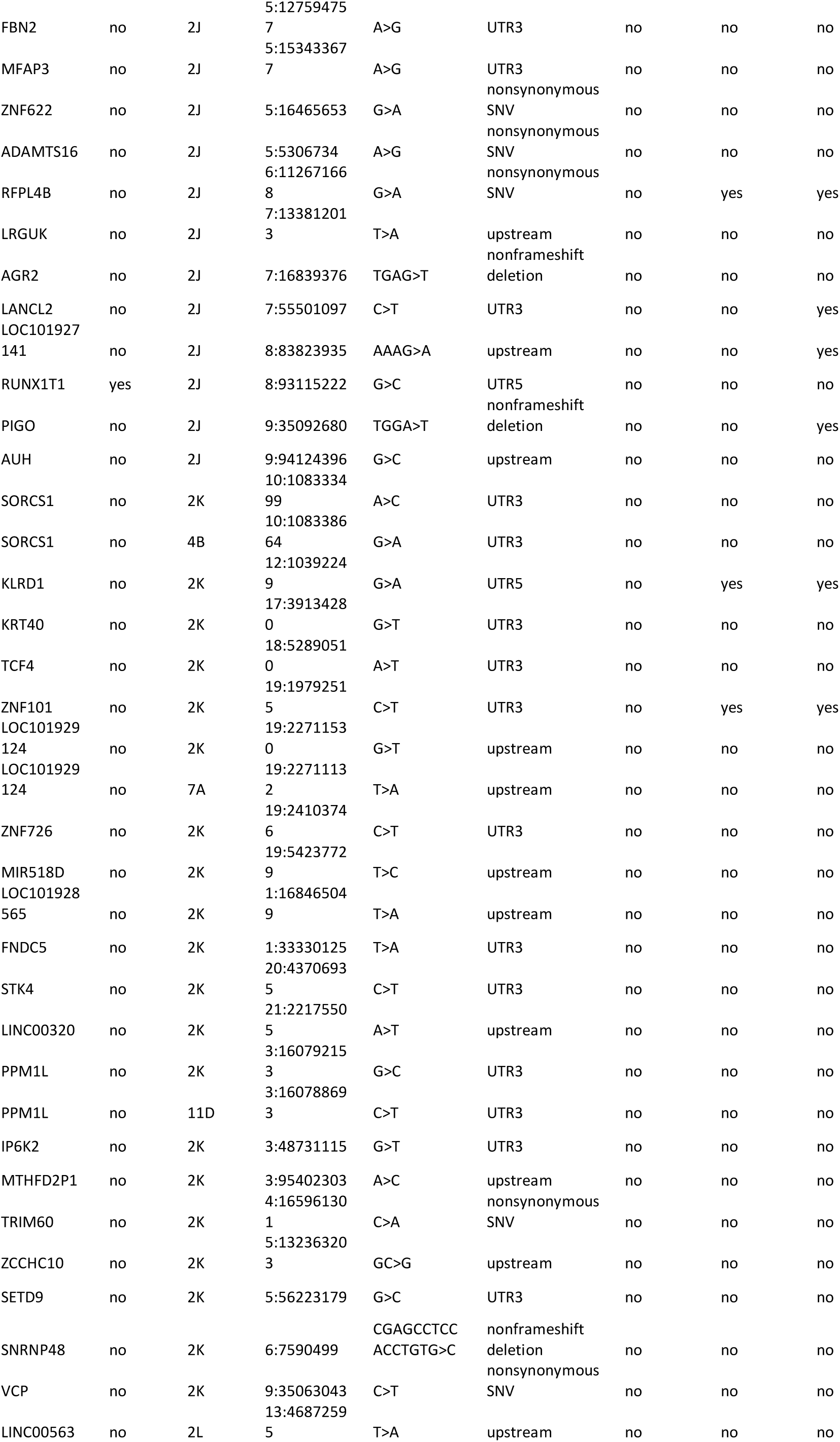
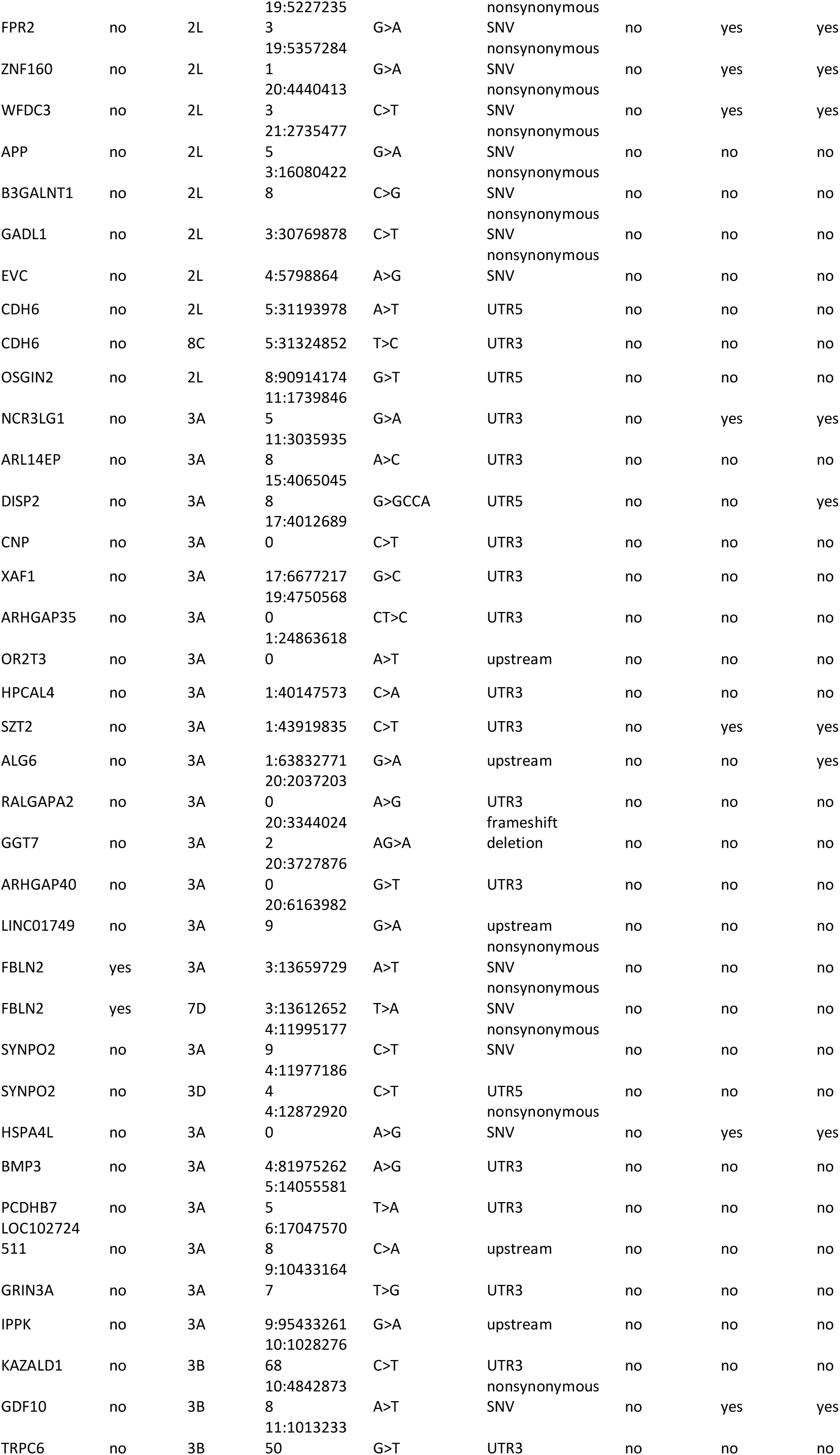
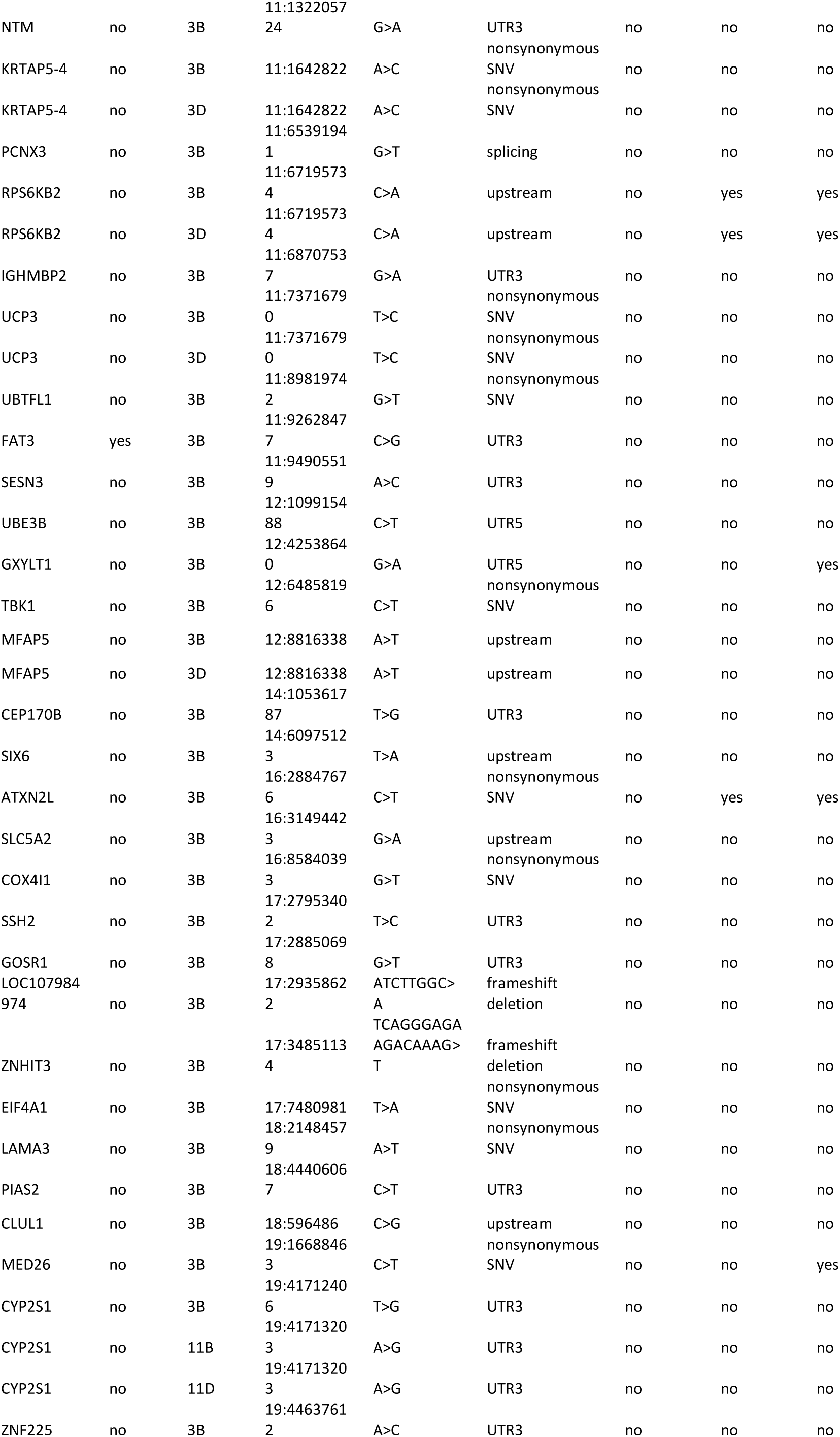
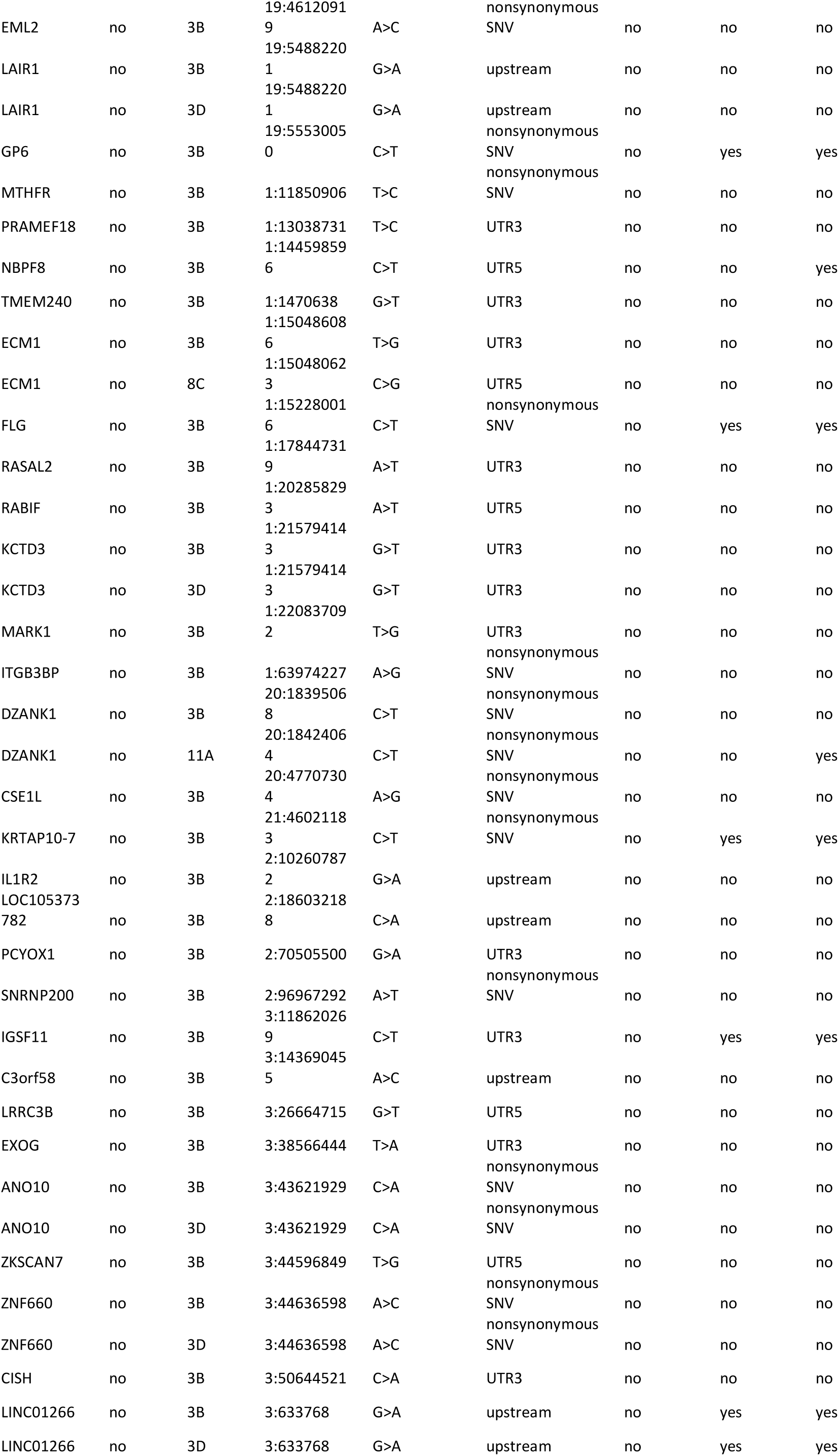
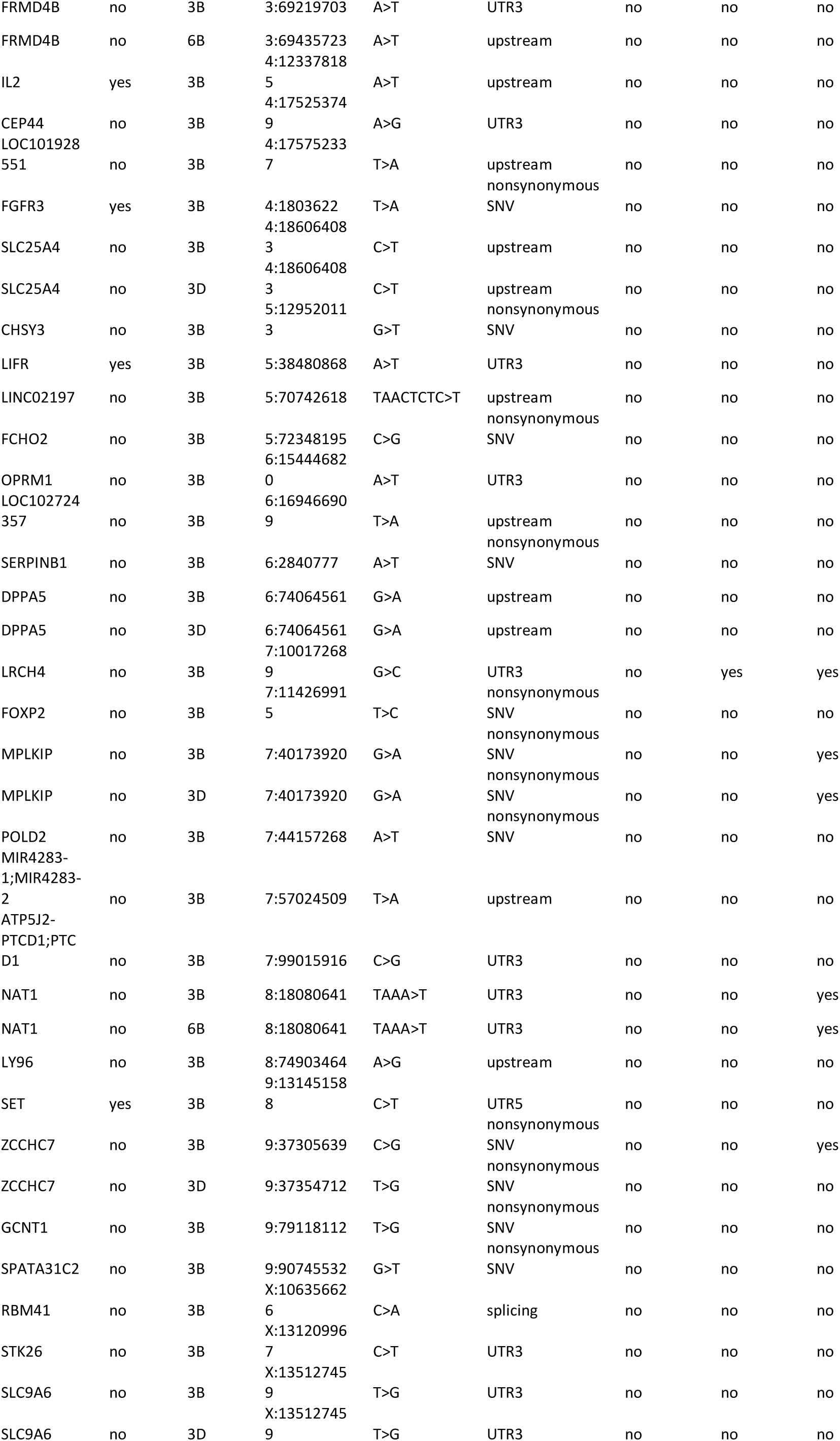
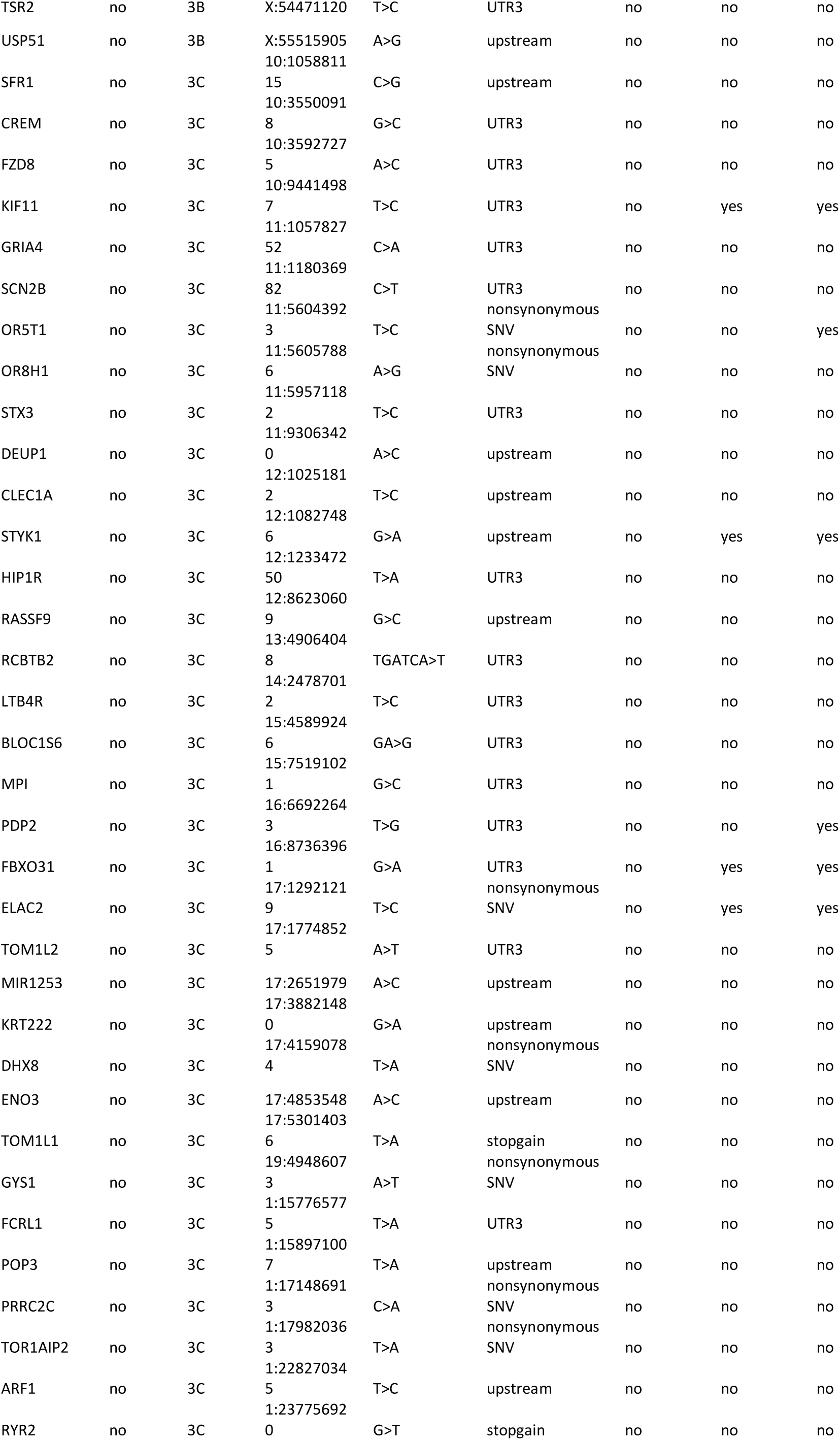
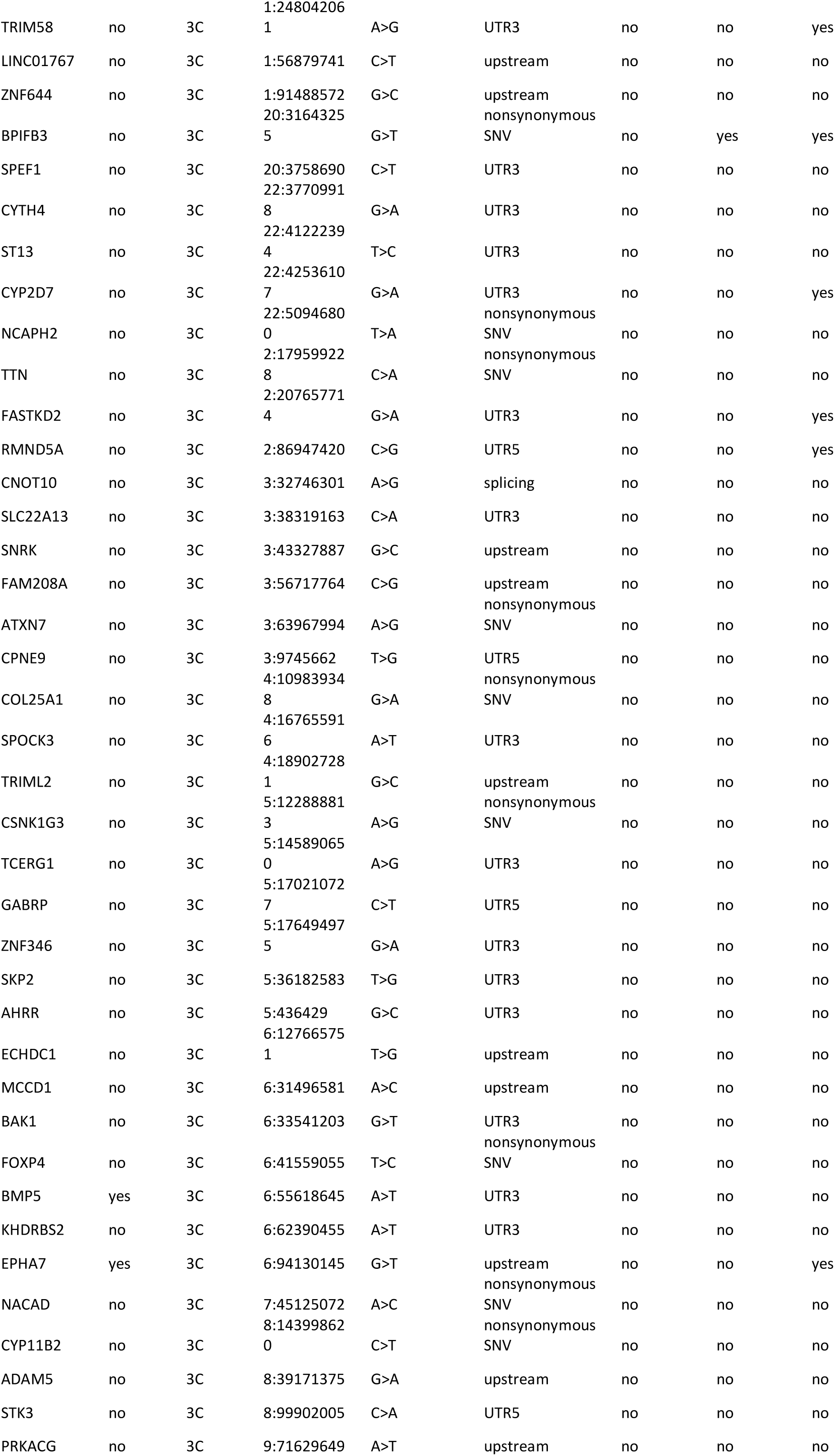
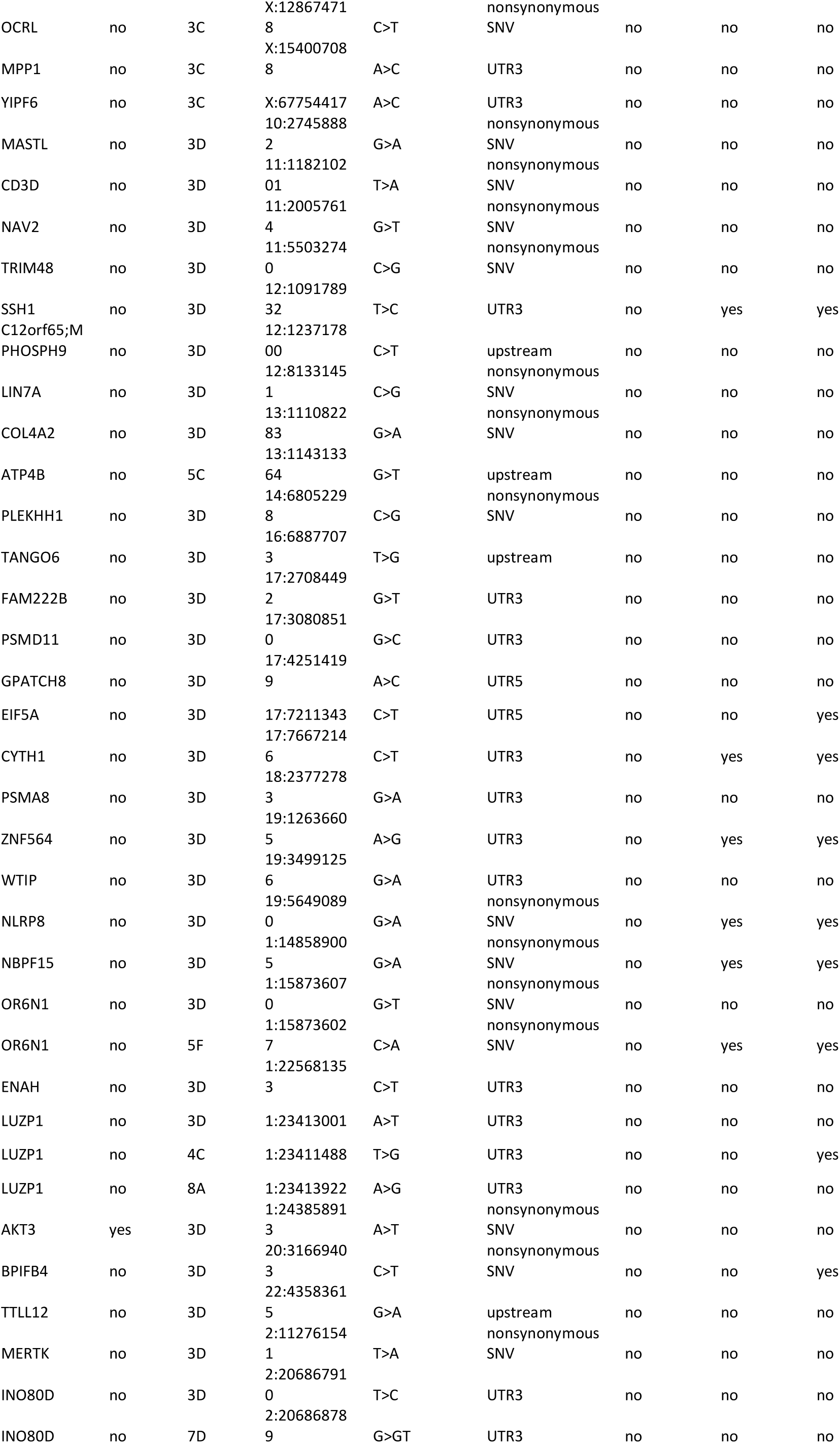
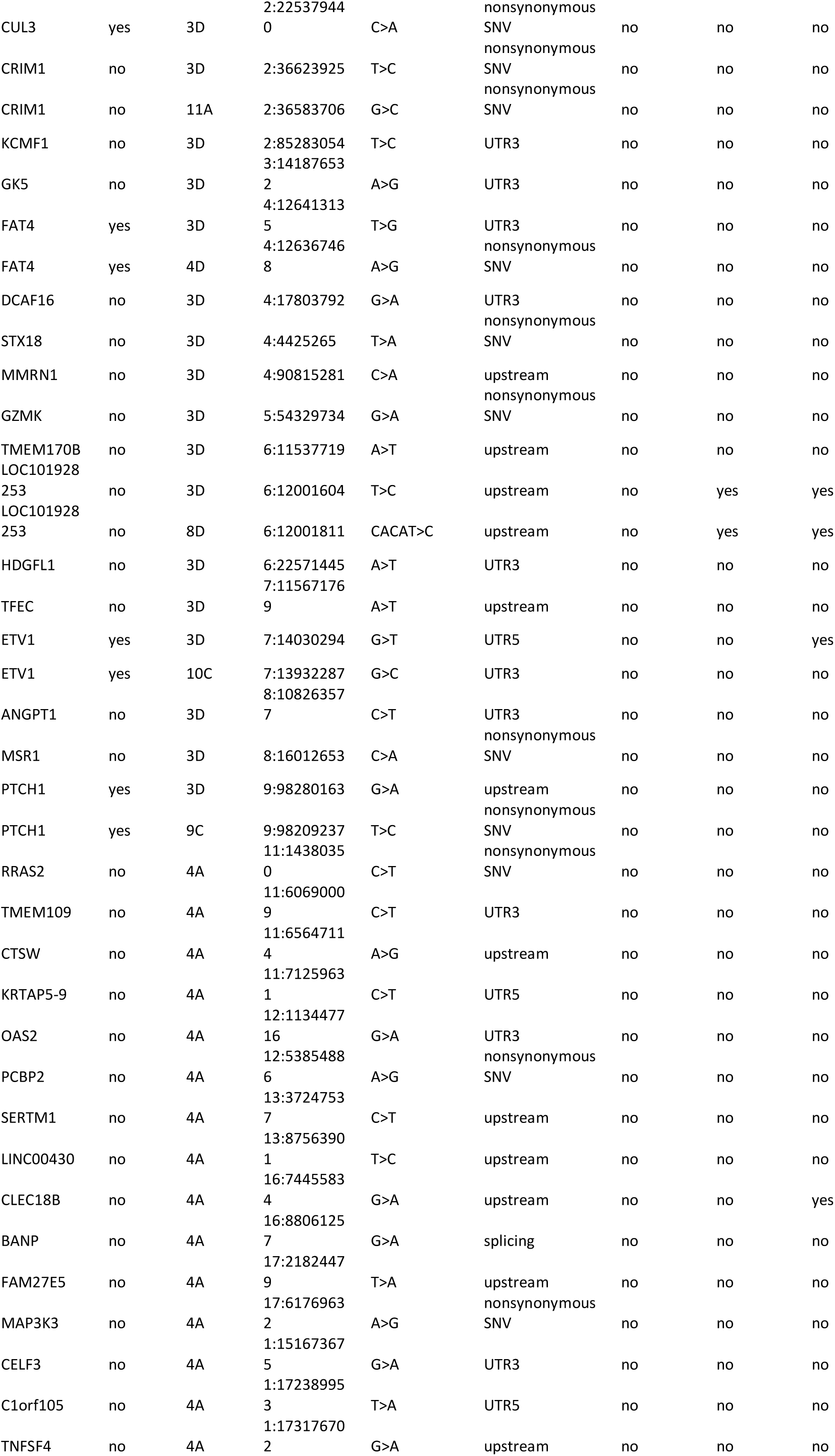
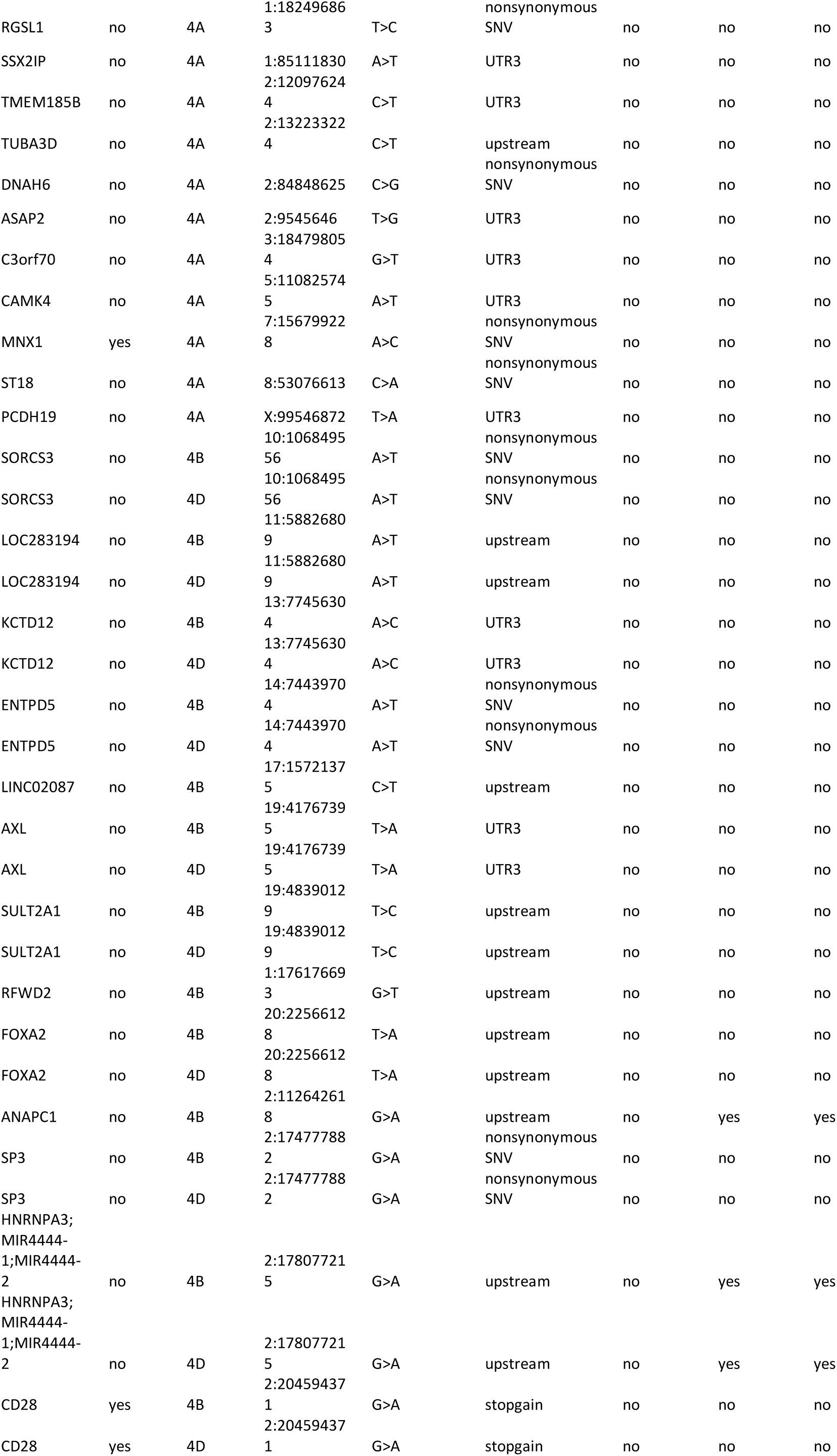
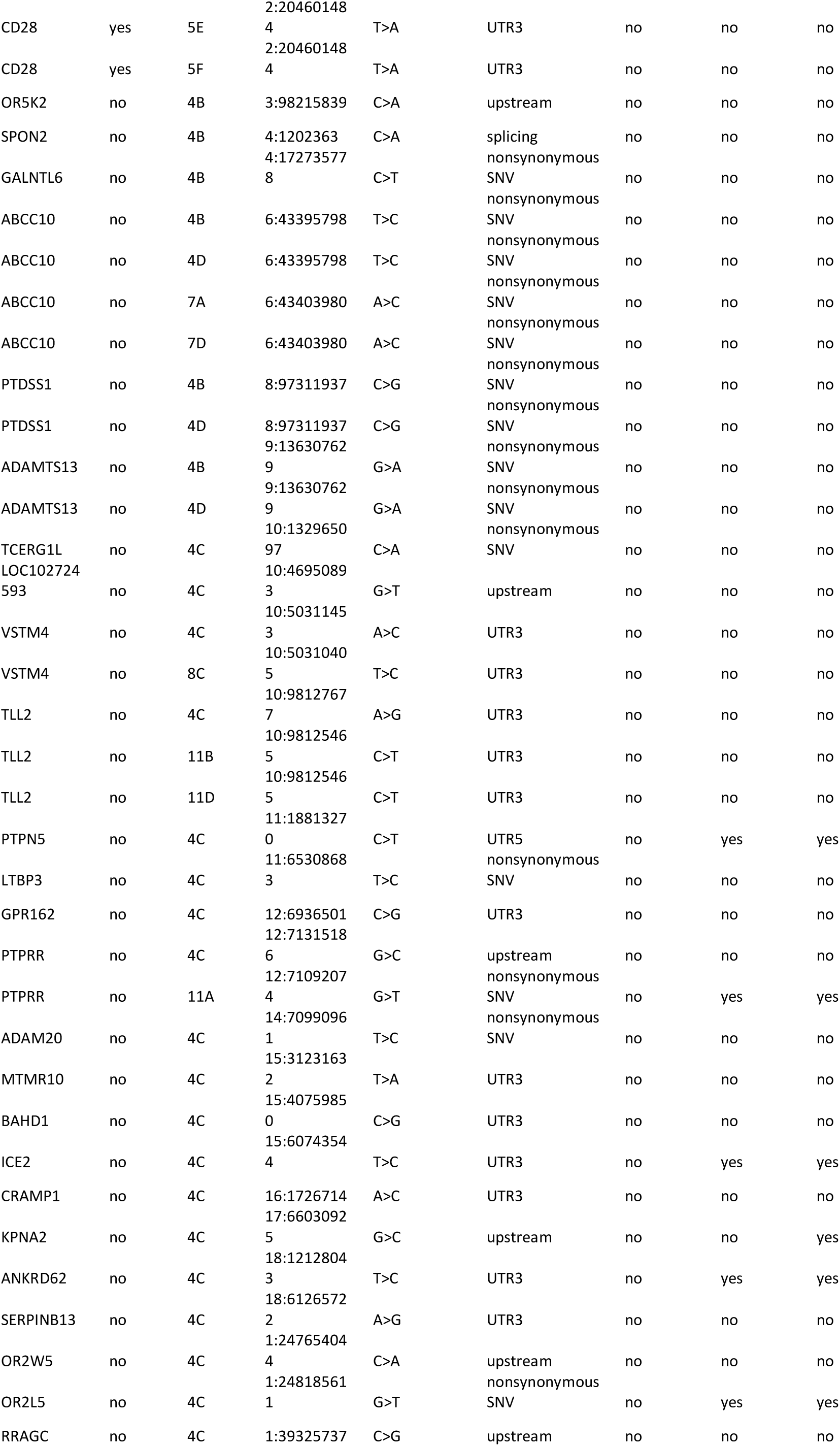
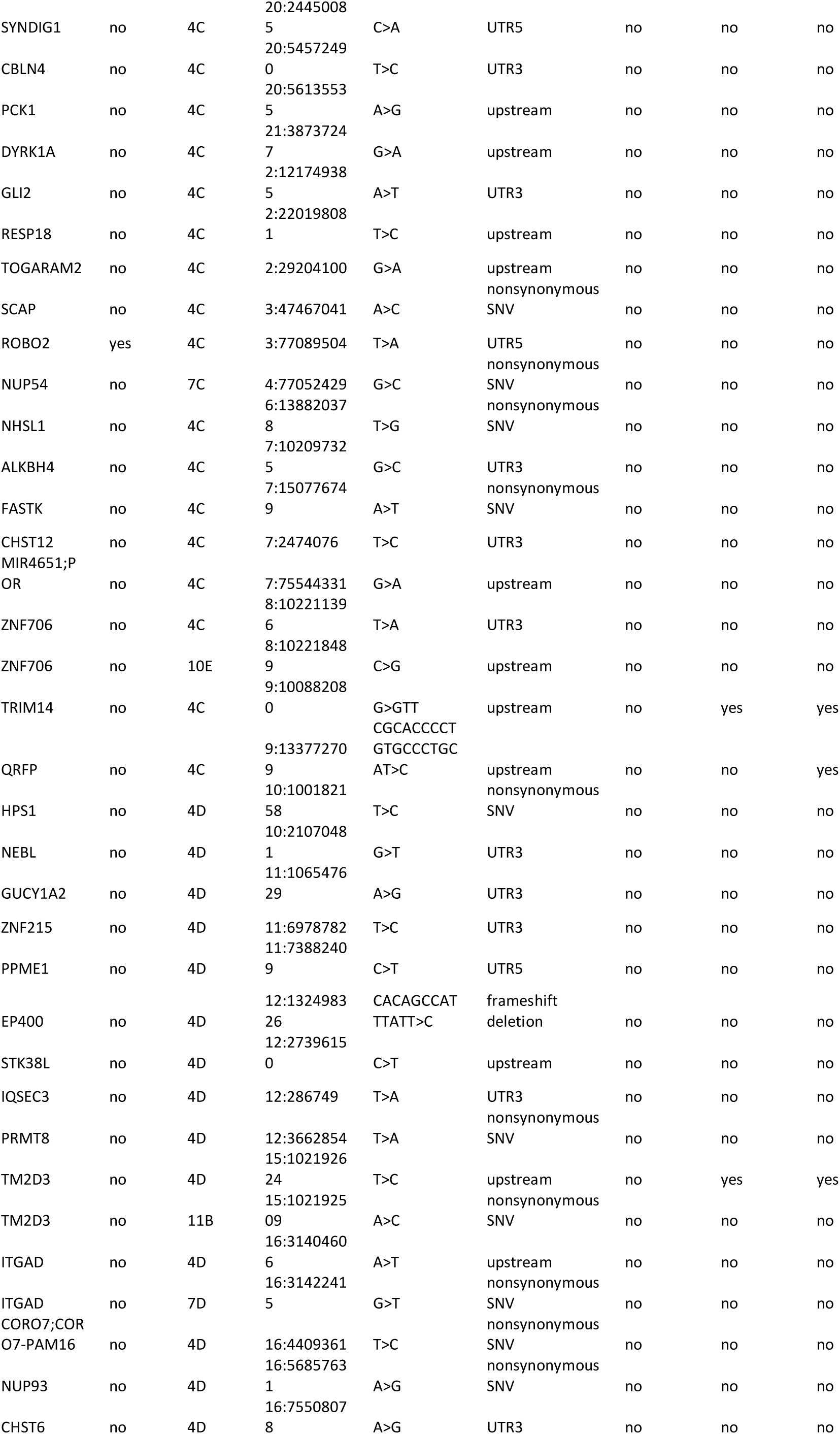
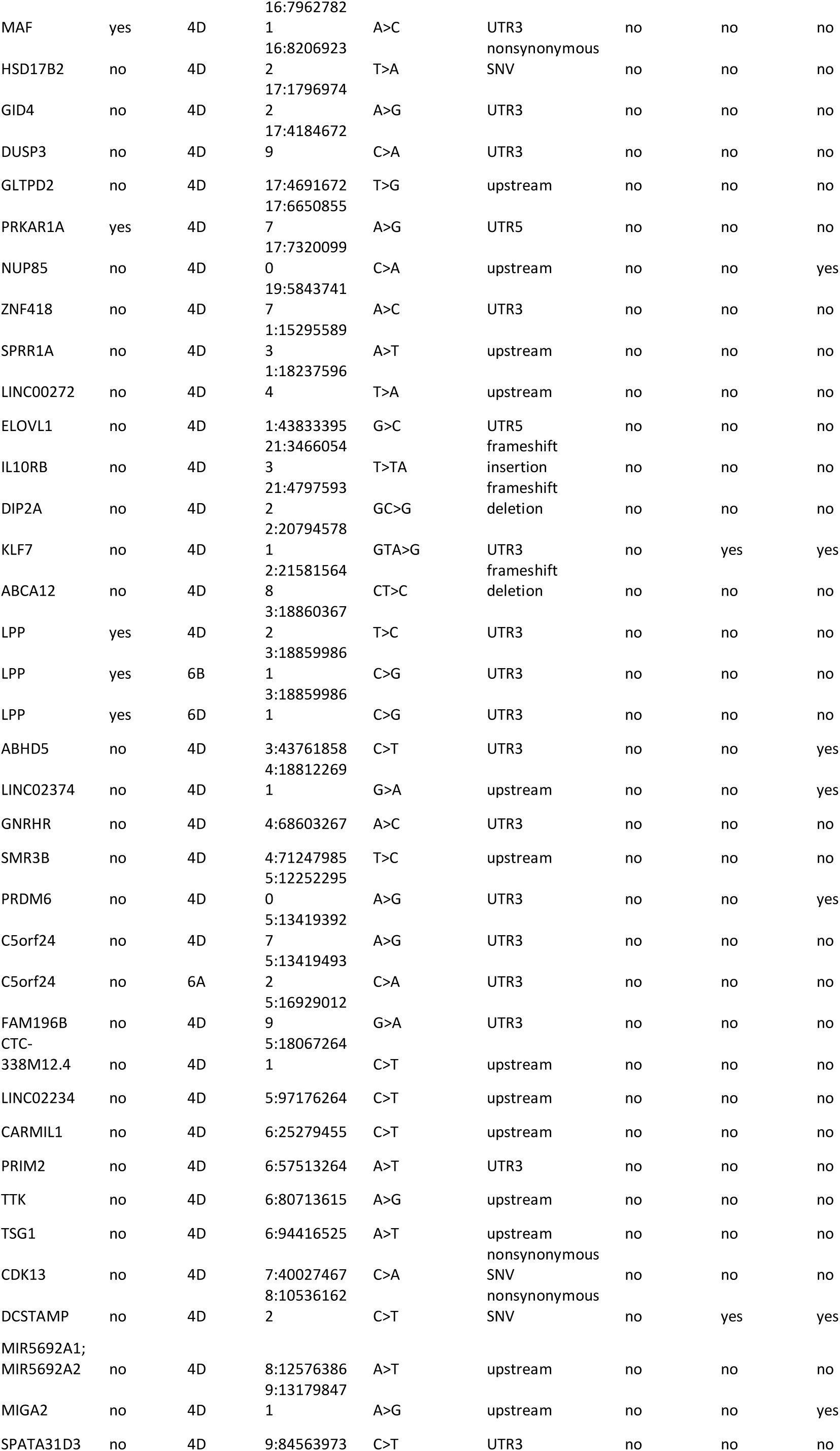
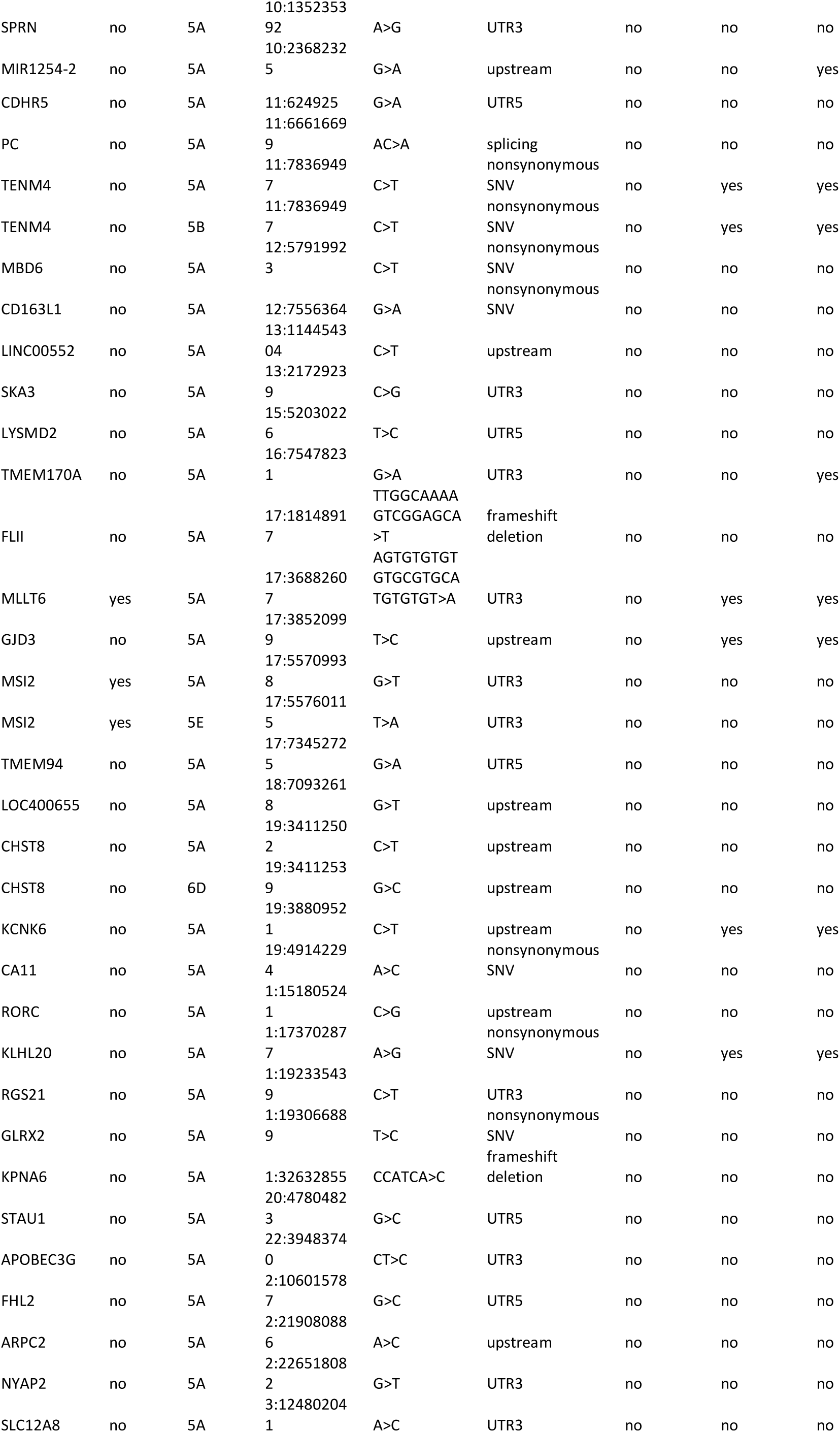
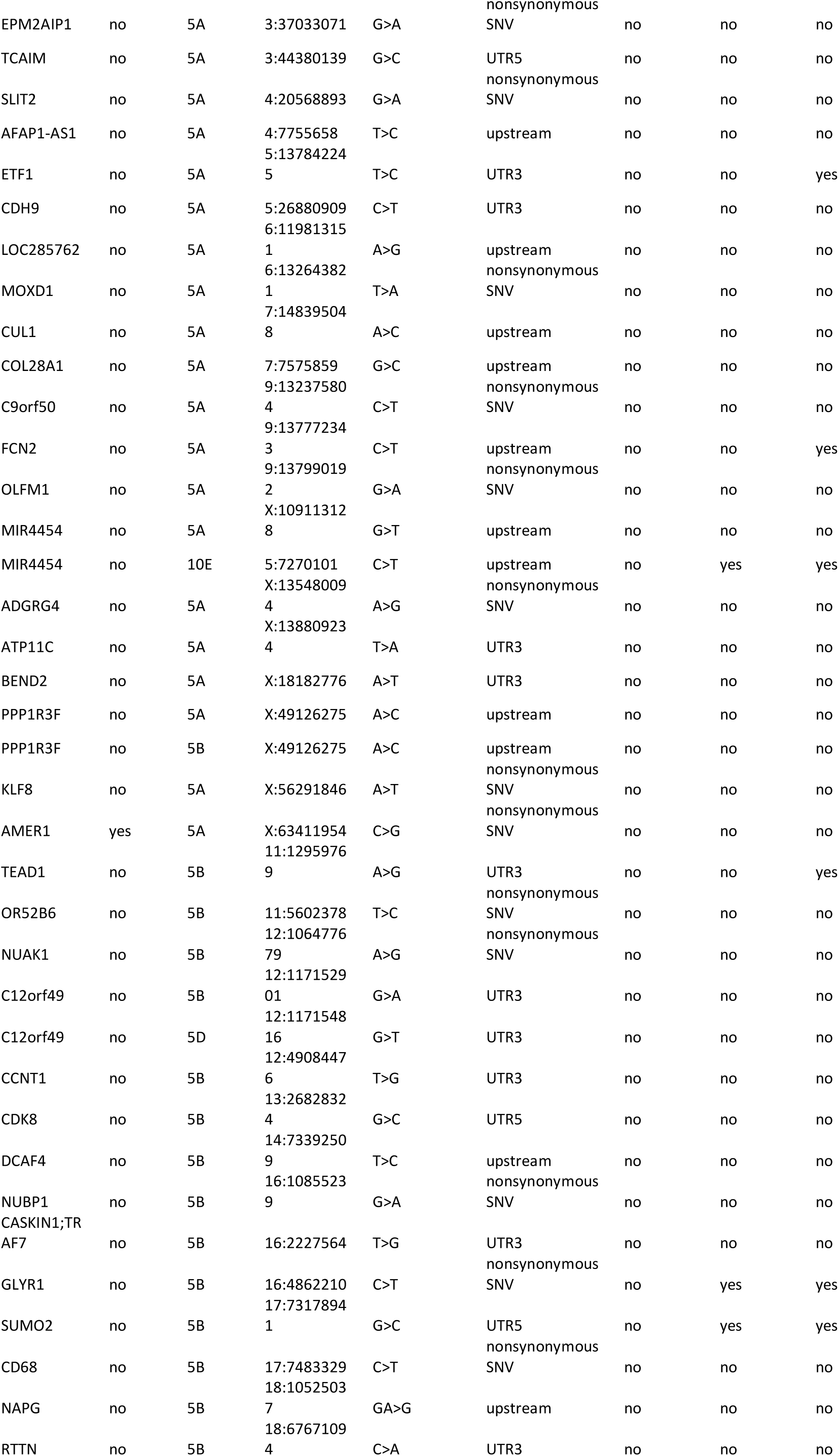
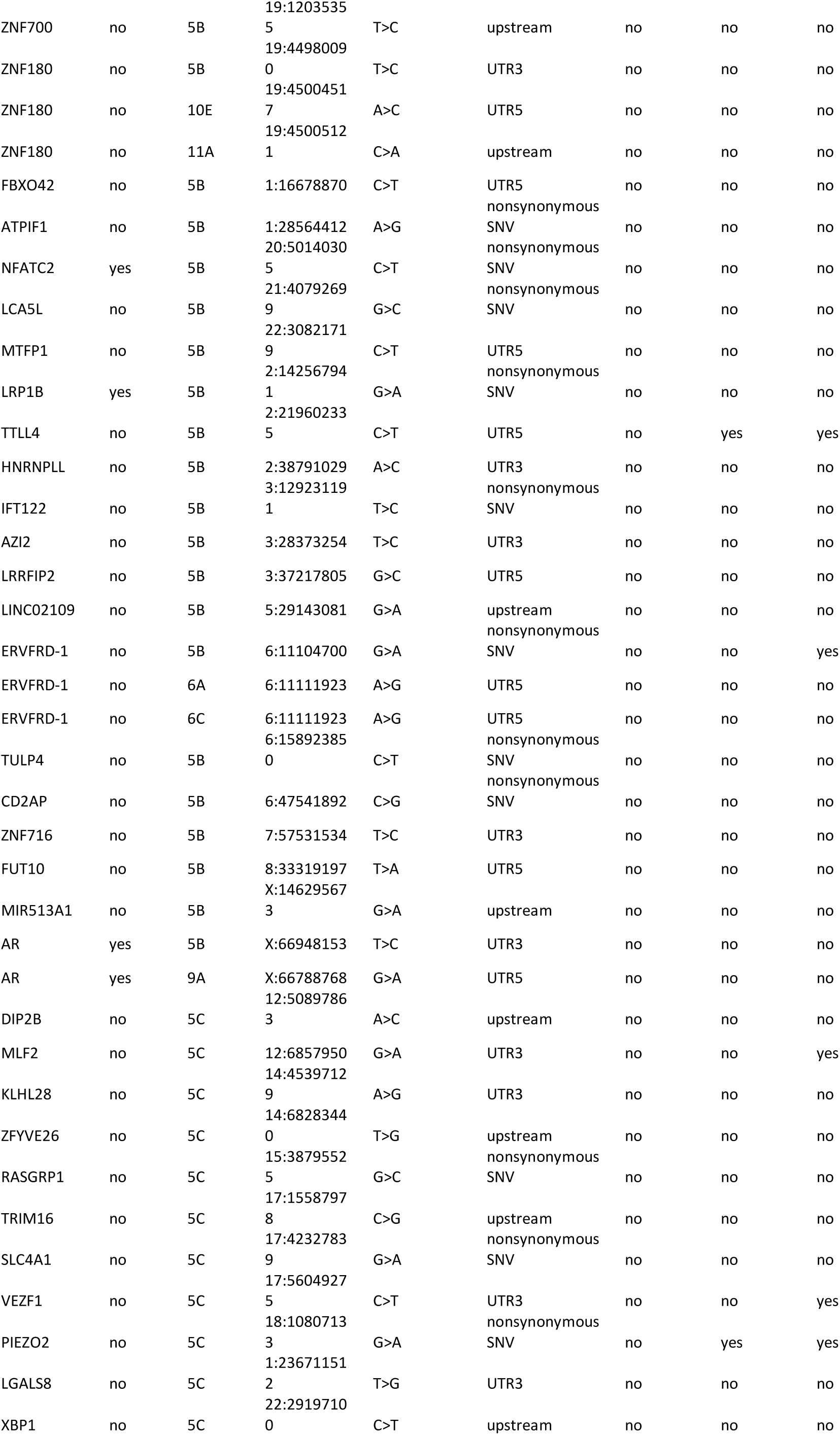
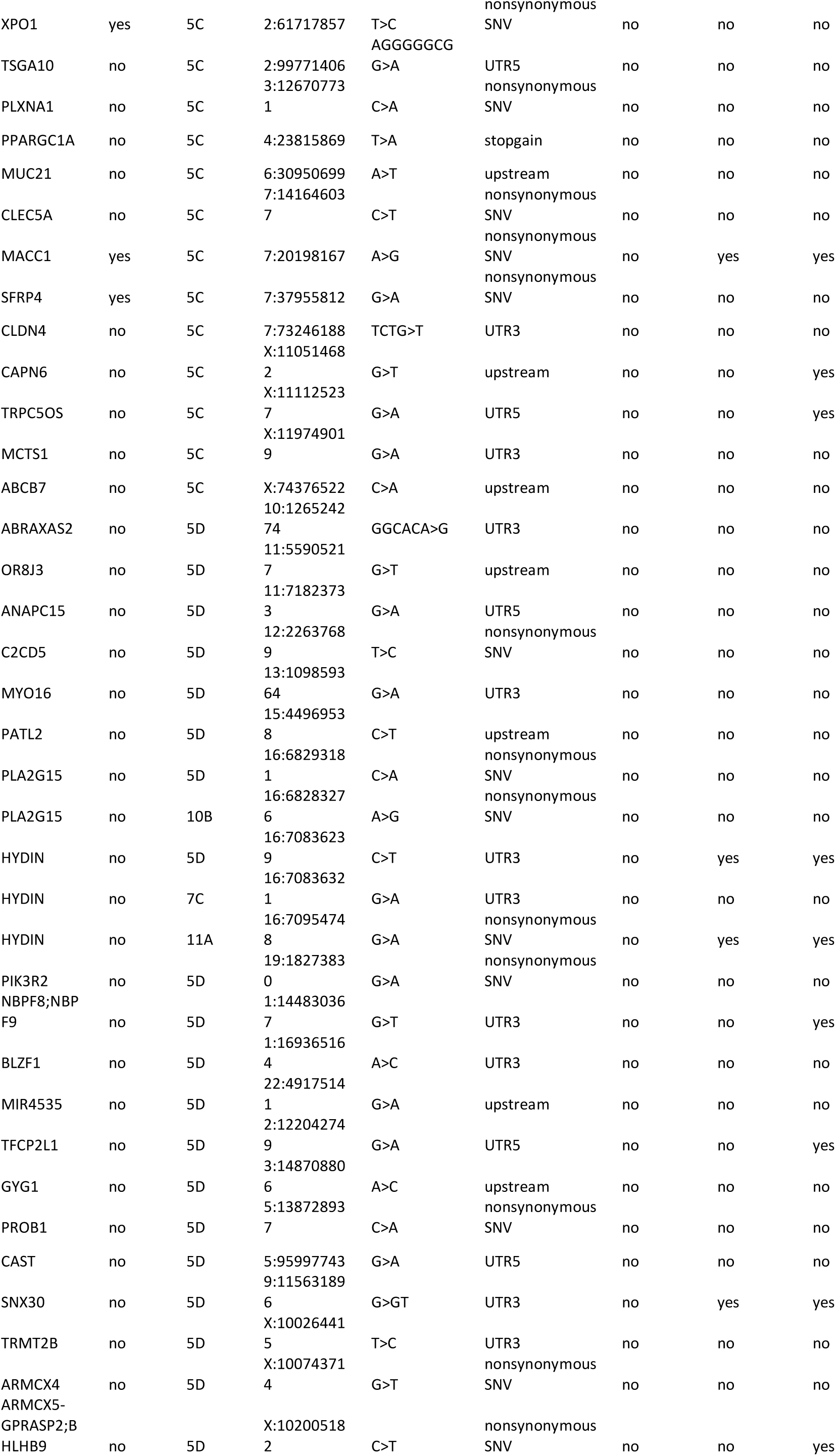
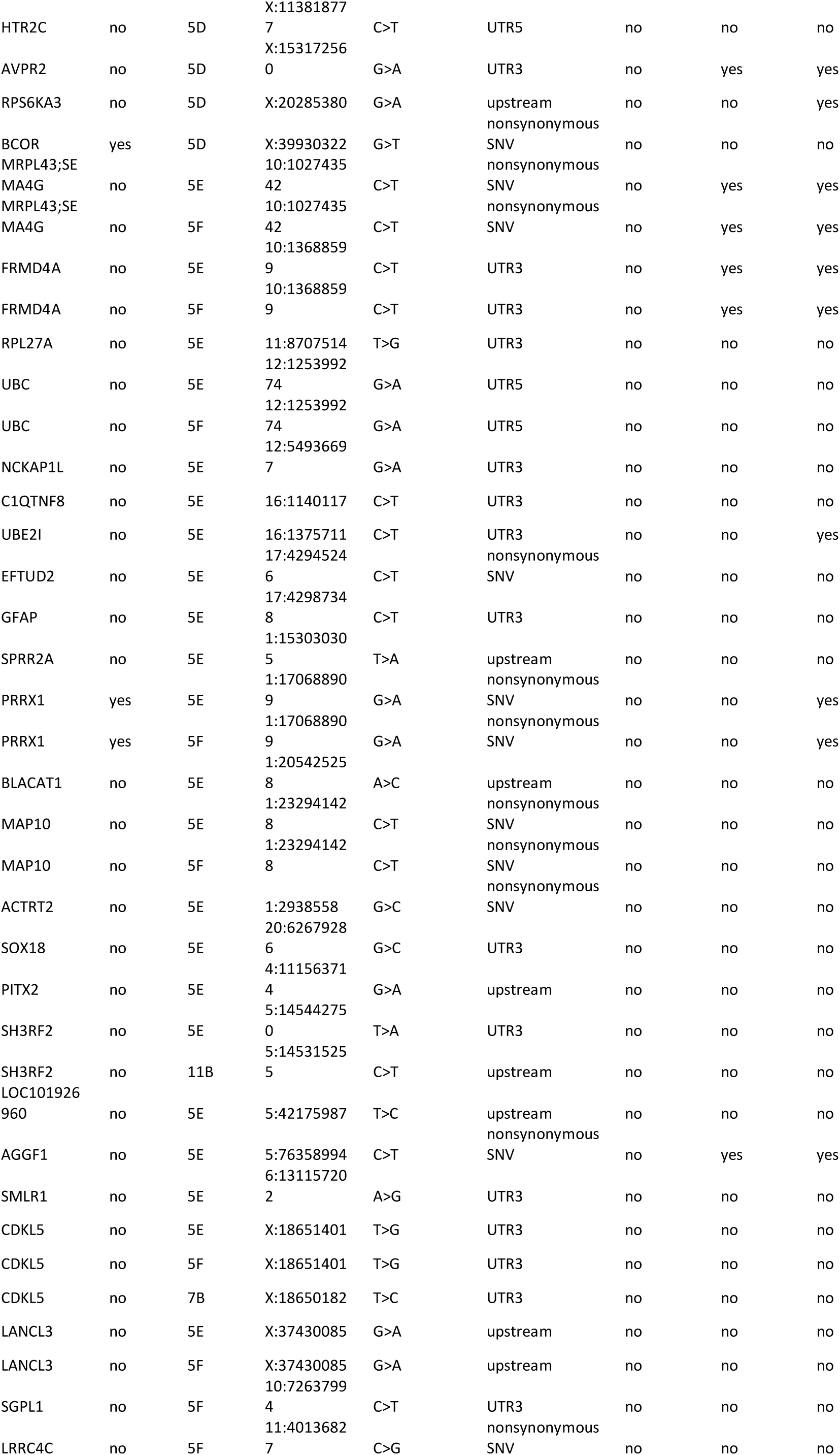
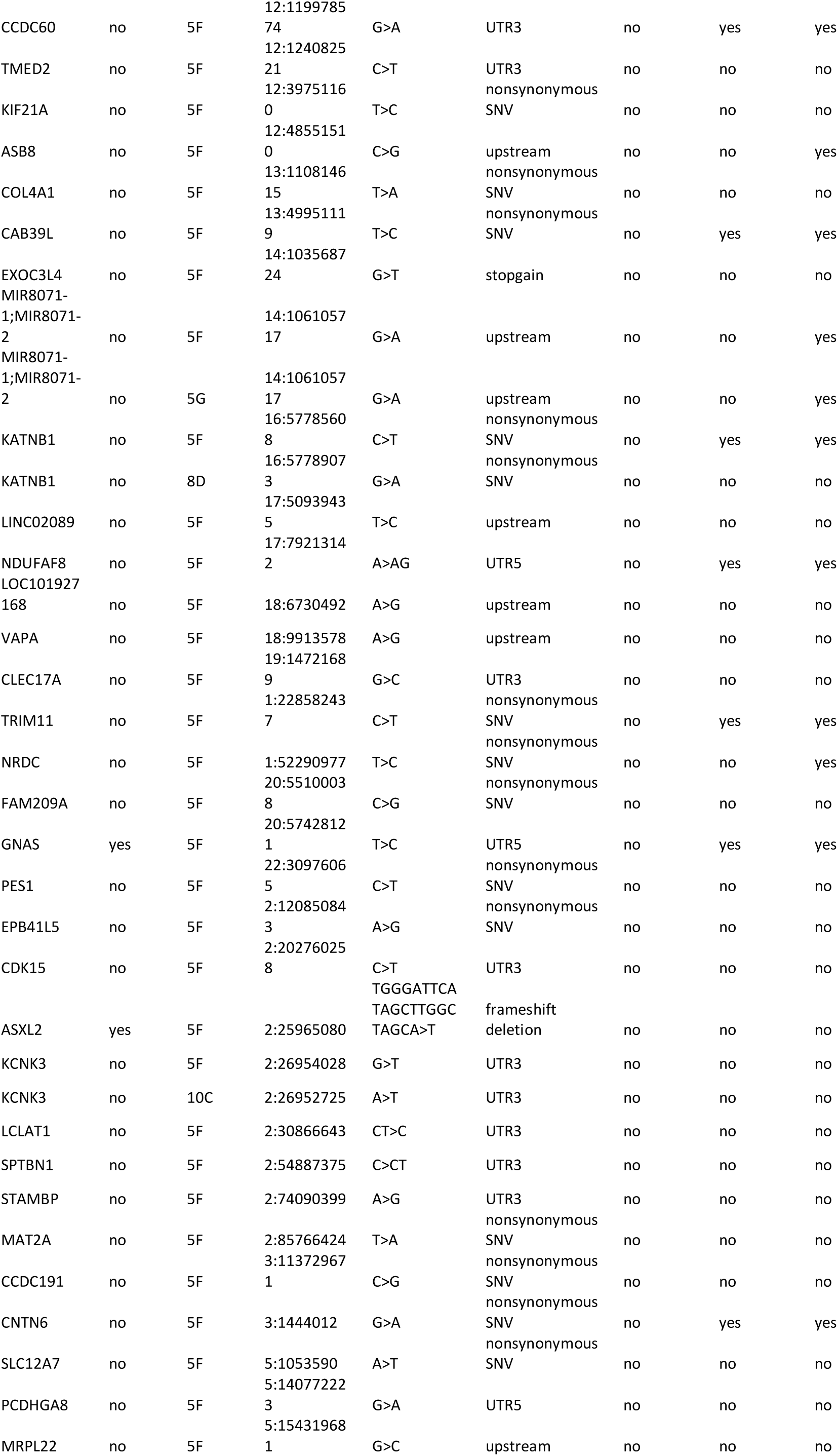
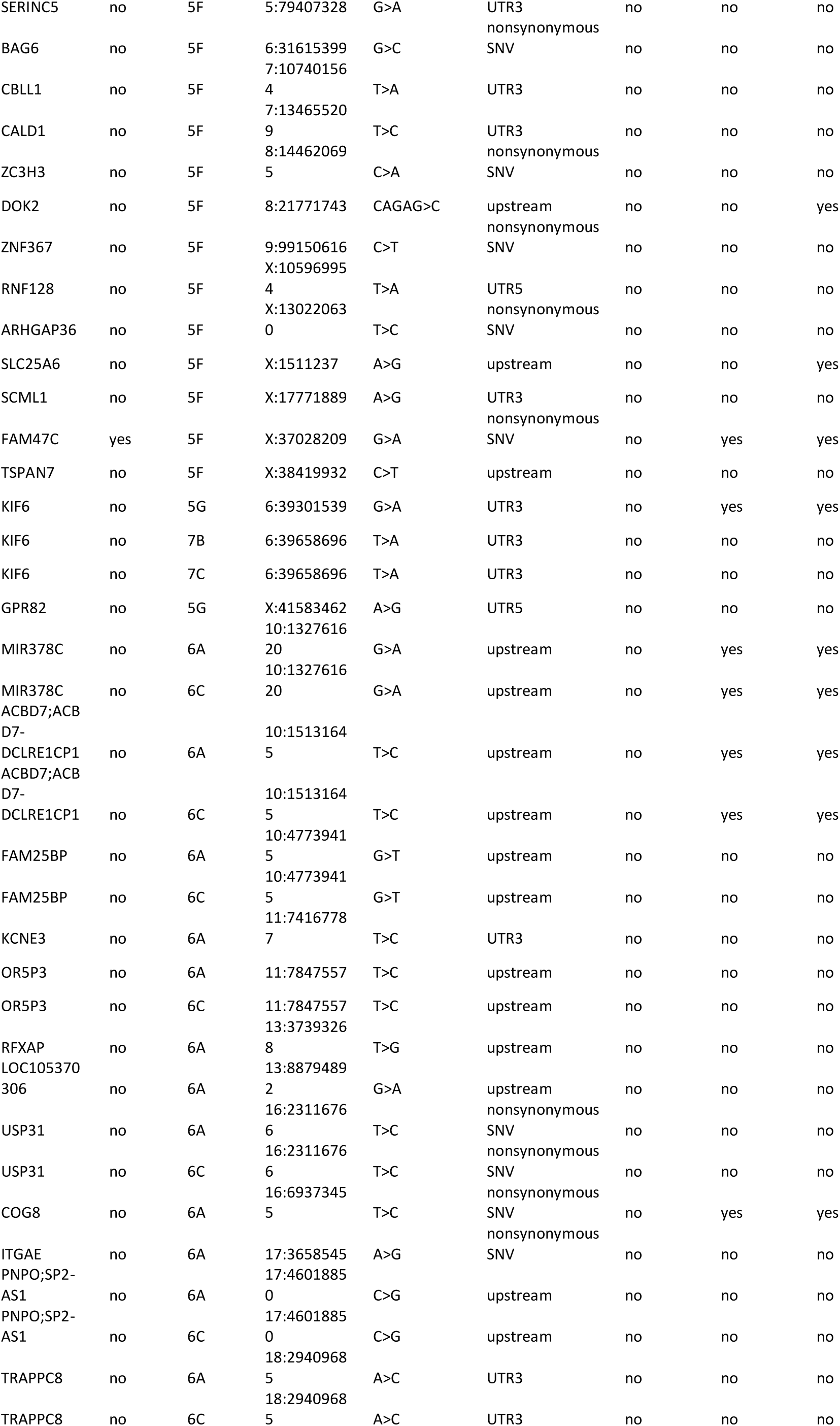
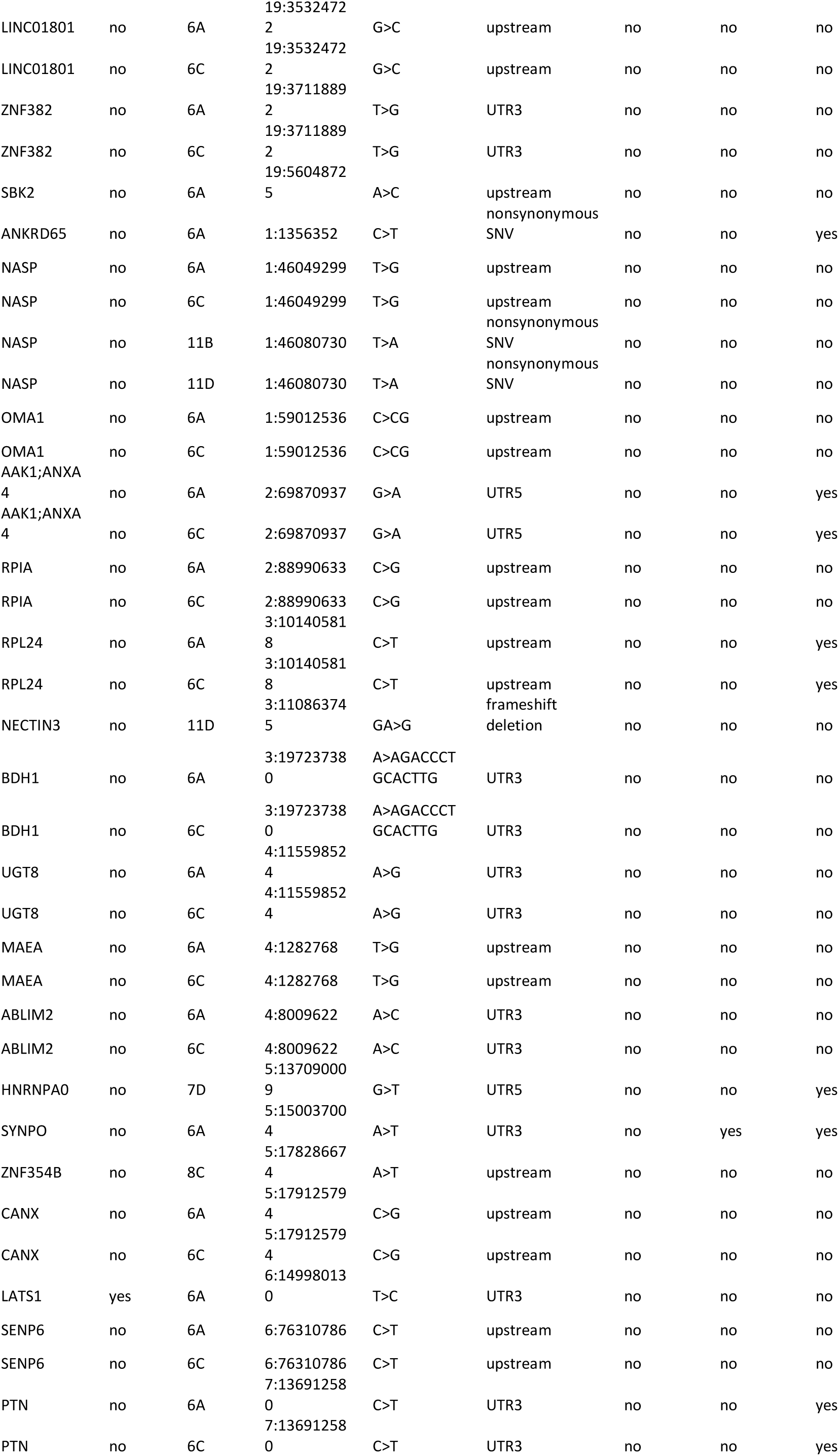
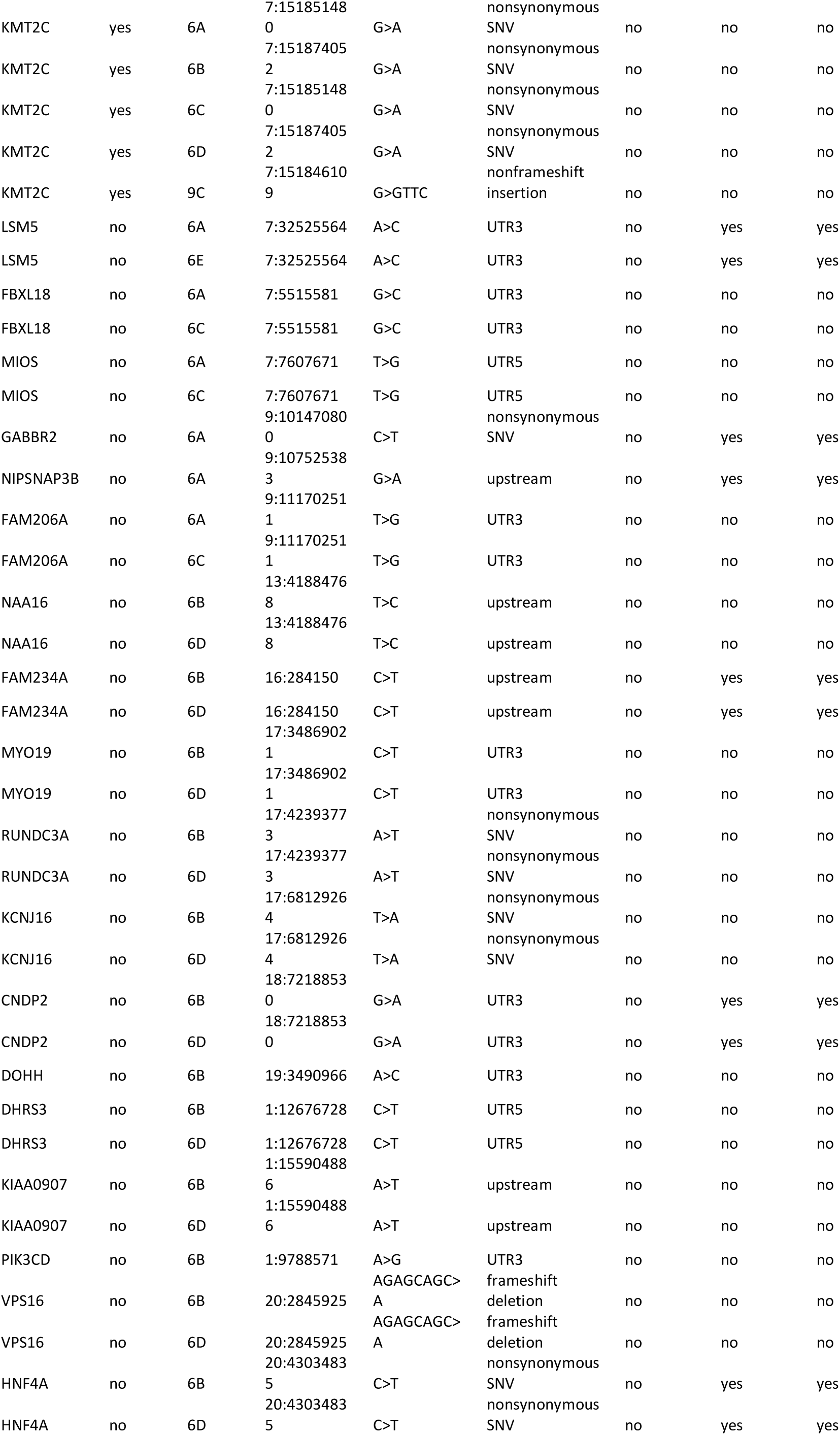
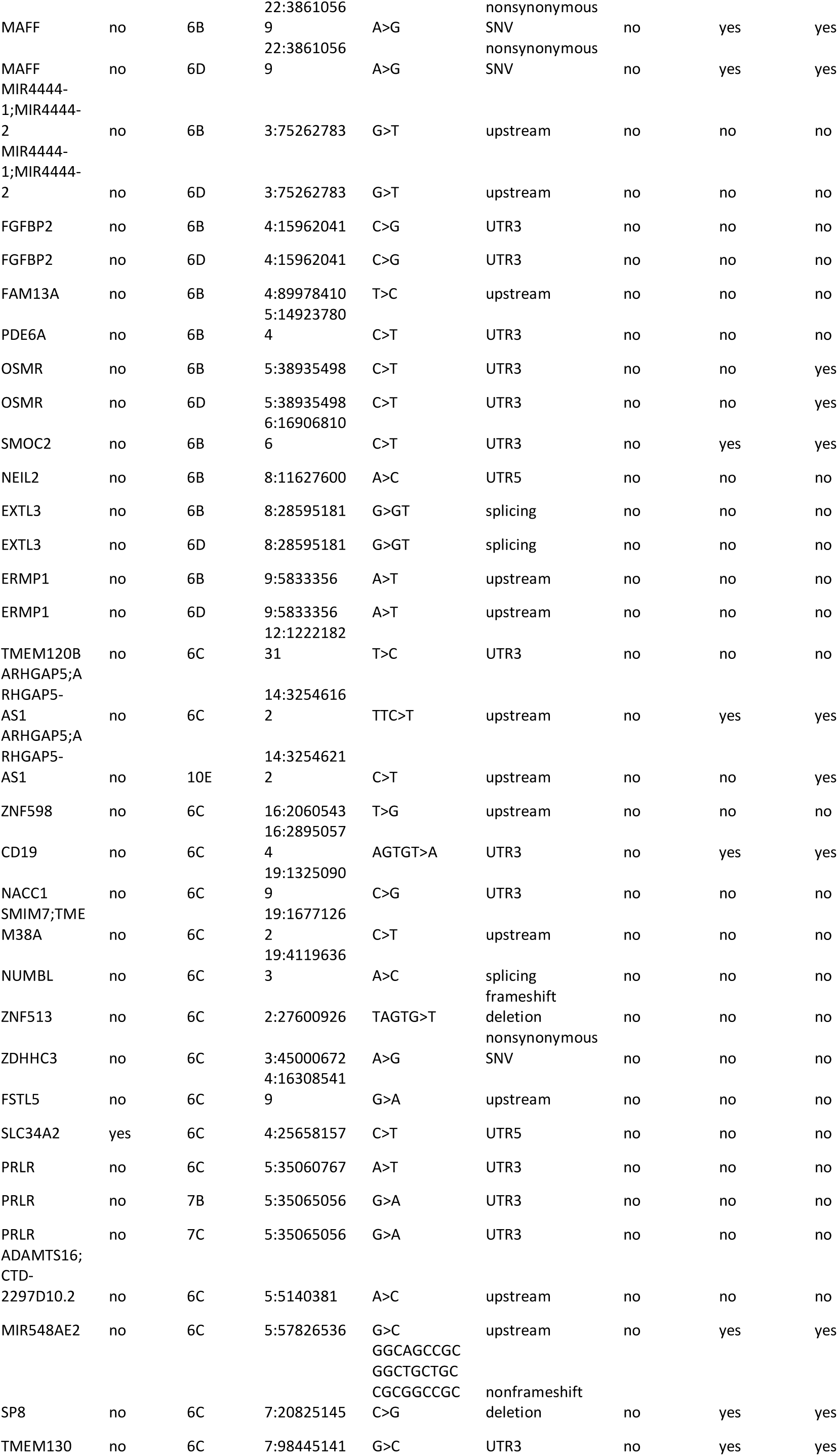
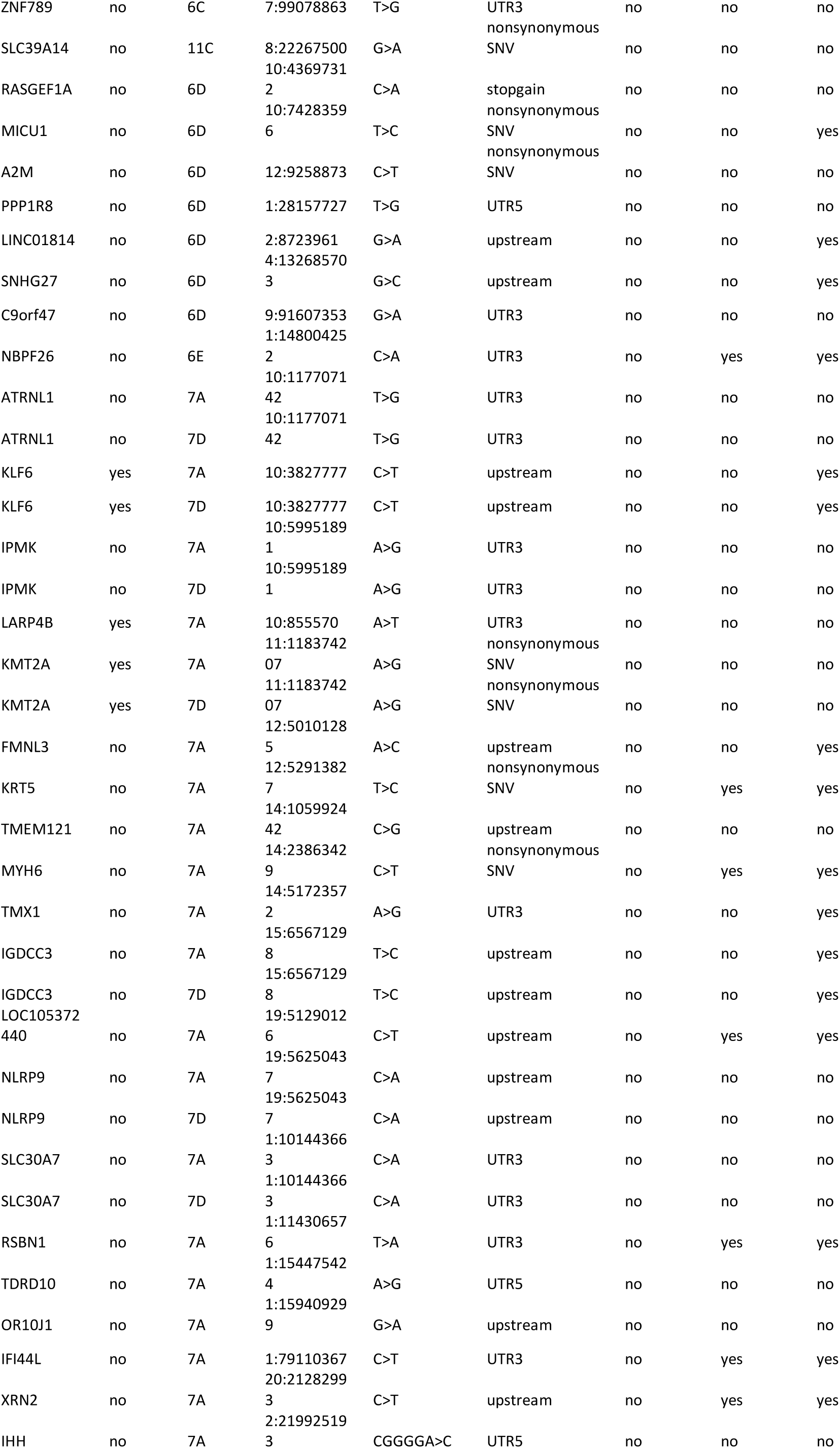
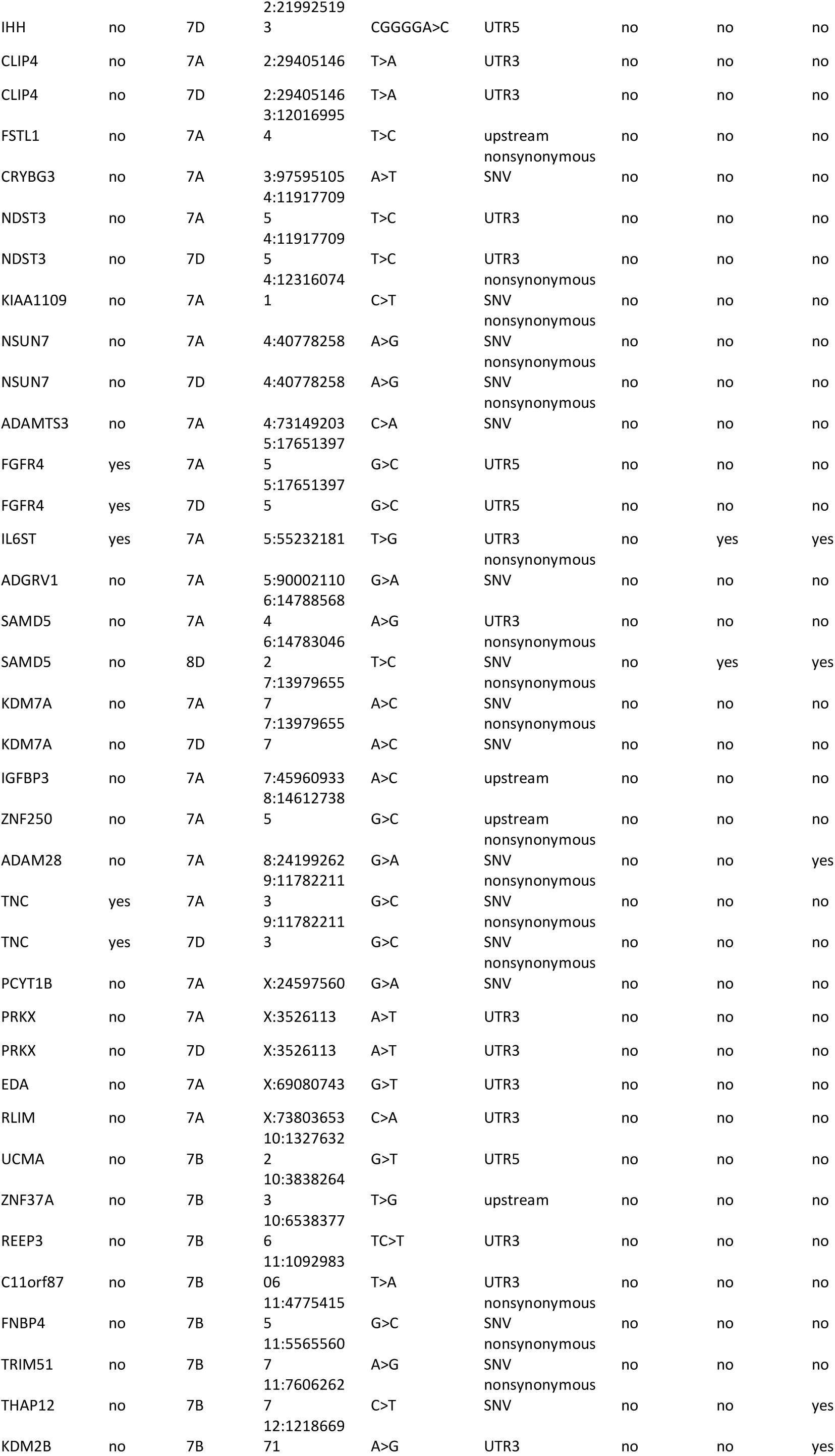
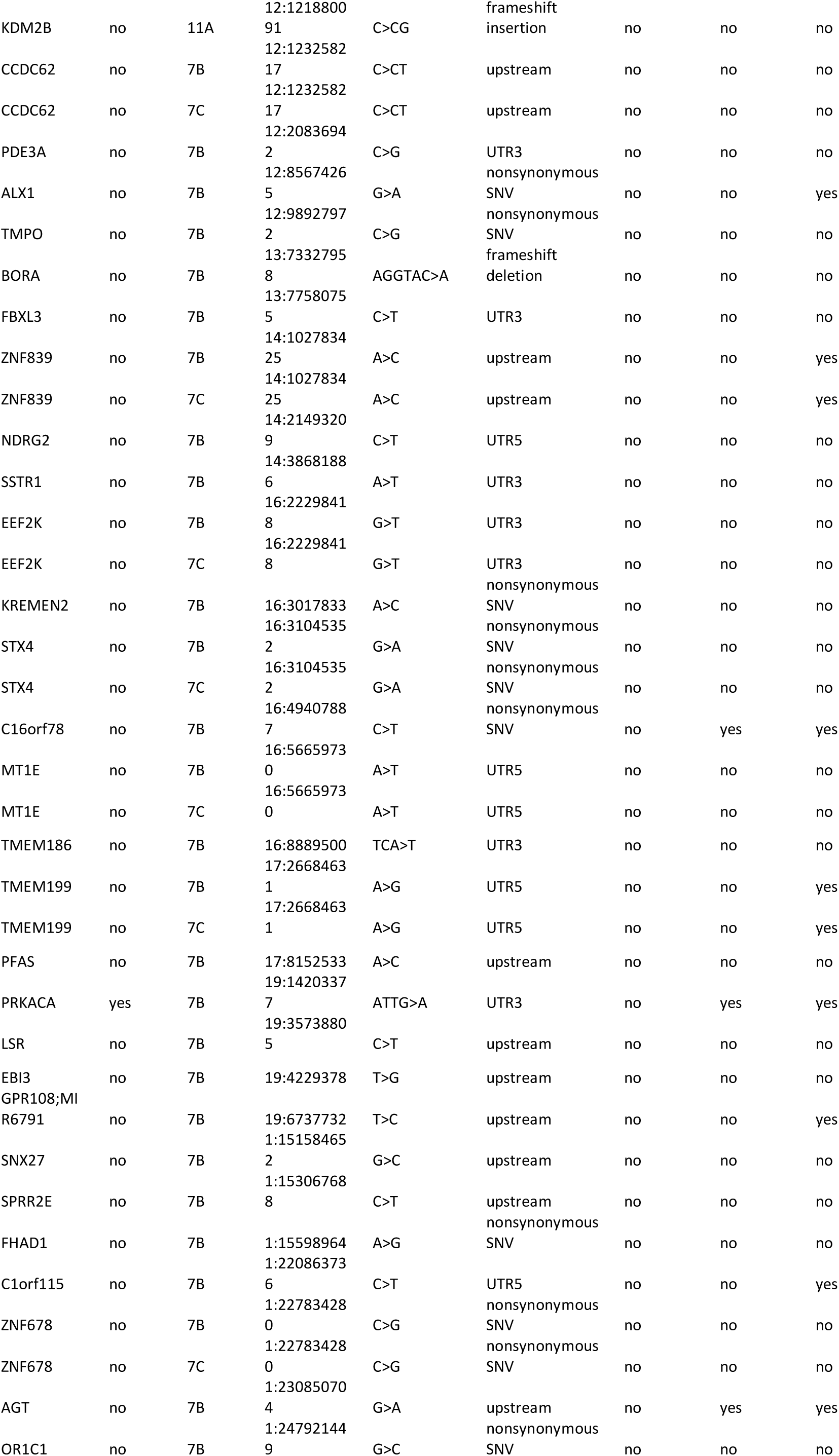
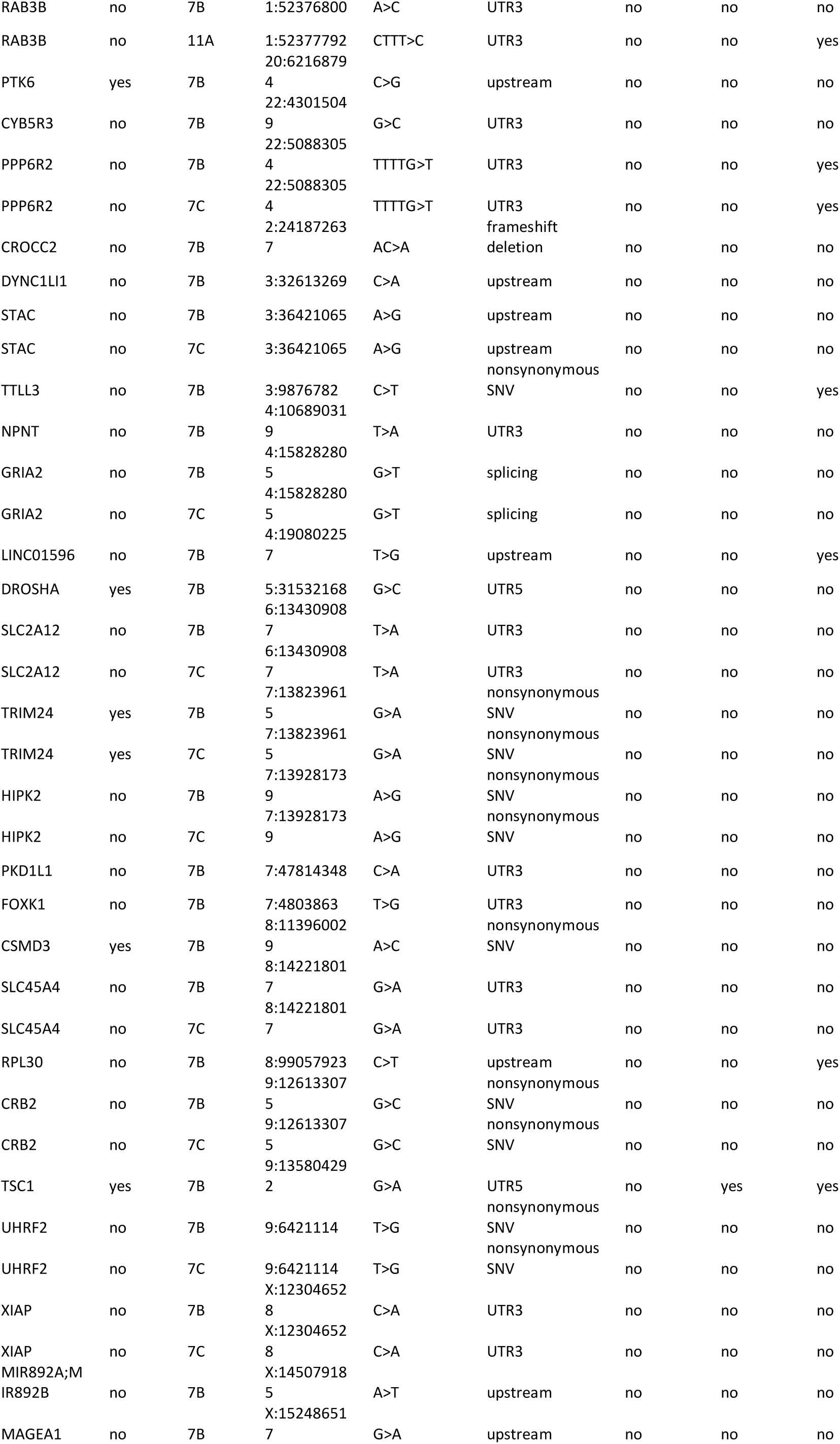
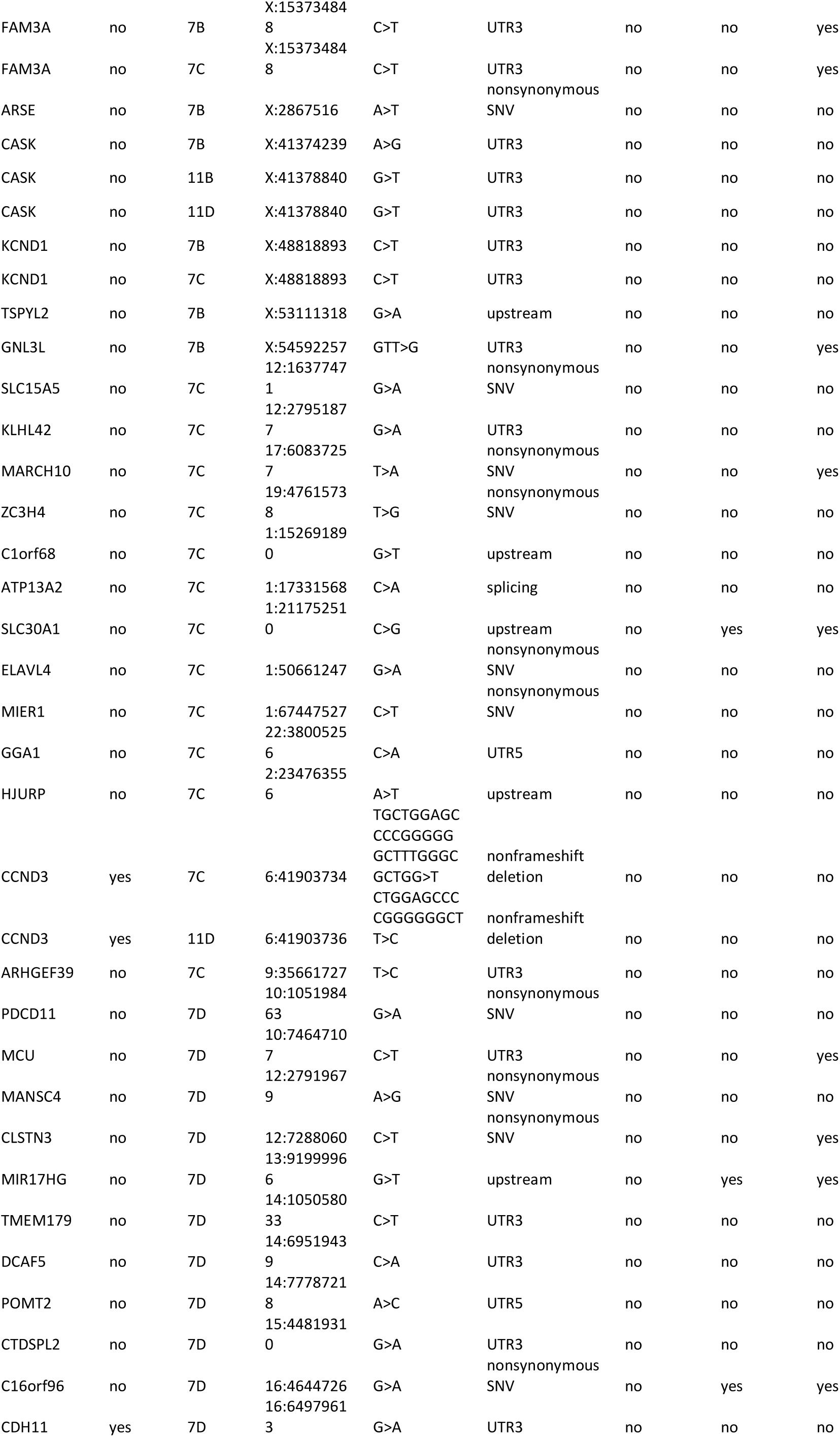
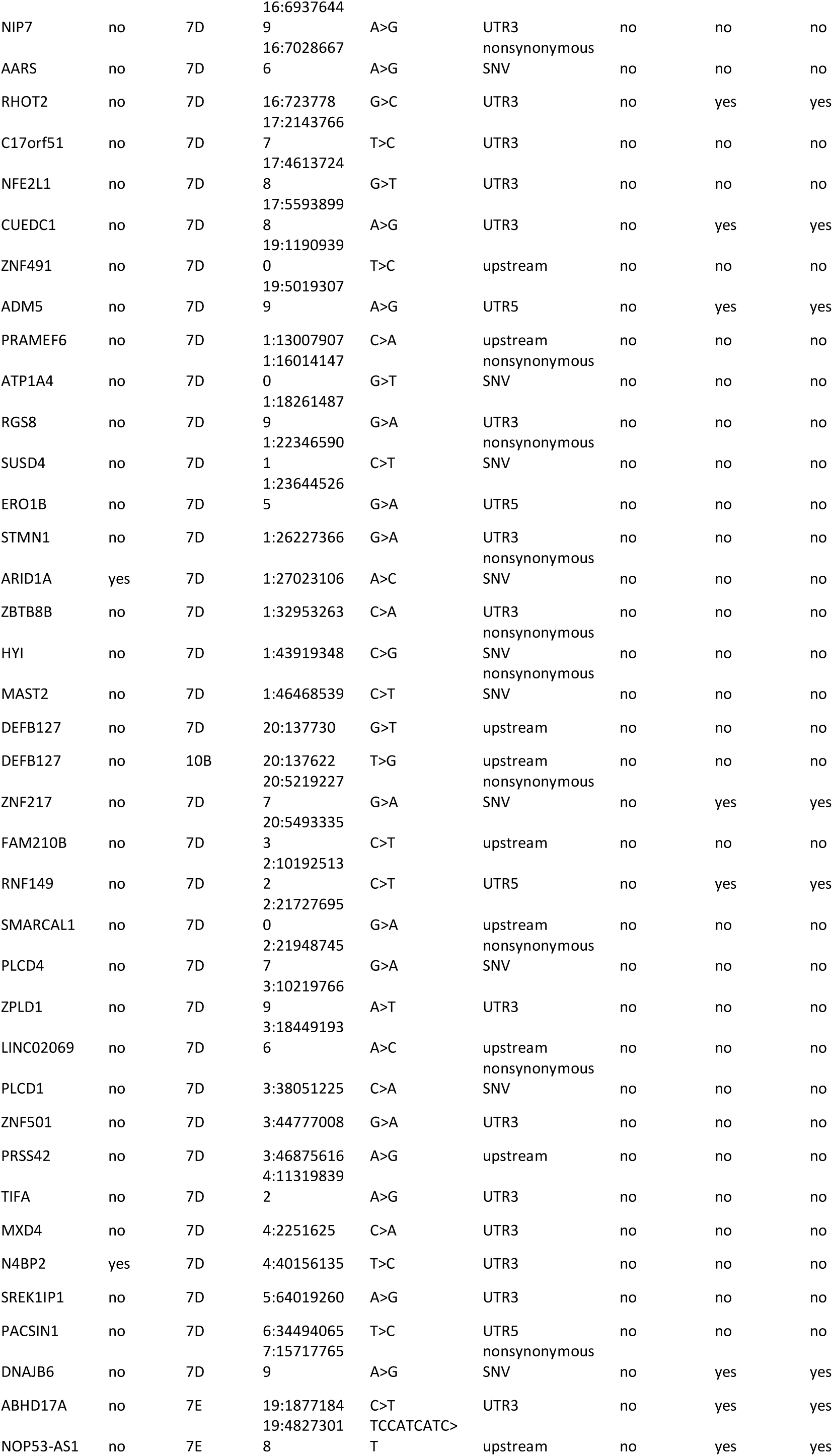
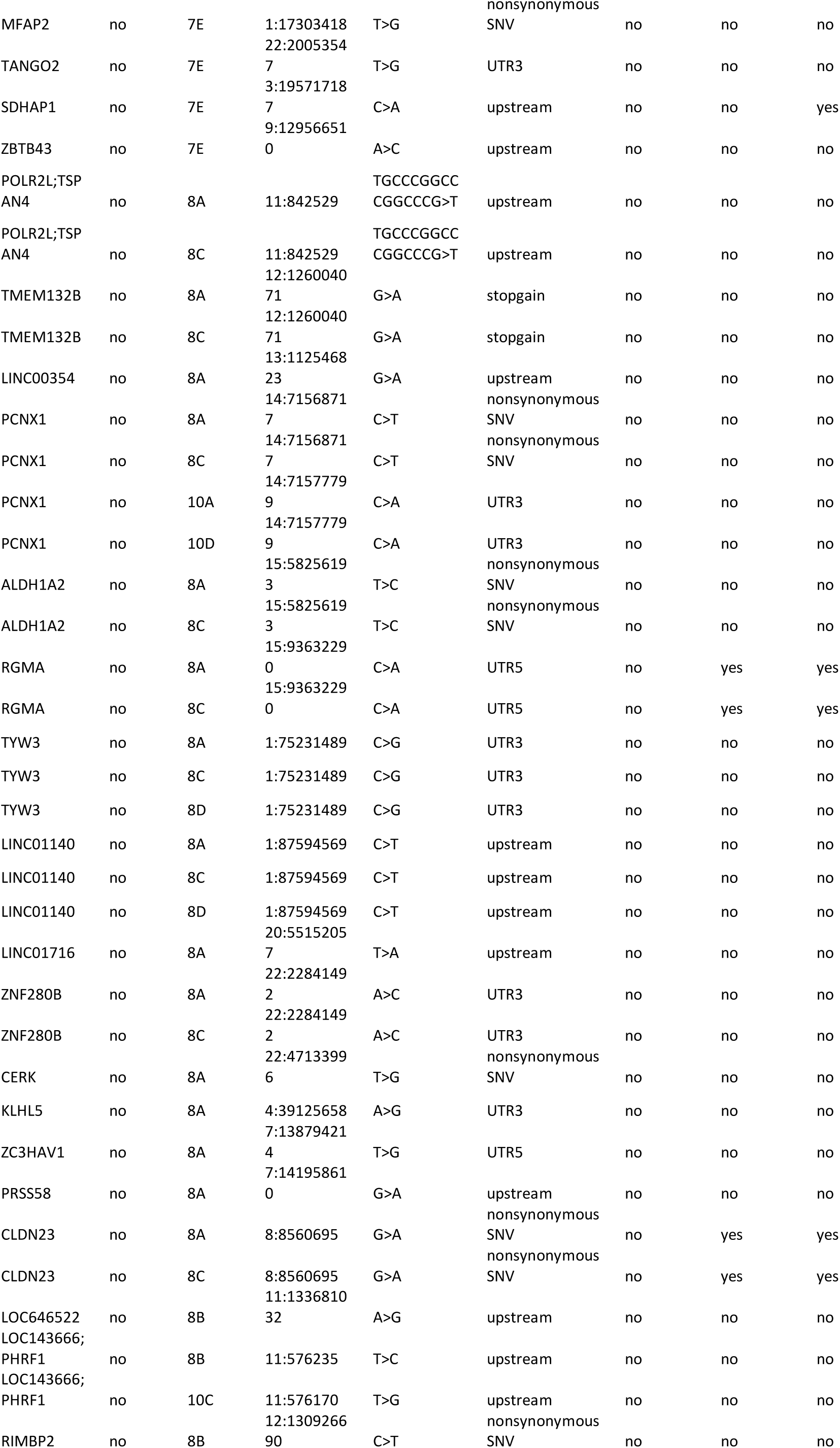
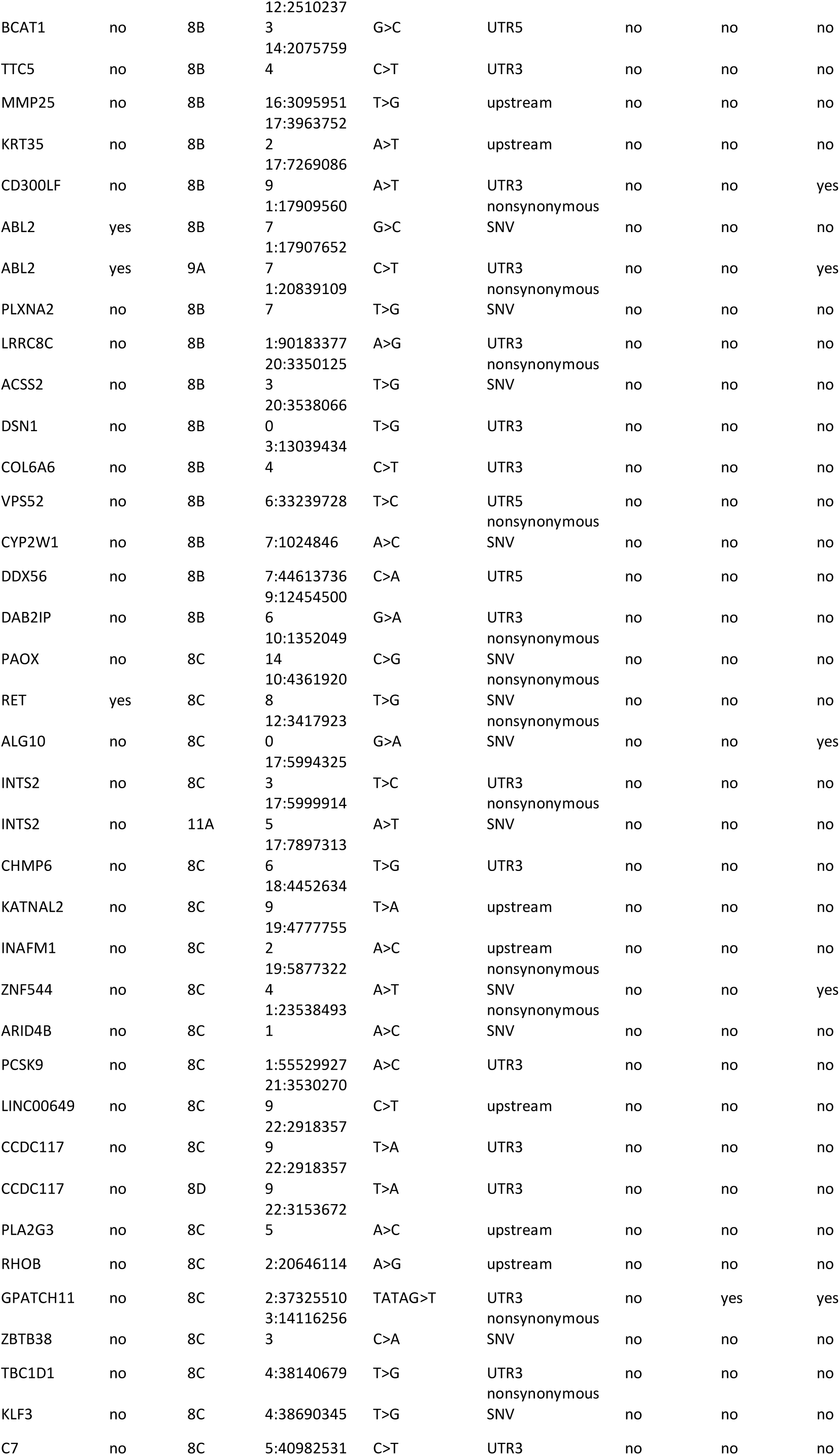
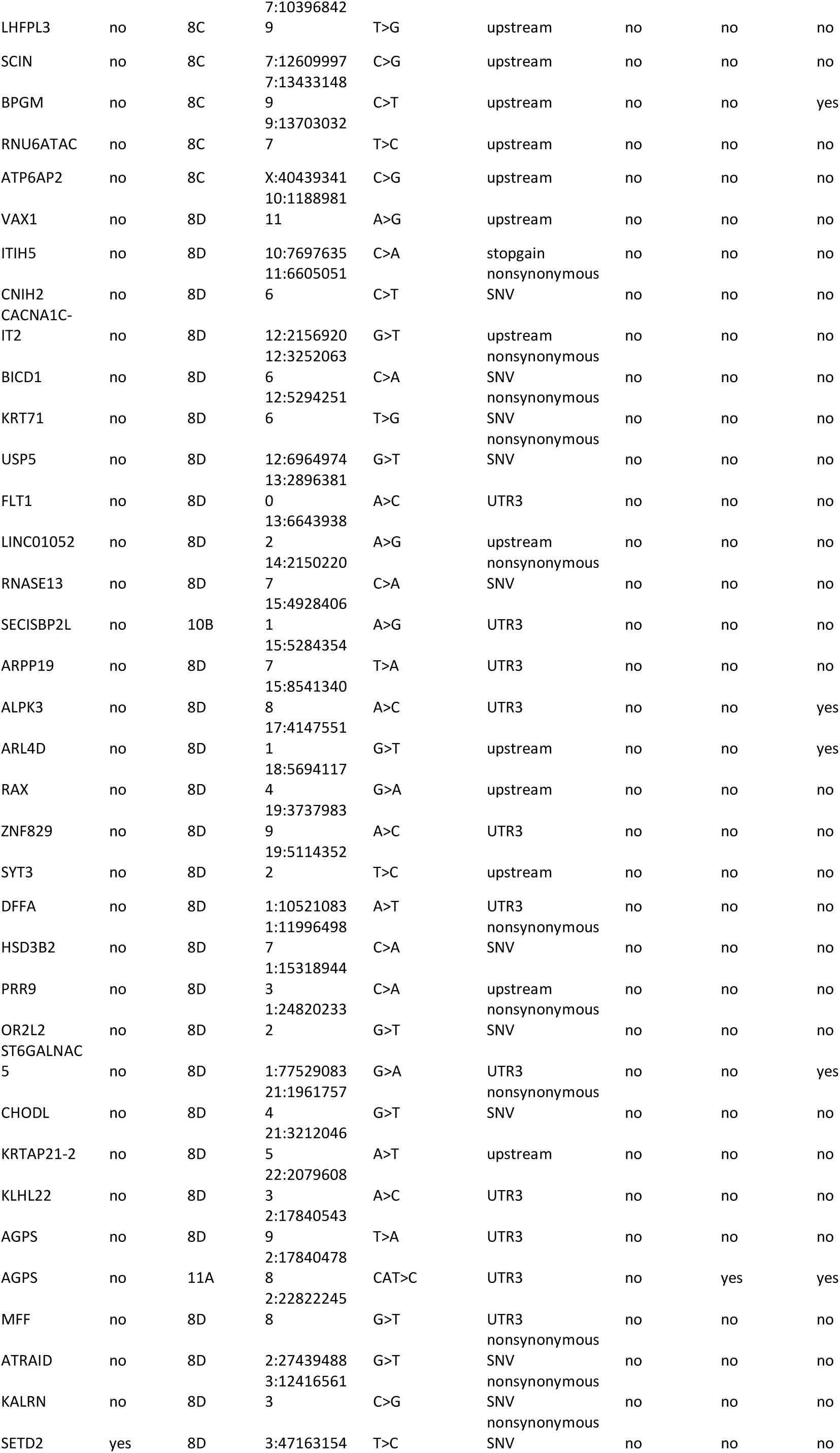
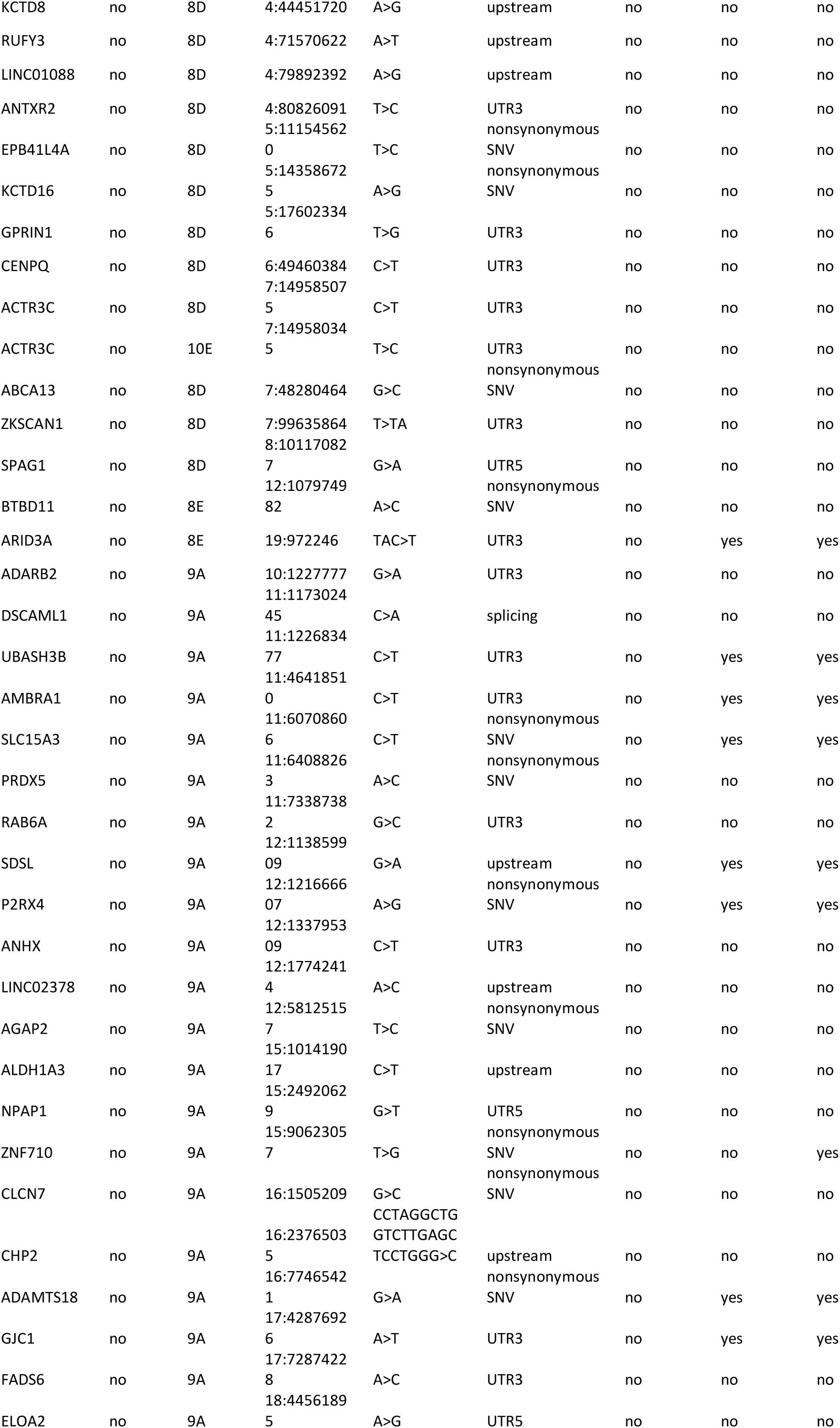
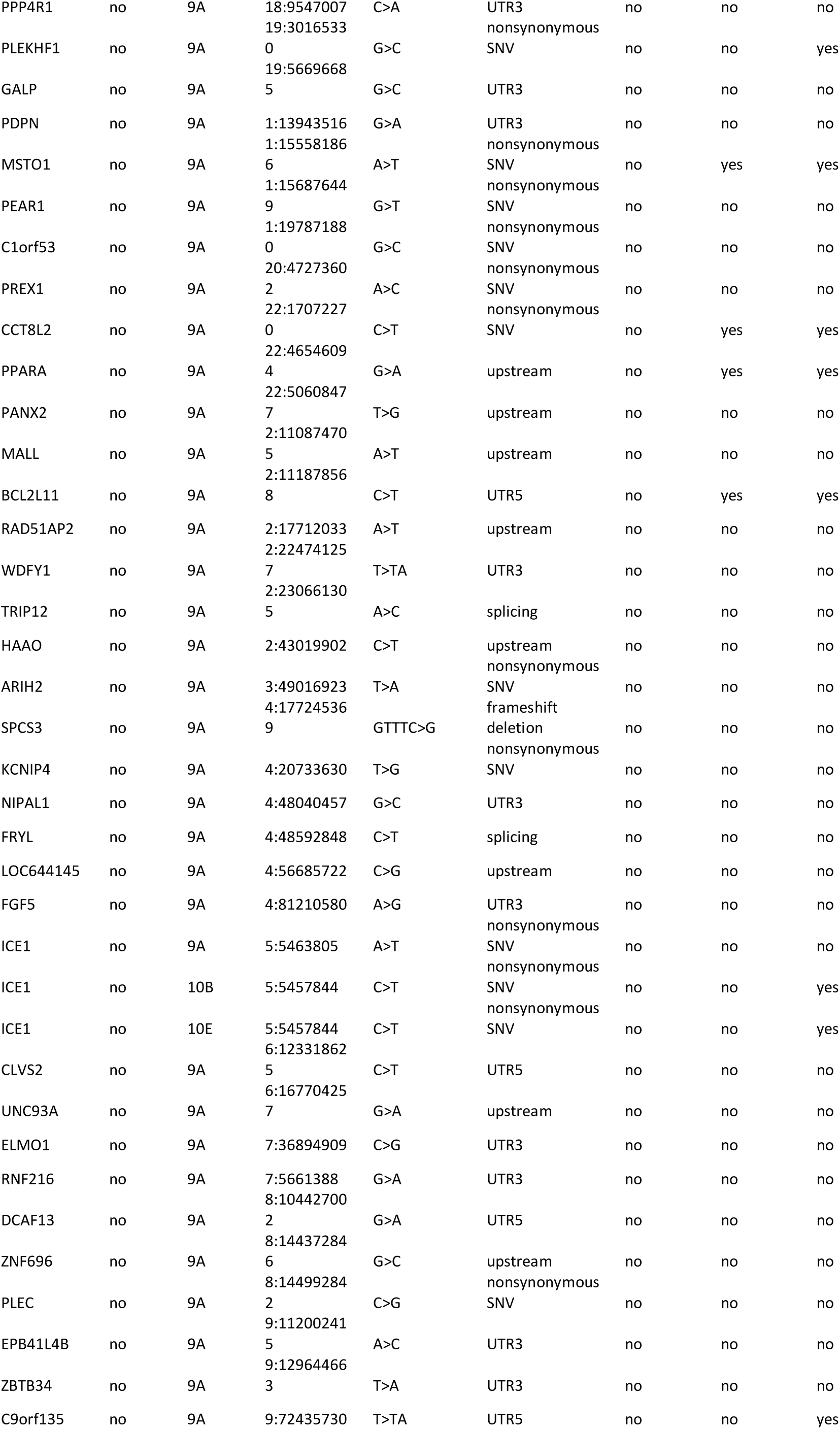
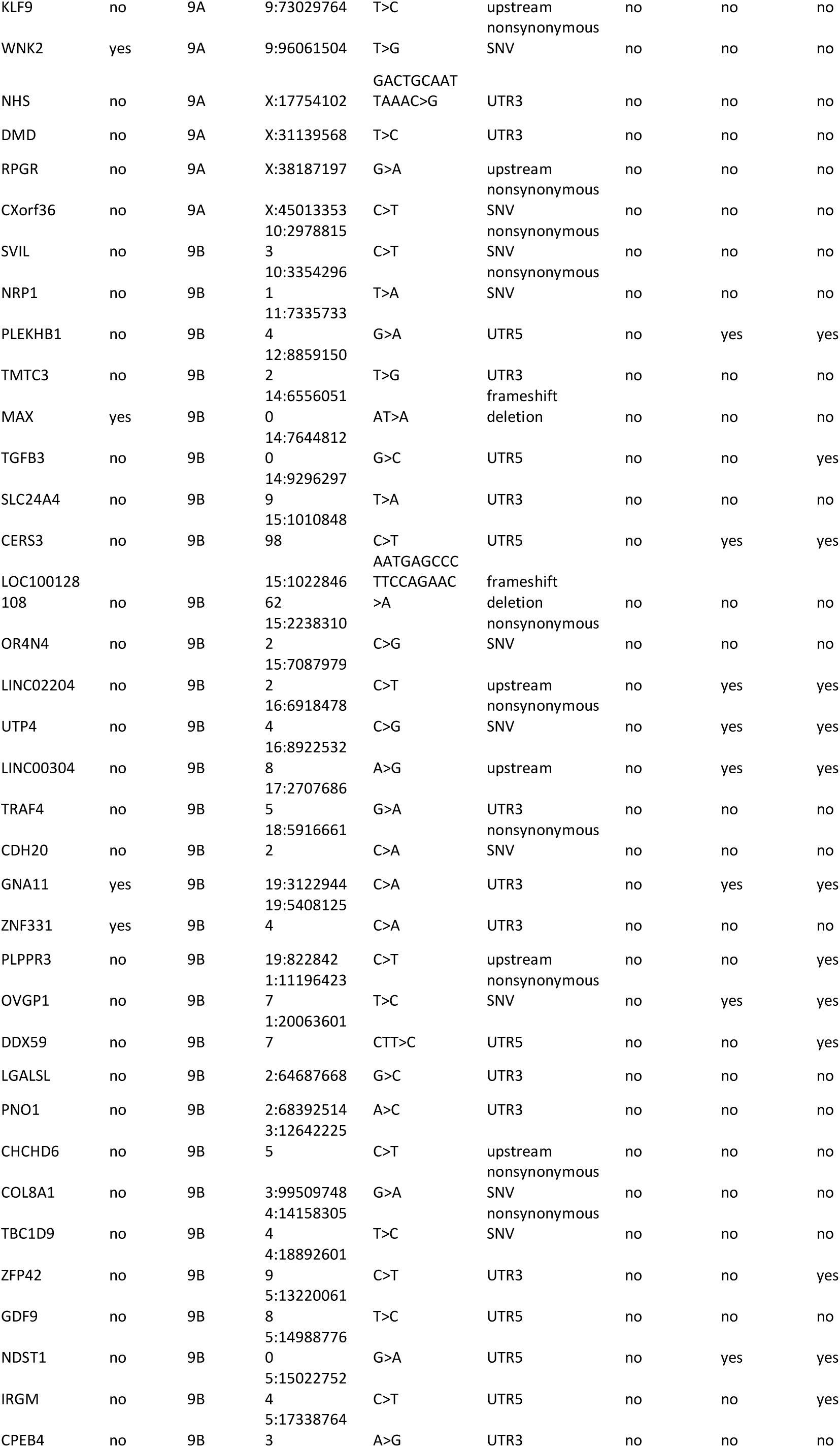
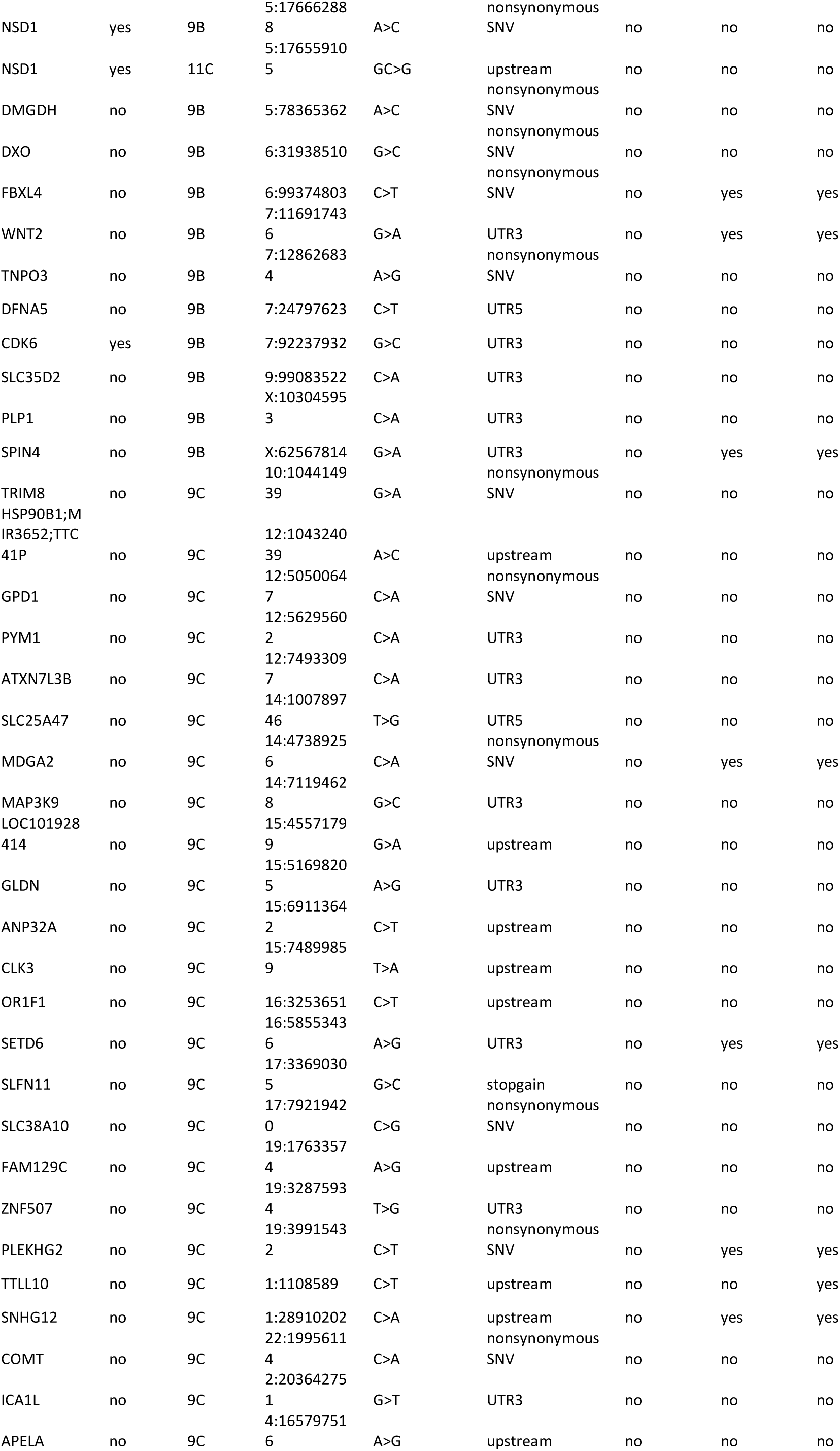
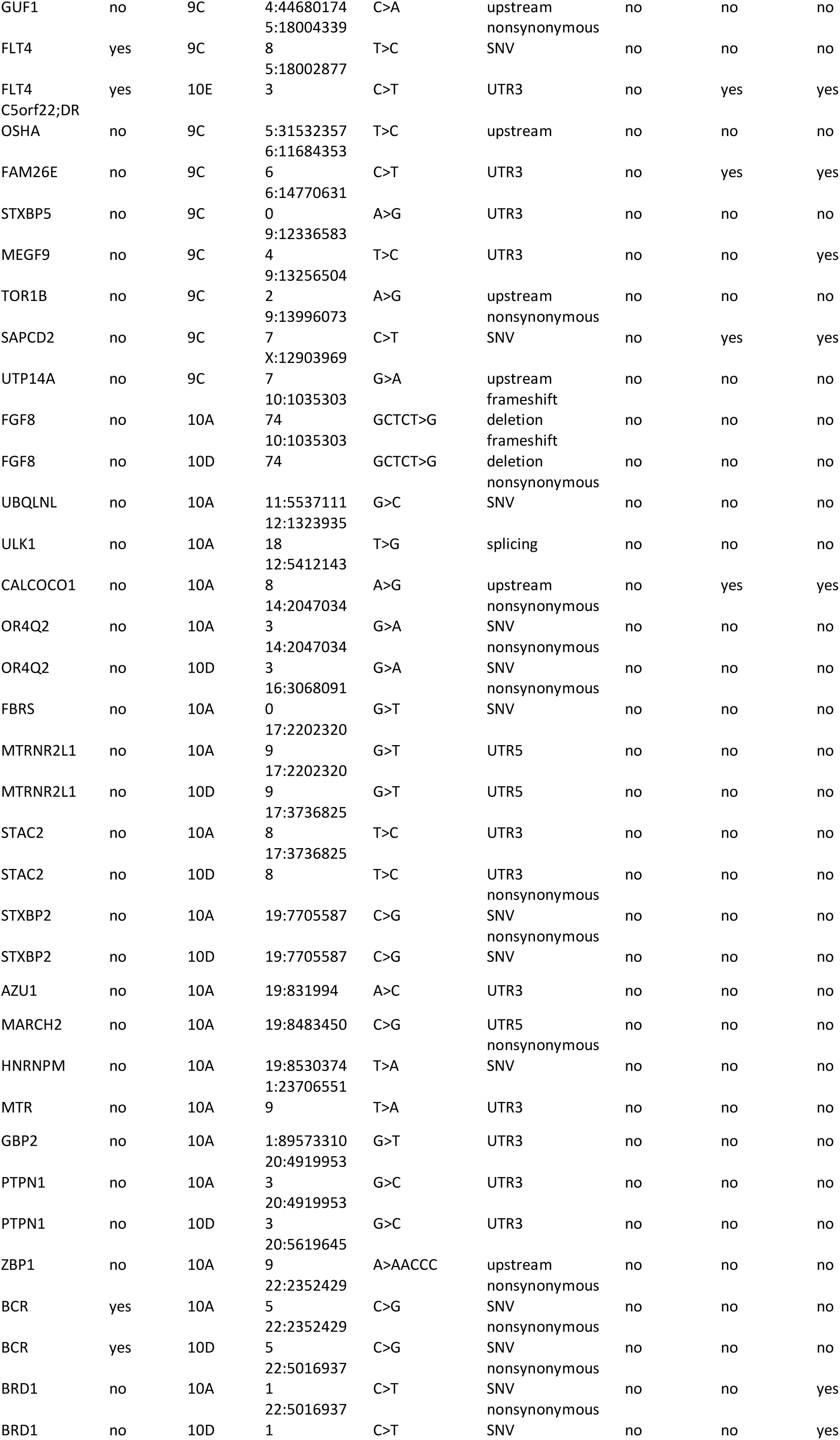
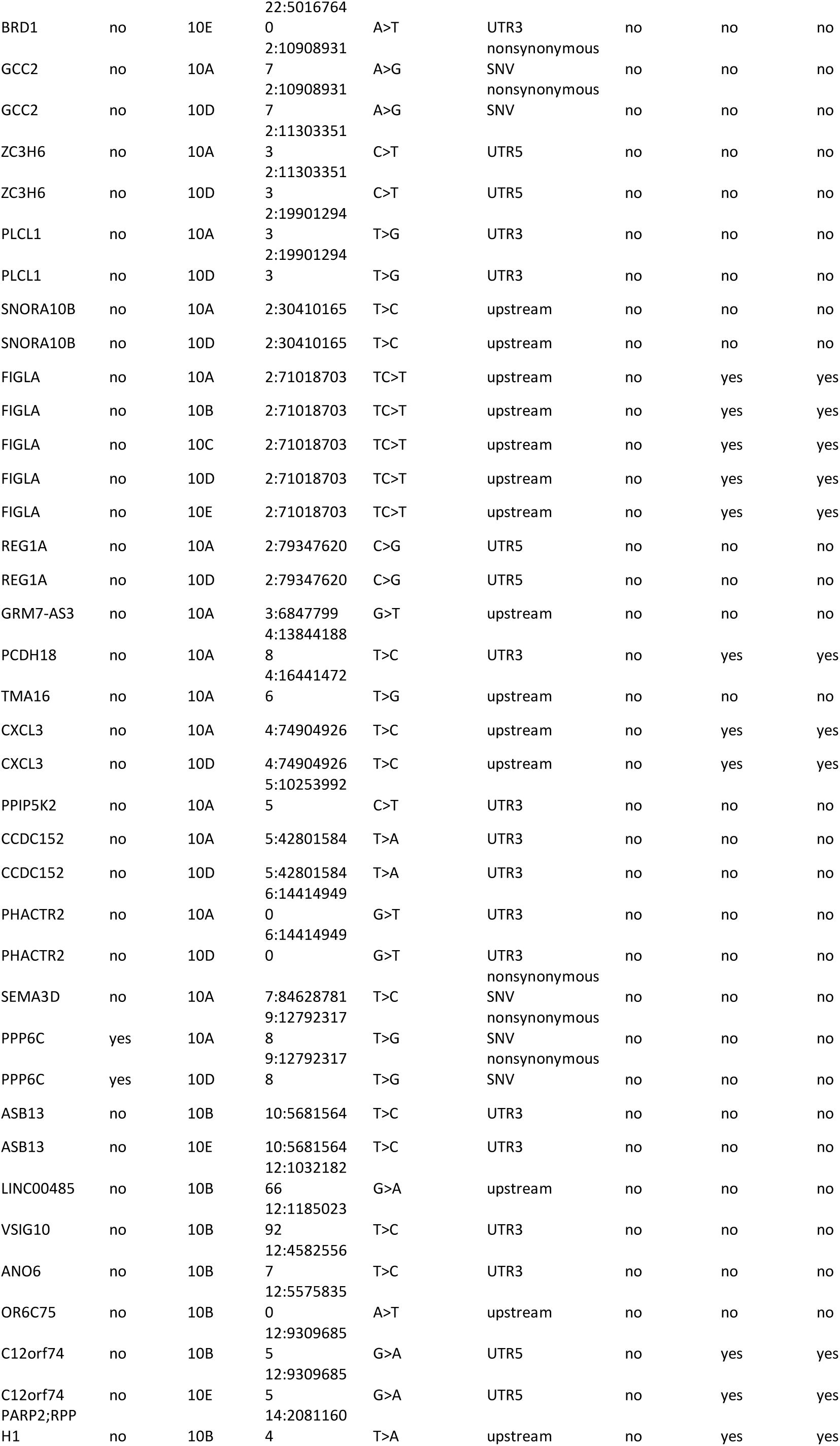
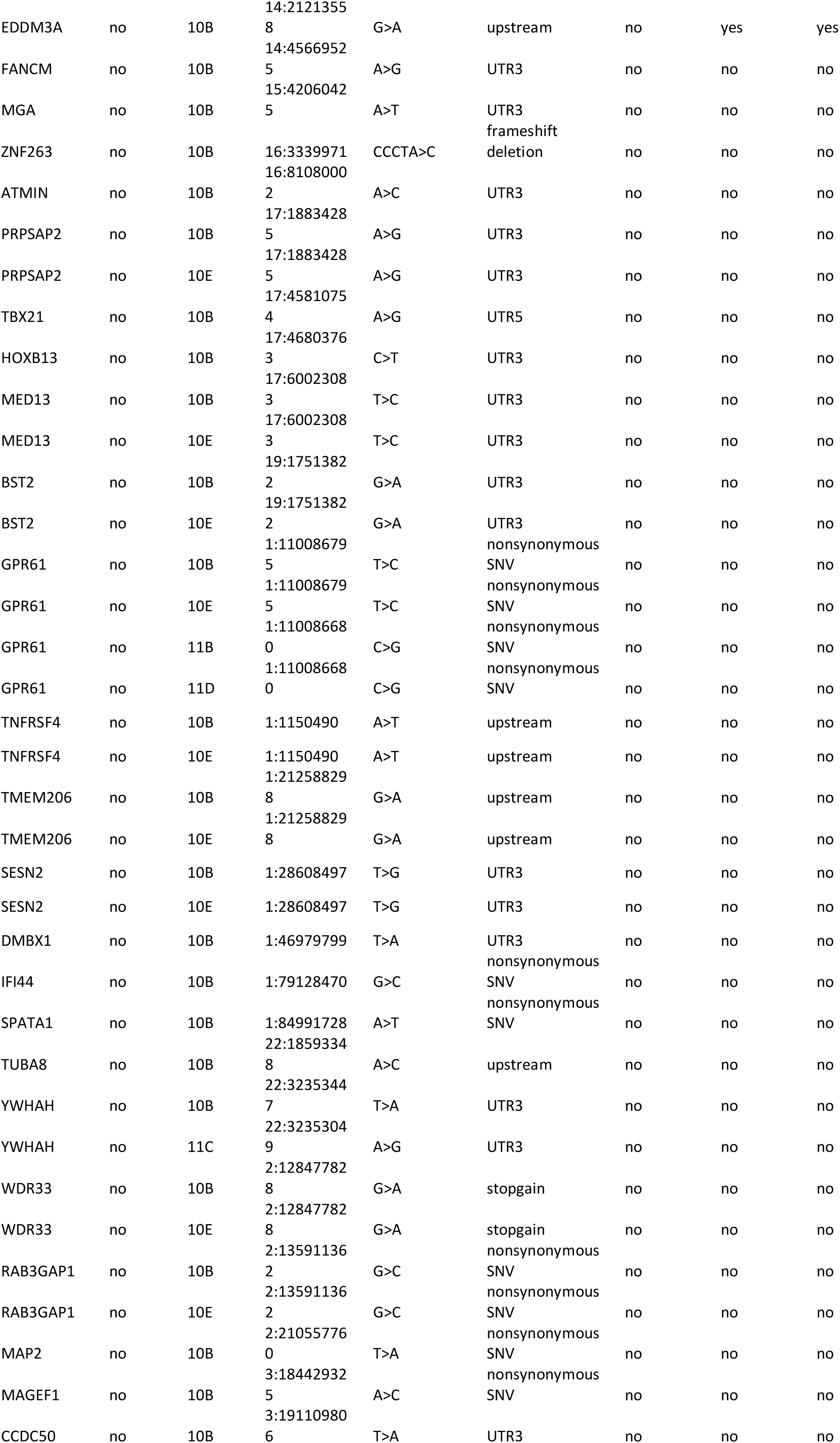
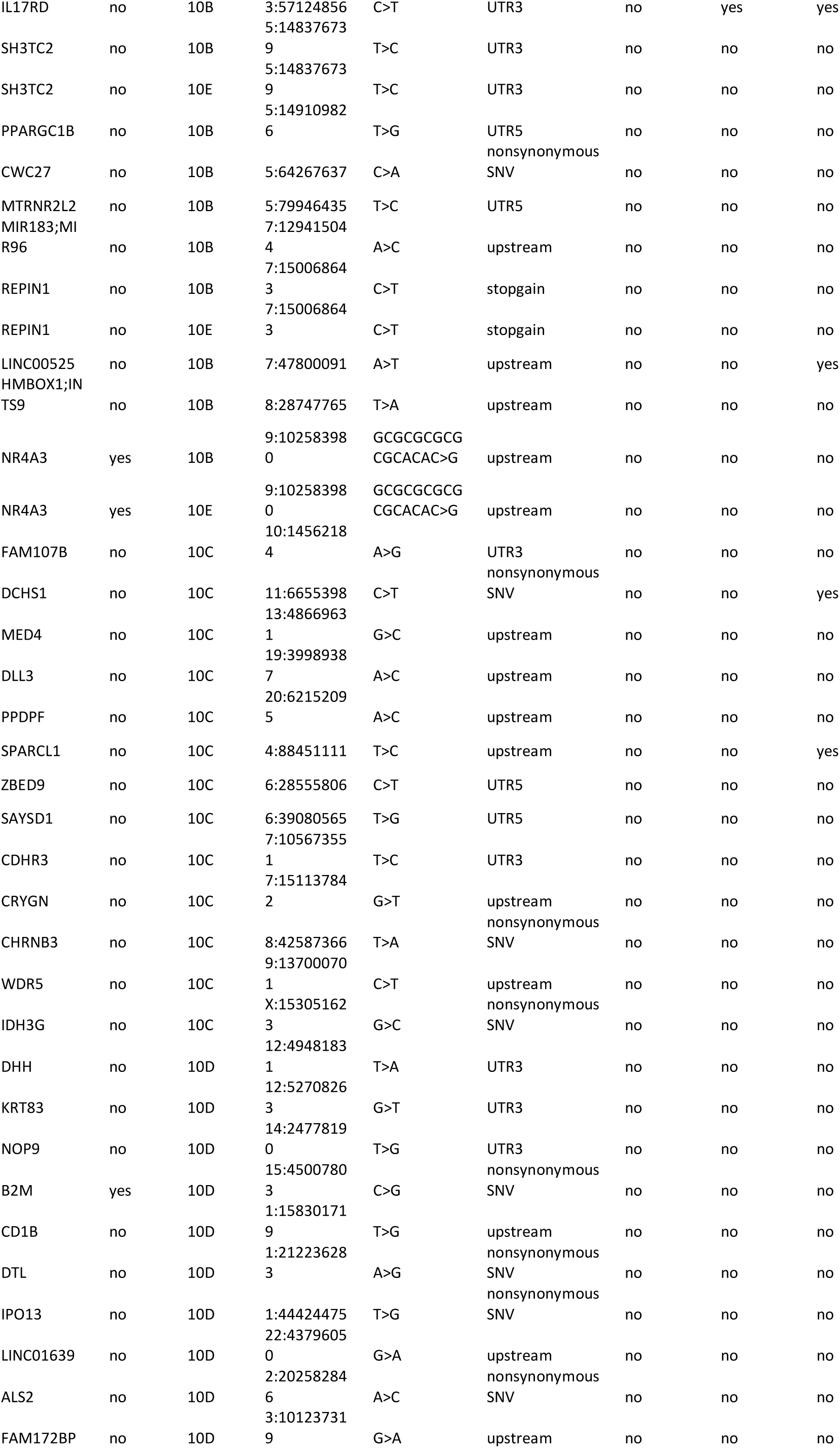
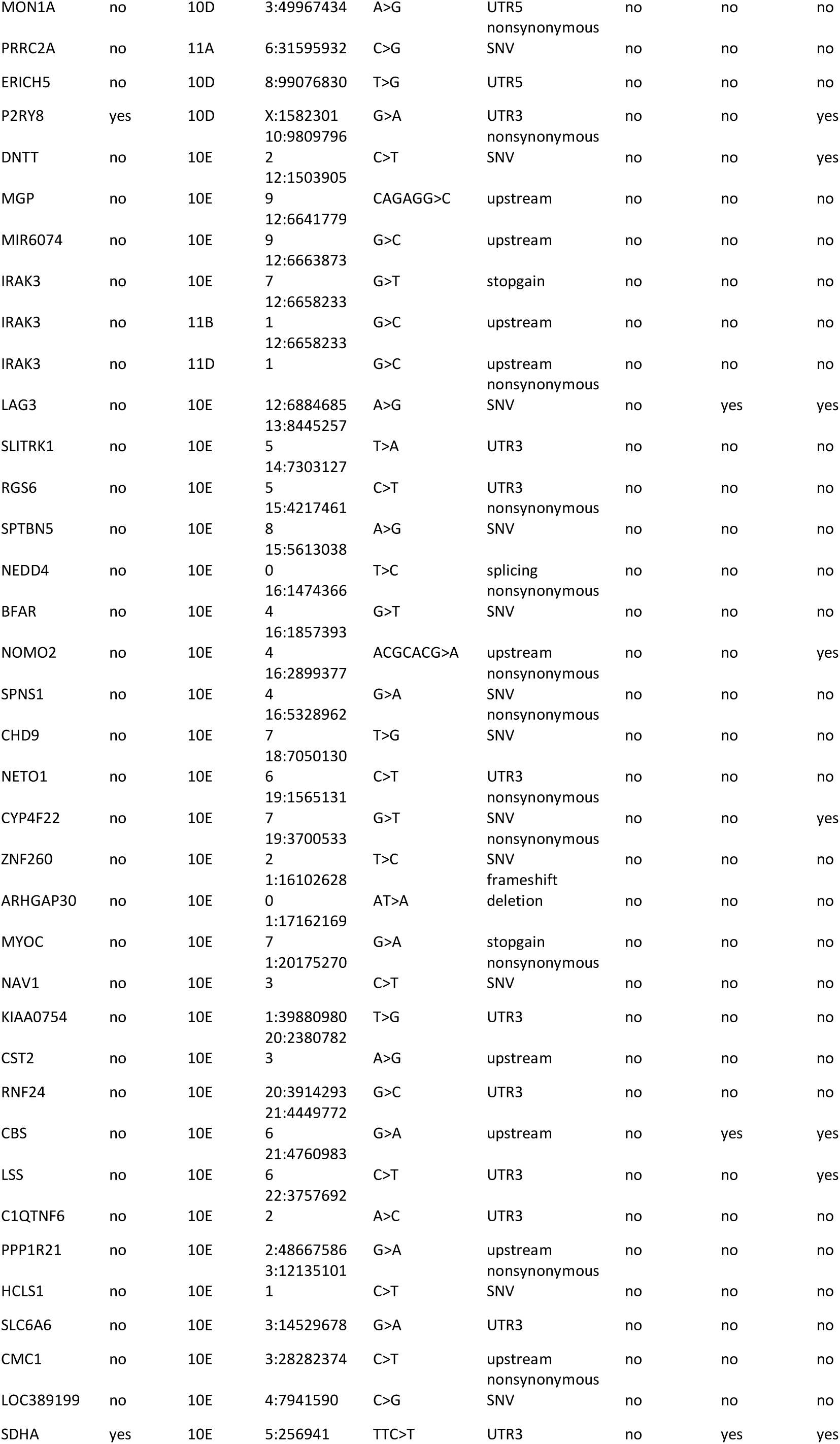
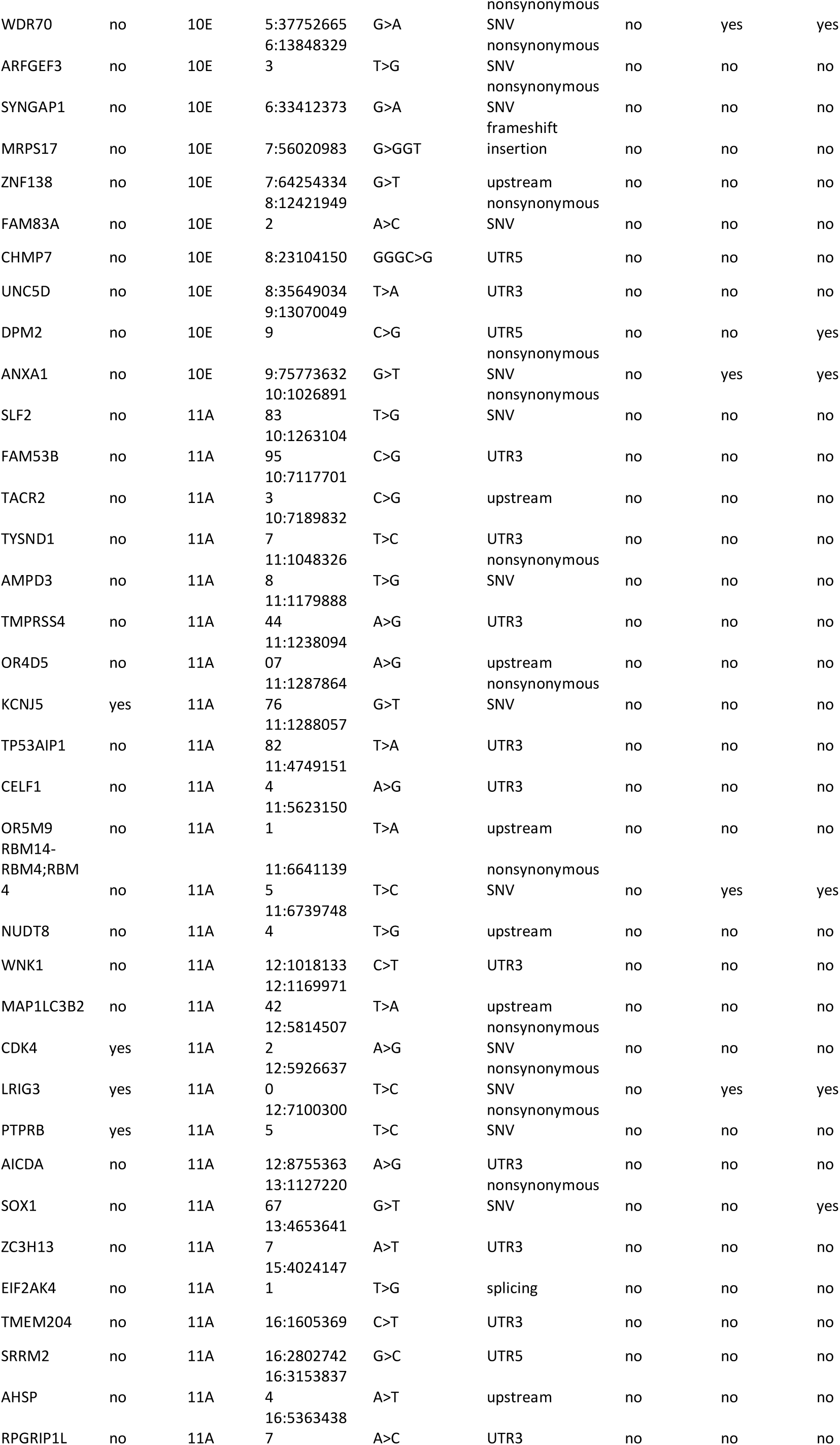
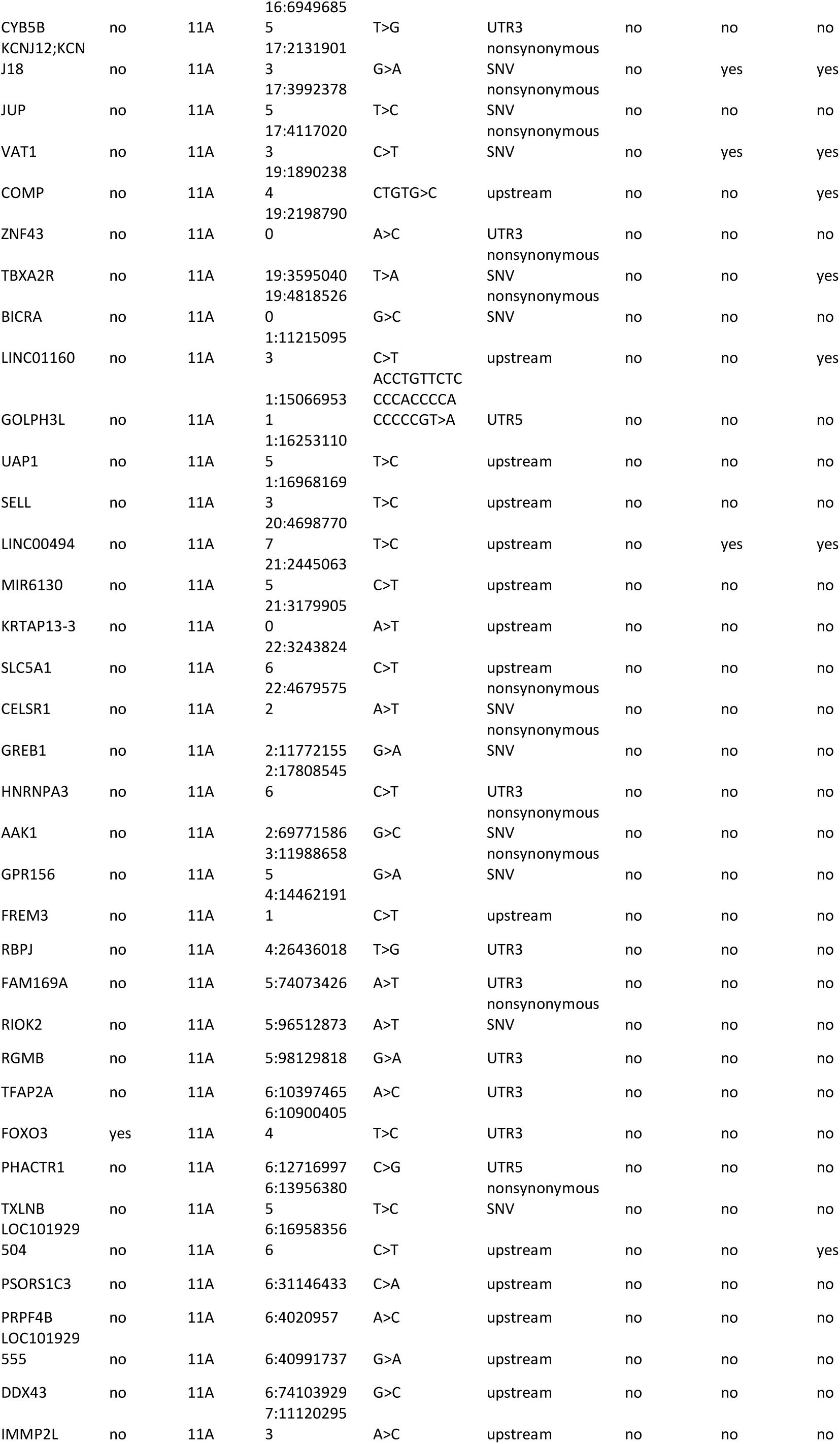
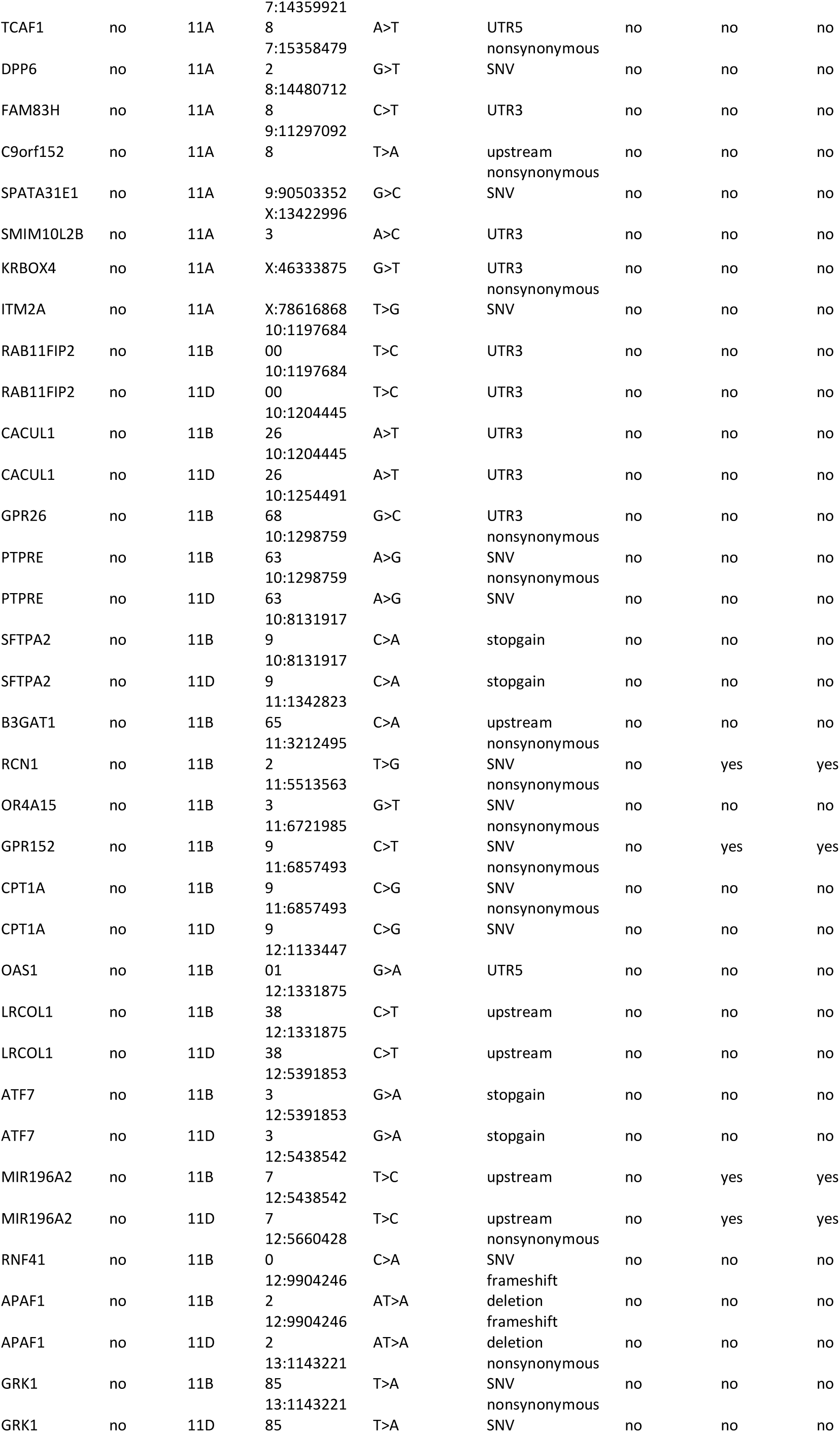
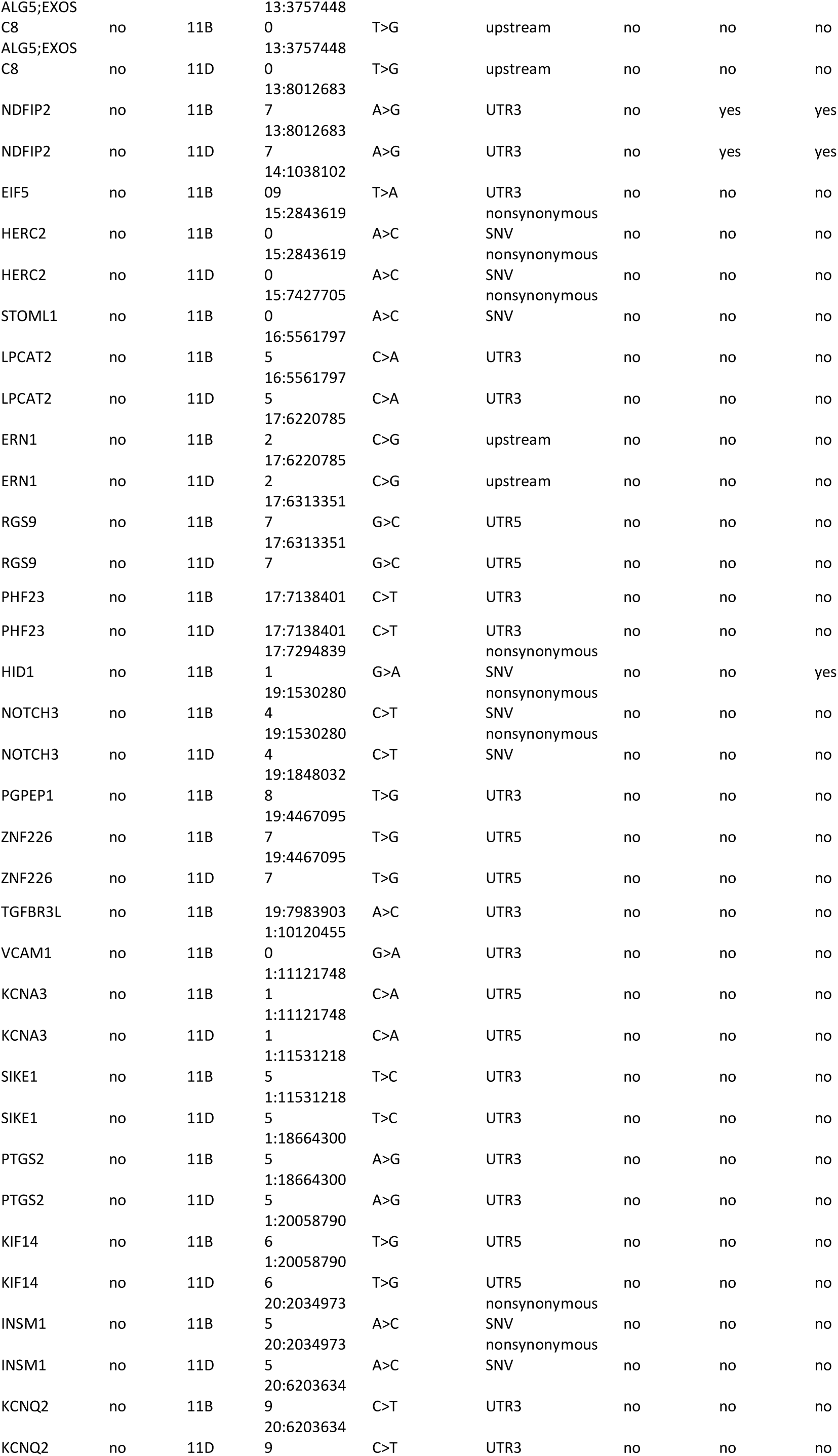
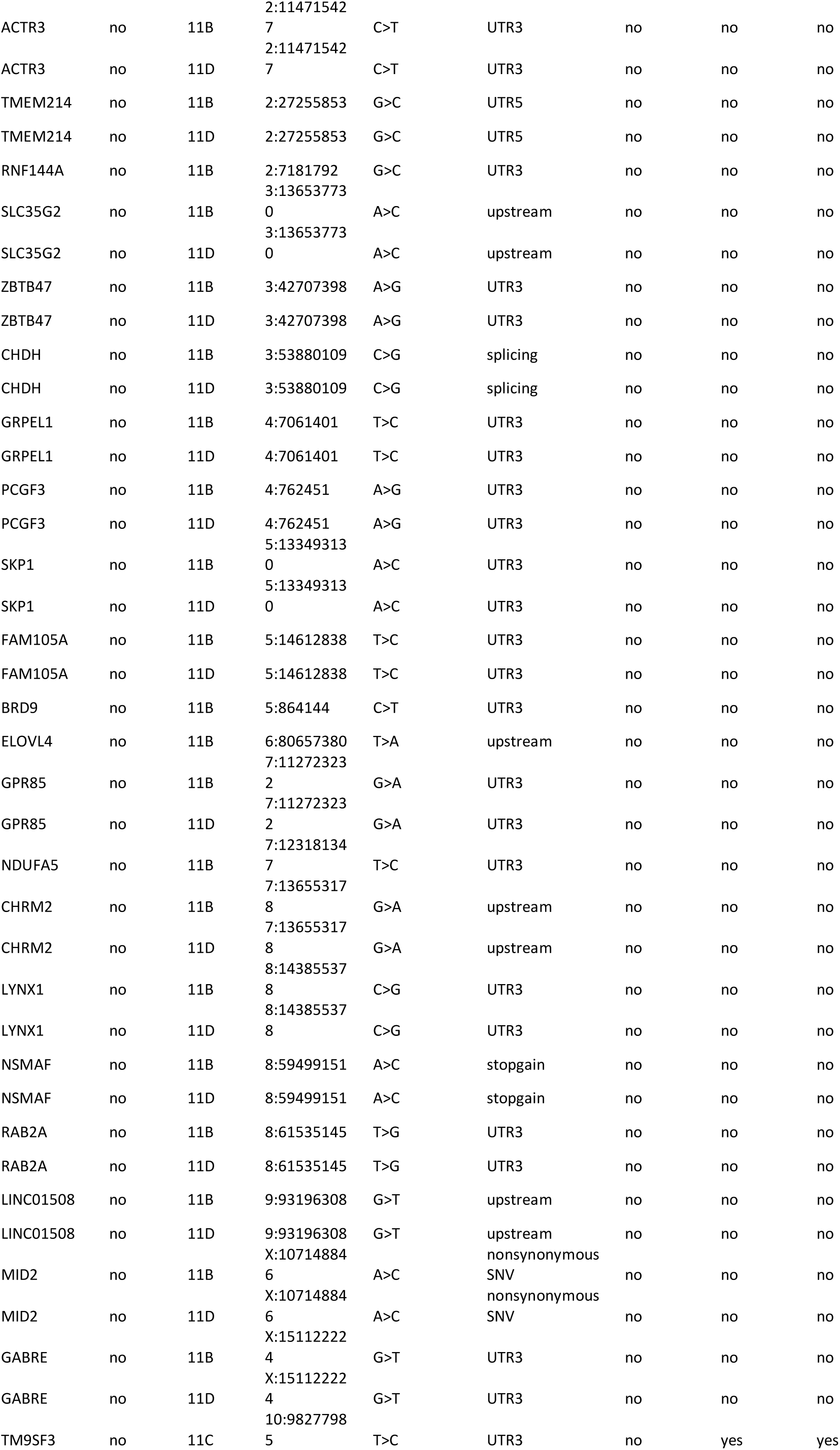
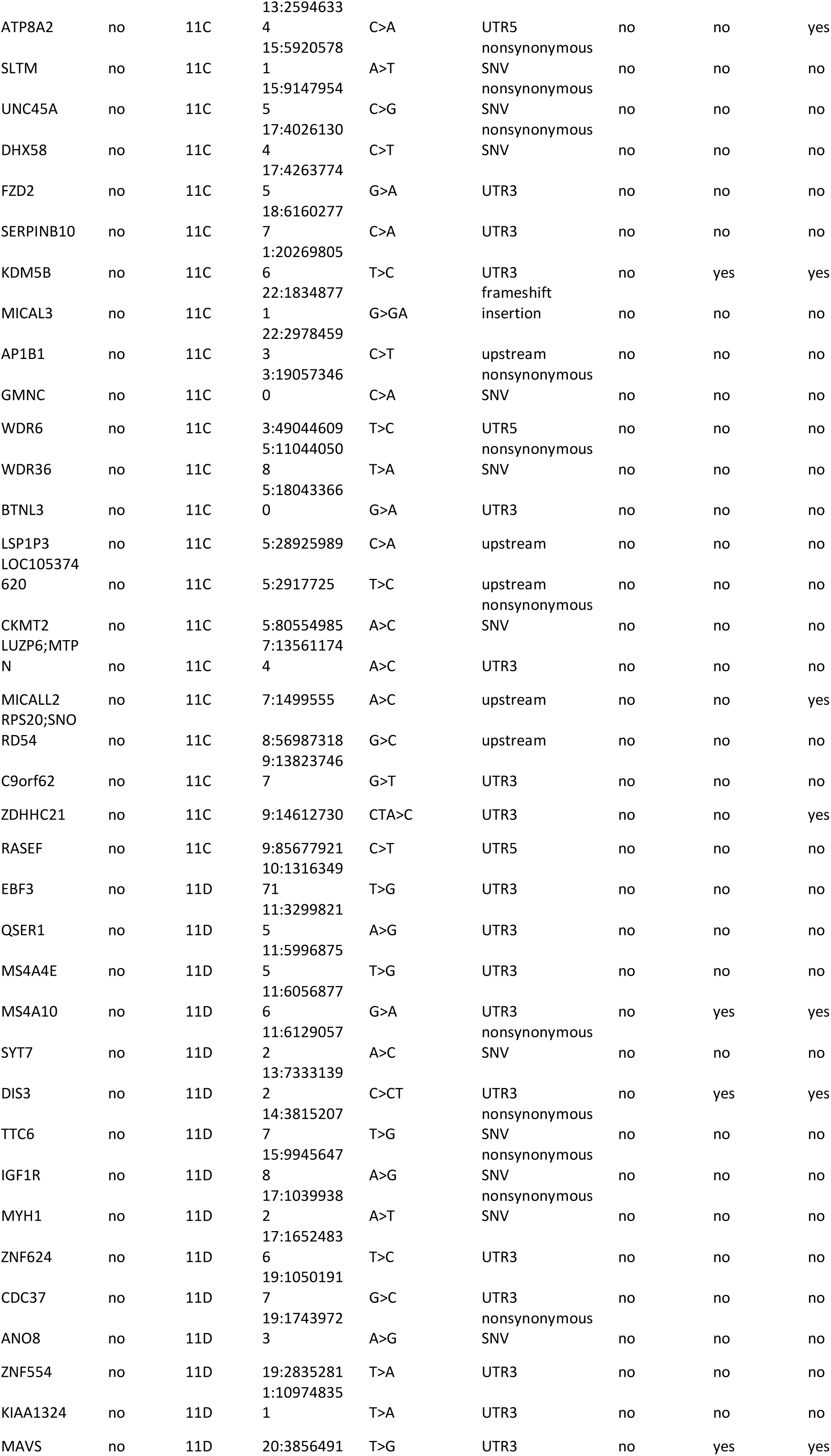
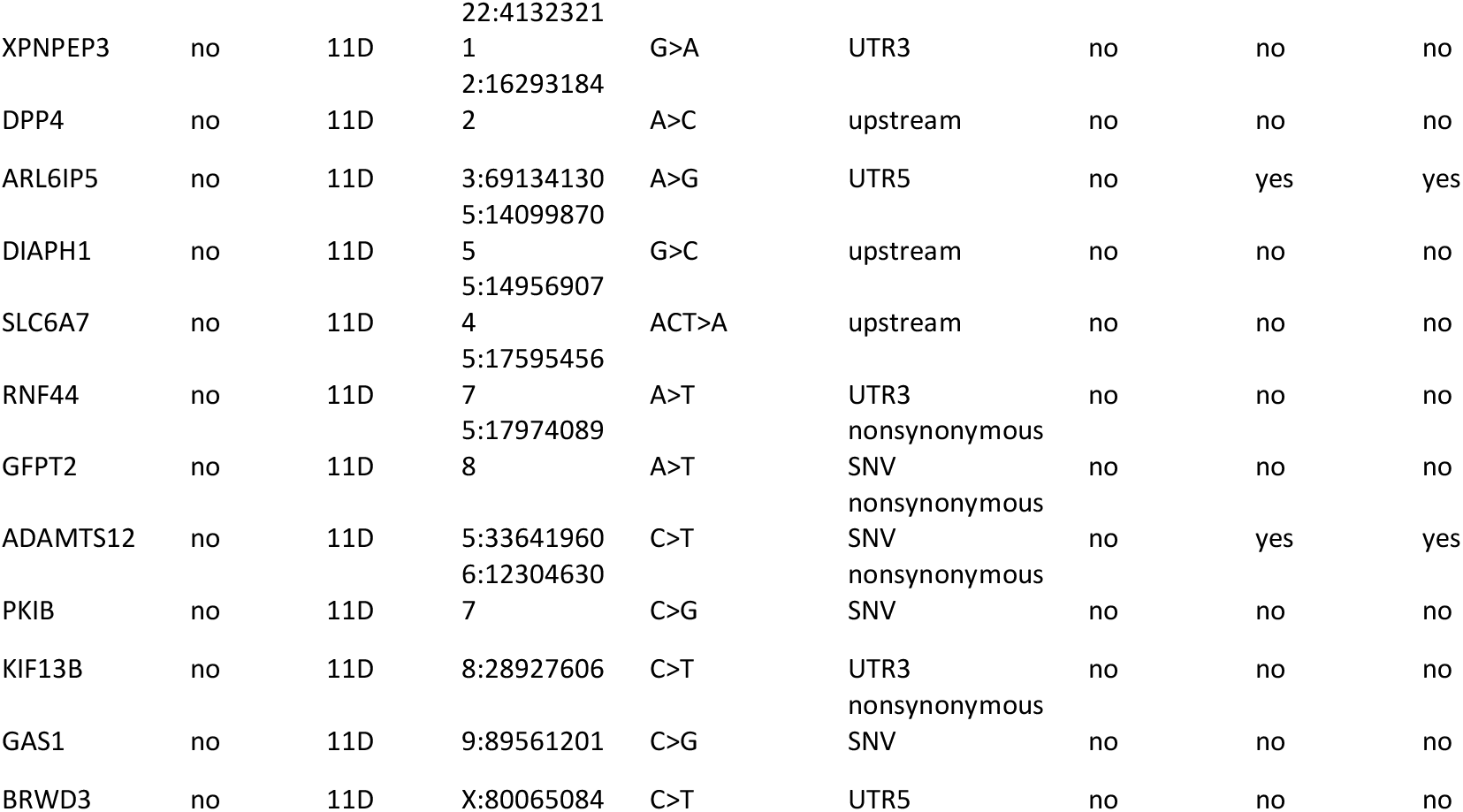
Somatic non-synonymous, splicing, upstream (promoter) or UTR mutations by gene based on calls filtered for dbSNP138 population variants.

## Notes

### Competing Interest Statement

The authors have declared no competing interest.

## References

Alexandrov, L.B., Kim, J., Haradhvala, N.J., Huang, M.N., Tian Ng, A.W., Wu, Y., Boot, A., Covington, K.R., Gordenin, D.A., Bergstrom, E.N., et al. (2020). The repertoire of mutational signatures in human cancer. Nature 578, 94–101.

Andersson, E., Sward, C., Stenman, G., Ahlman, H., and Nilsson, O. (2009). High-resolution genomic profiling reveals gain of chromosome 14 as a predictor of poor outcome in ileal carcinoids. Endocr Relat Cancer 16, 953–966.

Banck, M.S., Kanwar, R., Kulkarni, A.A., Boora, G.K., Metge, F., Kipp, B.R., Zhang, L., Thorland, E.C., Minn, K.T., Tentu, R., et al. (2013). The genomic landscape of small intestine neuroendocrine tumors. J Clin Invest 123, 2502–2508.

Bellono, N.W., Bayrer, J.R., Leitch, D.B., Castro, J., Zhang, C., O’Donnell, T.A., Brierley, S.M., Ingraham, H.A., and Julius, D. (2017). Enterochromaffin Cells Are Gut Chemosensors that Couple to Sensory Neural Pathways. Cell 170, 185–198

Chen, X., Schulz-Trieglaff, O., Shaw, R., Barnes, B., Schlesinger, F., Kallberg, M., Cox, A.J., Kruglyak, S., and Saunders, C.T. (2016). Manta: rapid detection of structural variants and indels for germline and cancer sequencing applications. Bioinformatics 32, 1220–1222.

Choi, A.B., Maxwell, J.E., Keck, K.J., Bellizzi, A.J., Dillon, J.S., O’Dorisio, T.M., and Howe, J.R. (2017). Is Multifocality an Indicator of Aggressive Behavior in Small Bowel Neuroendocrine Tumors? Pancreas 46, 1115–1120.

Crona, J., Gustavsson, T., Norlen, O., Edfeldt, K., Akerstrom, T., Westin, G., Hellman, P., Bjorklund, P., and Stalberg, P. (2015). Somatic Mutations and Genetic Heterogeneity at the CDKN1B Locus in Small Intestinal Neuroendocrine Tumors. Ann Surg Oncol 22 *Suppl 3*, S1428–1435.

Curtius, K., Wright, N.A., and Graham, T.A. (2018). An evolutionary perspective on field cancerization. Nat Rev Cancer 18, 19–32.

Dasari, A., Shen, C., Halperin, D., Zhao, B., Zhou, S., Xu, Y., Shih, T., and Yao, J.C. (2017). Trends in the Incidence, Prevalence, and Survival Outcomes in Patients With Neuroendocrine Tumors in the United States. JAMA Oncol 3, 1335–1342.

Dumanski, J.P., Rasi, C., Bjorklund, P., Davies, H., Ali, A.S., Gronberg, M., Welin, S., Sorbye, H., Gronbaek, H., Cunningham, J.L., et al. (2017). A MUTYH germline mutation is associated with small intestinal neuroendocrine tumors. Endocr Relat Cancer 24, 427–443.

Francis, J.M., Kiezun, A., Ramos, A.H., Serra, S., Pedamallu, C.S., Qian, Z.R., Banck, M.S., Kanwar, R., Kulkarni, A.A., Karpathakis, A., et al. (2013). Somatic mutation of CDKN1B in small intestine neuroendocrine tumors. Nat Genet 45, 1483–1486.

Gangi, A., Siegel, E., Barmparas, G., Lo, S., Jamil, L.H., Hendifar, A., Nissen, N.N., Wolin, E.M., and Amersi, F. (2018). Multifocality in Small Bowel Neuroendocrine Tumors. J Gastrointest Surg 22, 303–309.

Geoffroy, V., Herenger, Y., Kress, A., Stoetzel, C., Piton, A., Dollfus, H., and Muller, J. (2018). AnnotSV: an integrated tool for structural variations annotation. Bioinformatics 34, 3572–3574.

Graham, T.A., McDonald, S.A., and Wright, N.A. (2011). Field cancerization in the GI tract. Future Oncol 7, 981–993.

Guo, Z., Li, Q., Wilander, E., and Ponten, J. (2000). Clonality analysis of multifocal carcinoid tumours of the small intestine by X-chromosome inactivation analysis. J Pathol 190, 76–79.

Horn, S., Figl, A., Rachakonda, P.S., Fischer, C., Sucker, A., Gast, A., Kadel, S., Moll, I., Nagore, E., Hemminki, K., et al. (2013). TERT promoter mutations in familial and sporadic melanoma. Science 339, 959–961.

Huang, F.W., Hodis, E., Xu, M.J., Kryukov, G.V., Chin, L., and Garraway, L.A. (2013). Highly recurrent TERT promoter mutations in human melanoma. Science 339, 957–959.

Katona, T.M., Jones, T.D., Wang, M., Abdul-Karim, F.W., Cummings, O.W., and Cheng, L. (2006). Molecular evidence for independent origin of multifocal neuroendocrine tumors of the enteropancreatic axis. Cancer Res 66, 4936–4942.

Korbel, J.O., and Campbell, P.J. (2013). Criteria for inference of chromothripsis in cancer genomes. Cell 152, 1226–1236.

Krzywinski, M., Schein, J., Birol, I., Connors, J., Gascoyne, R., Horsman, D., Jones, S.J., and Marra, M.A. (2009). Circos: an information aesthetic for comparative genomics. Genome Res 19, 1639–1645.

Lawrence, M.S., Stojanov, P., Polak, P., Kryukov, G.V., Cibulskis, K., Sivachenko, A., Carter, S.L., Stewart, C., Mermel, C.H., Roberts, S.A., et al. (2013). Mutational heterogeneity in cancer and the search for new cancer-associated genes. Nature 499, 214–218.

Lodato, M.A., Rodin, R.E., Bohrson, C.L., Coulter, M.E., Barton, A.R., Kwon, M., Sherman, M.A., Vitzthum, C.M., Luquette, L.J., Yandava, C.N., et al. (2018). Aging and neurodegeneration are associated with increased mutations in single human neurons. Science 359, 555–559.

Lodato, M.A., Woodworth, M.B., Lee, S., Evrony, G.D., Mehta, B.K., Karger, A., Lee, S., Chittenden, T.W., D’Gama, A.M., Cai, X., et al. (2015). Somatic mutation in single human neurons tracks developmental and transcriptional history. Science 350, 94–98.

Magi, A., Pippucci, T., and Sidore, C. (2017). XCAVATOR: accurate detection and genotyping of copy number variants from second and third generation whole-genome sequencing experiments. BMC Genomics 18, 747.

Priestley, P., Baber, J., Lolkema, M.P., Steeghs, N., de Bruijn, E., Shale, C., Duyvesteyn, K., Haidari, S., van Hoeck, A., Onstenk, W., et al. (2019). Pan-cancer whole-genome analyses of metastatic solid tumours. Nature 575, 210–216.

Rosenthal, R., McGranahan, N., Herrero, J., Taylor, B.S., and Swanton, C. (2016). DeconstructSigs: delineating mutational processes in single tumors distinguishes DNA repair deficiencies and patterns of carcinoma evolution. Genome Biol 17, 31.

Sei, Y., Feng, J., Zhao, X., Forbes, J., Tang, D., Nagashima, K., Hanson, J., Quezado, M.M., Hughes, M.S., and Wank, S.A. (2016). Polyclonal Crypt Genesis and Development of Familial Small Intestinal Neuroendocrine Tumors. Gastroenterology 151, 140–151.

Sei, Y., Zhao, X., Forbes, J., Szymczak, S., Li, Q., Trivedi, A., Voellinger, M., Joy, G., Feng, J., Whatley, M., et al. (2015). A Hereditary Form of Small Intestinal Carcinoid Associated With a Germline Mutation in Inositol Polyphosphate Multikinase. Gastroenterology 149, 67–78.

Stecher, G., Tamura, K., and Kumar, S. (2020). Molecular Evolutionary Genetics Analysis (MEGA) for macOS. Mol Biol Evol 37, 1237–1239.

Tate, J.G., Bamford, S., Jubb, H.C., Sondka, Z., Beare, D.M., Bindal, N., Boutselakis, H., Cole, C.G., Creatore, C., Dawson, E., et al. (2019). COSMIC: the Catalogue Of Somatic Mutations In Cancer. Nucleic Acids Res 47, D941–D947.

Walsh, K.M., Choi, M., Oberg, K., Kulke, M.H., Yao, J.C., Wu, C., Jurkiewicz, M., Hsu, L.I., Hooshmand, S.M., Hassan, M., et al. (2011). A pilot genome-wide association study shows genomic variants enriched in the non-tumor cells of patients with well-differentiated neuroendocrine tumors of the ileum. Endocr Relat Cancer 18, 171–180.

Walter, D., Harter, P.N., Battke, F., Winkelmann, R., Schneider, M., Holzer, K., Koch, C., Bojunga, J., Zeuzem, S., Hansmann, M.L., et al. (2018). Genetic heterogeneity of primary lesion and metastasis in small intestine neuroendocrine tumors. Sci Rep 8, 3811.

Wang, Y.Z., Carrasquillo, J.P., McCord, E., Vidrine, R., Lobo, M.L., Zamin, S.A., Boudreaux, P., and Woltering, E. (2014). Reappraisal of lymphatic mapping for midgut neuroendocrine patients undergoing cytoreductive surgery. Surgery 156, 1498–1502; discussion 1502-1493.

Yates, L.R., and Campbell, P.J. (2012). Evolution of the cancer genome. Nat Rev Genet 13, 795–806.

Zhang, Z., Makinen, N., Kasai, Y., Kim, G.E., Diosdado, B., Nakakura, E., and Meyerson, M. (2020). Patterns of chromosome 18 loss of heterozygosity in multifocal ileal neuroendocrine tumors. Genes Chromosomes Cancer.

## References

Larson, N.B., and Fridley, B.L. (2013). PurBayes: estimating tumor cellularity and subclonality in next-generation sequencing data. Bioinformatics 29, 1888–1889.

McLaren, W., Pritchard, B., Rios, D., Chen, Y., Flicek, P., and Cunningham, F. (2010). Deriving the consequences of genomic variants with the Ensembl API and SNP Effect Predictor. Bioinformatics 26, 2069–2070.

Nersisyan, L., and Arakelyan, A. (2015). Computel: computation of mean telomere length from whole-genome next-generation sequencing data. PLoS One 10, e0125201.

Tang, K.W., Alaei-Mahabadi, B., Samuelsson, T., Lindh, M., and Larsson, E. (2013). The landscape of viral expression and host gene fusion and adaptation in human cancer. Nat Commun 4, 2513.

Tubio, J.M.C., Li, Y., Ju, Y.S., Martincorena, I., Cooke, S.L., Tojo, M., Gundem, G., Pipinikas, C.P., Zamora, J., Raine, K., et al. (2014). Mobile DNA in cancer. Extensive transduction of nonrepetitive DNA mediated by L1 retrotransposition in cancer genomes. Science 345, 1251343.

Wang, K., Li, M., and Hakonarson, H. (2010). ANNOVAR: functional annotation of genetic variants from high-throughput sequencing data. Nucleic Acids Res 38, e164.

